# In vitro spatiotemporal reconstruction of human skeletal muscle organogenesis

**DOI:** 10.1101/2024.05.10.593520

**Authors:** Lampros Mavrommatis, Nassam Daya, Leon Volke, I-Na Lu, Heidi Zhuge, Martin Stehling, Dagmar Zeuschner, Hyun-Woo Jeong, Ji-Hun Yang, Gerd Meyer zu Hörste, Beate Brand-Saberi, Hans R. Schöler, Matthias Vorgerd, Holm Zaehres

## Abstract

Spatiotemporal recapitulation of long-range trajectories for lineages that influence body patterning along the medio-lateral and proximal-distal axes during embryogenesis in an *in vitro* system remains elusive. Here we introduce a three-dimensional organoid approach, termed Gastruloids-Lateraloid-Musculoids (GLMs), to model human neural crest, lateral plate mesoderm and skeletal muscle lineage development at the forelimb level following gastrulation and during limb patterning. GLMs harvest neuro-mesodermal progenitors with the potential to establish neural and paraxial mesodermal populations, while single cell analyses and spatial transcriptomics demonstrate promotion of mesodermal lineage segregation during gastrulation and spatial recapitulation of migration events along the medio-lateral axis for vagal neural crest, hypaxial myogenesis and lateral plate mesodermal lineages. Comparative analyses to developmental atlases and adult muscle stem cell data confirm a pool of hypaxial migrating myogenic progenitors that in a niche dependent manner change their embryonic anatomical developmental program to a fetal myogenic program, thus enabling them to resist specification in a cell autonomous manner and facilitate long term *in vitro* expansion. GLMs model human myogenesis at the forelimb level, establish fetal muscle stem cells equivalent to those that sustain the growth phase of the embryo and provide a 3D *in vitro* system for investigating neural crest, early fore-gut and lateral plate mesoderm development.

## Introduction

Mammalian body axis patterning, specification and elongation are initiated during gastrulation, whereby forebrain emerges at the anterior portion of neural plate, while more posteriorly and closer to primitive streak transient embryonic structures and signaling centers, indispensable for successful neural tube, lateral plate mesoderm and early skeletal muscle organogenesis, are developed during body axis elongation. The original Nieuwkoop’s “activation-transformation” hypothesis on neural induction and central nervous system development in amphibian embryos suggested an initial anterior neural plate induction that is followed by the formation of caudal neural regions via patterning of this anterior tissue with posteriorising signals^1,2^. Upcoming studies on neural induction during organogenesis updated this model by further suggesting neuro-mesodermal progenitors (axial stem cells, NMPs), arising from primitive streak-associated caudal lateral epiblast with a subsequent ongoing decision between neural and mesodermal fates, as a key population influencing body axis elongation^3,4,5,6,7,8^. Recently, single-cell transcriptomic characterization of a gastrulating human embryo further indicated overlapping expression of markers of established mesodermal sub-types, such as paraxial or lateral plate mesoderm^9^, thus suggesting presence of transitional mesodermal states during initial body axis patterning. Breaking medio-lateral symmetry at forelimb level following gastrulation is associated with lateral plate mesoderm formation in a BMP dependent manner, reviewed by Prummel et al.^10^, while limb bud initiation at forelimb level is modulated by the adjacent somitic mesoderm, that in the absence of retinoic acid signalling limb bud initiation is impaired^11,12, 13,14,15,16^.

Establishing 3D systems that simulate gastrulation *in vitro*^17,18,19,20,21,22,23,24,25^, and derive neuro-mesodermal progenitors from human and mouse hiPSCs^26,27,28,29,30^ offered the possibility to investigate gene regulatory networks and signaling pathways that govern NMPs identity and influence body axes pattering^31^, while subsequent studies harness their ability to self-organize and generate either neuromuscular organoids^32^ or via matrigel support to simulate body axis elongation for paraxial mesodermal lineage^33,34,35^, or in co-development with neural lineage in mouse and human models^21,36^. Axially patterned embryonic organoids capture cardiogenesis *in vitro*^37,38,39^ together with fore-gut development in humans^40^, while recent two dimensional studies with hiPSCs demonstrate a neuro-mesodermal mediated neural crest origin^41,42^. To our knowledge, current anterior-posterior patterned stem cell developmental models are unable to spatially recapitulate developmental trajectories along the medio-lateral or proximal distal axis. Here to address this limitation we present a method of immediate matrix support and concomitant growth factor applications on hiPSCs aggregates to model hypaxial myogenesis, limb mesenchyme and neural crest migration via an intermediate neuro-mesodermal progenitor stage and to mimic patterning at forelimb level along the major body axes during embryogenesis. GLMs (Gastruloids/Lateraloids/Musculoids) support and as a developmental model uniquely recapitulate skeletal muscle trajectory at all stages of human fetal forelimb development. GLM-derived myogenic progenitors and biopsy derived adult muscle stem cells were compared to human skeletal muscle reference atlases and present the *in vitro* generation of fetal muscle stem cells that exhibit properties of long-term *in vitro* self-renewal and specification resistance, a key characteristic that defines *in vivo* fetal and adult muscle stem cells^43,44^.

## Results

### Gastrulation and mesodermal segregation at early stages

From a pluripotent embryonic body state, human PSC-derived aggregates expressing the pluripotency markers, e.g NANOG, at day of Matrigel embedding **(Figure 1A, Day 0)**, underwent gastrulation following stimulation with Wnt activation (CHIR99021), BMP inhibition, (LDN193189) and bFGF. Immunocytochemistry analysis at Day 2, indicates presence of epiblast NANOG^+^, SOX2^+^ populations at the core of GLM, surrounded by cells expressing the early gastrulation marker Goosecoid (GSC), while more ventrally and at a different to NANOG^+^, SOX2^+^ section planes, presence of TBXT, GATA6, and SOX17 populations, an indication of ongoing gastrulation and of mesodermal, primitive endodermal fate **(Figure 1A,H, Day2)**. FOXA2 expression at the core of the structure further indicates a definitive endodermal/axial mesodermal fate, and of mesodermal and endodermal populations at the periphery of the structure via presence of GATA6^+^ and GATA6^+^/FOXA2^+^ populations, respectively **(Figure 1H, Day2)**. This conformation along the radial axis reassembles the micro-patterned induced differentiation of human embryonic stem cells (hESCs) to capture gastrulation-like events in two dimensions^45^. Our system further, by supporting development in three-dimensions, preserves a caudal epiblast state, CDX2^+^, on a different level from the mesodermal and endodermal state and generates a dorso-ventral like axis that differentiates their development at following stages **(Figure 1A,B,D, asterisks, 2A,F)**. At gastrulation stage, single cell expression profiling and trajectory analysis indicates an epiblast stage, governed by HOX gene regulatory networks (calculated using SCENIC pipeline^91^) but organoids do not exhibit any axial elongation at histological level, and from a mesodermal and endodermal state **(Figure 1E,F, Figure S1A,B)**. Investigation on the mesodermal differential potential and cell fate probabilities at Day 5 using Palantir algorithm^95^ indicates that from a primitive streak state (*SOX2^+^,NKX1-2^+^,TBXT^+^)*, nascent mesodermal populations bifurcate to a paraxial, (*FOXC1^+^,TCF15^+^,MEOX1^+^)* and to a lateral plate mesodermal state, (*FOXF1^+^,GATA6^+^,HAND2^+^).* Pseudotime and RNA velocity analysis during mesodermal lineage segregation indicates BMP4 and RALDH2 upregulation for lateral and paraxial mesodermal fate respectively, while using Palantir algorithm we describe that each state excludes bipotent cell fate probability, thus lineage specification occurs at this stage **(Figure 1G, Figure S1A,B**). Mapping of single cell expression profiling derived from human gastrulating embryo^9^ and GLMs at Day 7 to spatial transcriptomic data from Day5 GLMs via CytoSPACE pipeline^90^, demonstrates similar mapping between CS7 embryo and Day7 GLMs and highlights a section plane that simulates gastrulation, with presence of endodermal SOX17, FOXA2 populations and an emergent mesodermal identity, that further segregates to lateral plate, GATA6^+^, and paraxial/pre-somitic populations, TBX6^+^ **(Figure 1I, Figure S1E-G,S2A,B)**. Applying a Semi-supervised (SCANVI) deep learning model^99,100^ to human gastrulating embryo highlights similar dynamics during human primitive streak mesoderm development, as well as predicts presence of nascent, emergent and advanced mesoderm^9^, while 3D modelling of the CS8 human gastrulating embryo^46^ further depicts paraxial, lateral plate and intermediate mesodermal segregation at this stage **(Figure S1C,D,G).** Investigation at the histological level depicts a core TBXT^+^ gastrulating mesodermal population in the absence of SOX2 expression **(Figure 1A, Day2).** Furthermore, spatial transcriptomics analysis at this stage and immunocytochemistry pictures indicates an initial dorsoventral development during gastrulation stage, with an epiblast/ectodermal state, expressing *NANOG, CDX2, SOX2*, at its dorsal portion and an endodermal, mesodermal state ventrally **(Figure 1A,D,H)**. Interestingly, adapting conditions to our system that are present at tail bud stage and during axial elongation, such as prolonged Wnt activation (CHIR99021), BMP inhibition, (LDN193189), and FGF stimulation, induces at its dorsal caudal embryo-like epiblast portion neuro-mesodermal progenitor populations, expressing, SOX2 and TBXT, and simulates body axis elongation **(Figure 1B,C, Figure S2C-E).** This geometry influences GLMs spatiotemporal development and is responsible for the two independent trajectories present during paraxial mesoderm formation within GLMs. At its dorsal exterior, the neural epithelium at specific site adopts neuro-mesodermal fate and enters paraxial mesoderm and neural crest development via an intermediate neuro-mesodermal progenitor (NMP stage), while at its ventral interior following gastrulation it channels its differentiation via emerging mesodermal identity towards paraxial mesoderm and lateral plate limb bud development **(Figure 1A,B,D,I, Figure S3D,E).**

**Figure 1.**
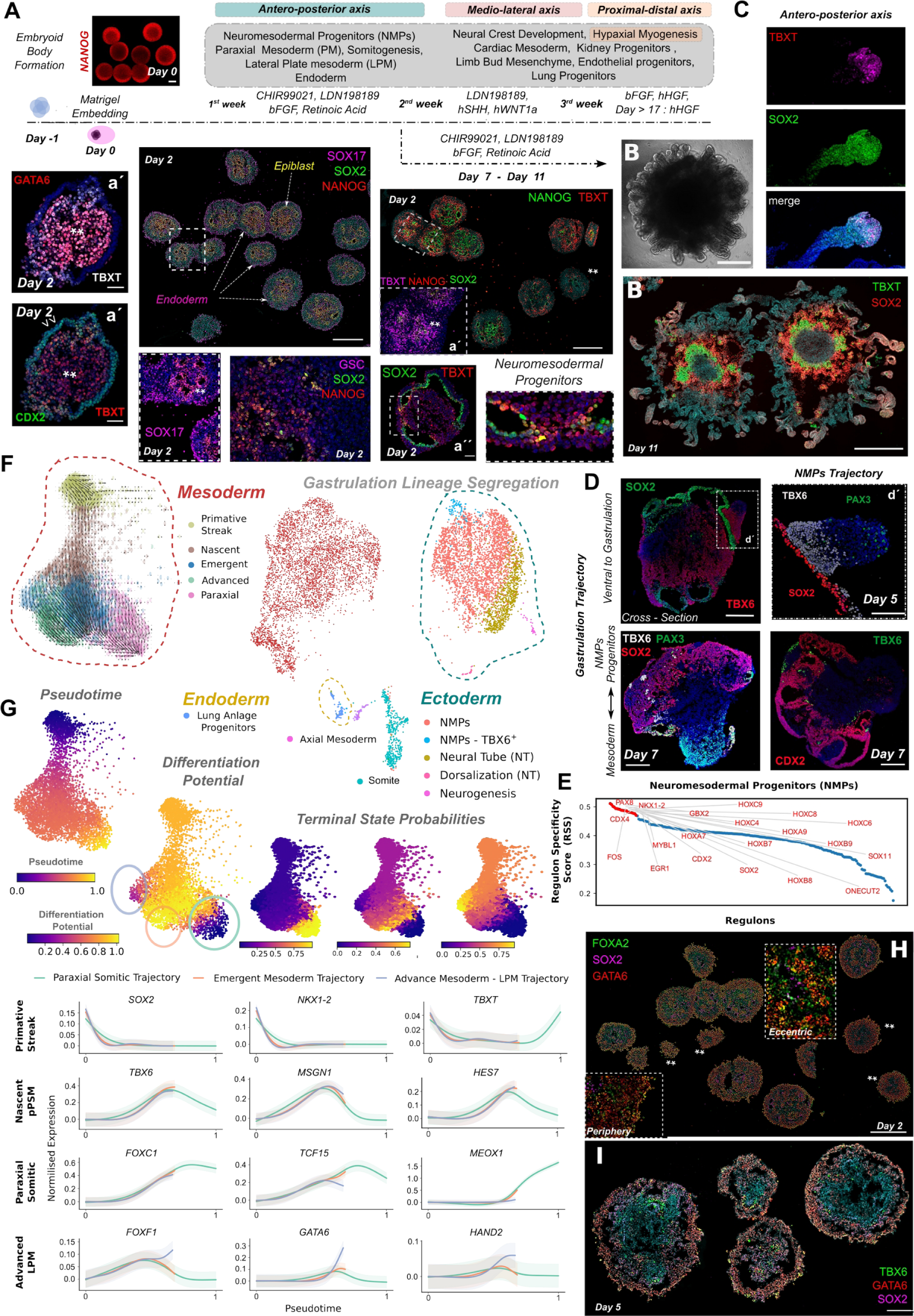
Gastrulation and mesodermal lineage segregation at early stages of GLM development. **(A)** Scheme depicting cytokine composition and GLM spatiotemporal development of the corresponding lineages along the major body axes. Bright-field and immunocytochemistry images illustrating histological key stages of GLM development simulating gastrulation in a dorsoventral plane. Dorsal/central portion is composed of NANOG^+^/SOX2^+^ epiblast populations, that ventrally undergo gastrulation, GSC^+^/TBXT^+^ populations, and generate SOX17 endodermal populations. Scale Bar: 500um, 50uM in a′**(B)** Following gastrulation at Day7 continuation of the CHIR99021, LDN198189, bFGF and Retinoc acid stimulation leads to antero-posterior axis formation at GLMs dorsal part, via expansion of neuro-mesodermal progenitor (NMPs) derived at Day2. The core consists of epiblast, gastrulating populations, and **(C)** the surface from NMPs (SOX2^+^/TBXT^+^) simulating tail bud and body axis elongation. **(D)** Immunocytochemistry pictures illustrating somitogenesis at dorsal location via NMP trajectory and at ventral location following gastrulation and mesoderm segregation. Scale Bar: 200um,100uM in d′ **(E)** Regulon score specificity (RSS) at neural cluster highlights HOX GRNs upregulation together with NMP specific NKX1-2 Regulon. **(F)** Single cell expression profiling (UMAP) at Day 7 indicates ongoing gastrulation and derivation of mesodermal, ectodermal and endodermal populations. RNA velocity analysis on force directed graph embedding indicates from a primitive streak state mesodermal lineage segregation towards lateral plate and paraxial mesodermal trajectories. **(G)** Examining differentiation potential and cell fate probabilities during mesodermal segregation using Palantir algorithm. Gene expression trends for primitive streak, *NKX1-2*, *TBXT,* nascent pre-somitic mesoderm (pPSM), *TBX6, HES7, MSGN1,* somitic, *FOXC1, TCF15, MEOX1* and LPM, *FOXF1, GATA6, HAND2,* markers. Trends are colored based on lineages presented in Fig.1d, Shaded region represents 1 s.d. **(H)** Immunocytochemistry picture at Day 2 at gastrulation plane during GLM development depicts at the core progenitors with axial mesodermal identity FOXA2^+^/GATA6^−^, surrounded at the periphery from mesodermal, FOXA2^+^/GATA6^−^, and endodermal, FOXA2^+^/GATA6^+^ populations. Scale Bar: 500um. **(I)** Immunocytochemistry picture at Day 5 at level ventral to gastrulation plane during GLM development depicts bipotent with emerging mesodermal identity progenitors, TBX6^+^/GATA6^+^, together with progenitors of lateral plate mesoderm, GATA6^+^, and of paraxial mesoderm, TBX6^+^, origin. Scale Bar: 200um.

### Lineage spatiotemporal development following gastrulation - Lateraloids

At the early stages of GLM development, stimulating PSC-derived aggregates with CHIR99021, BMP inhibitor LDN193189, bFGF and retinoic acid promotes gastrulation without patterning the residing CDX2^+^/GBX2^+^ caudal epiblast neural populations **(Figure 1A,E)**. To promote dermomyotomal hypaxial fate on paraxial mesodermal populations, in a first step we simulated developmental cues secreted from dorsal neural tube (WNT1A) and notochord (hSHH) during embryogenesis while maintaining constant BMP inhibition to avoid excessive lateral plate mesoderm formation, followed by a second myogenic induction step that included FGF, HGF stimulation **(Figure 1A).** Force-directed *k*-nearest neighbor graph and RNA velocity demonstrated spatiotemporal development and presence of mesodermal, endodermal and ectodermal developmental trajectories following gastrulation **(Figure 2A).** At this stage WNT1, an agent influencing neural crest development^47^, drives differentiation of the caudal epiblast, located dorsally at GLMs, towards dorsal neural tube and neural crest development **(Figure 2A, Figure S3C).** Neural crest induction was further verified by the presence of TFAP2A^+^ populations that marked the neural plate border formation at PAX6^+^ dorsal neural tube regions and preceded the generation of a SOX10^+^ neural crest migrating stream **(Figure 2H).** During migration, neural crest follows cells fate decisions that lead to sensory - autonomic bifurcation when mapped to mouse trunk 9.5 neural crest^107^ **(Figure S3C).** Similar to the gastrulation stage, we noticed that in comparison to 2D differentiation approaches^48^ BMP inhibitor LDN193189 (0.5nM) in GLMs did not inhibit lateral plate mesoderm development. Consequently, ventrally, lateral plate mesoderm during the second week of GLM development differentiates towards cardiac field, expressing *HAND1, HAND2, MEIS2, GATA3, GATA4,* and limb bud initiation, expressing *FGF10, SHOX2, PRRX2, PRRX1, HGF* trajectories **(Figure 2A, Figure S3A)**, while after the second week only towards limb bud mesenchymal trajectory, *PRRX1, PRRX2, TWIST1, TWIST2,* that undergoes maturation and expresses osteogenic markers, *OGN, POSTN,* as well as establishes fibro-adipogenic progenitors, *PDGFRA^+^, PDGFRB^+^*, able to enter adipogenesis **(Figure 2A, Figure S3A,4G-I)**. Applying the Semi-supervised (SCANVI) deep learning model to human embryo, at stages following gastrulation^110,111^, CS10–CS16, demonstrates medio-lateral and proximal-distal development for LPM (lateral plate mesoderm) within GLMs, as GLM derived limb bud exhibits forelimb axial anatomical identity, in comparison to GLM derived endothelial cells and cardiac LPM, that exhibit visceral and trunk anatomical identity respectively **(Figure 2K,L).**

**Figure 2.**
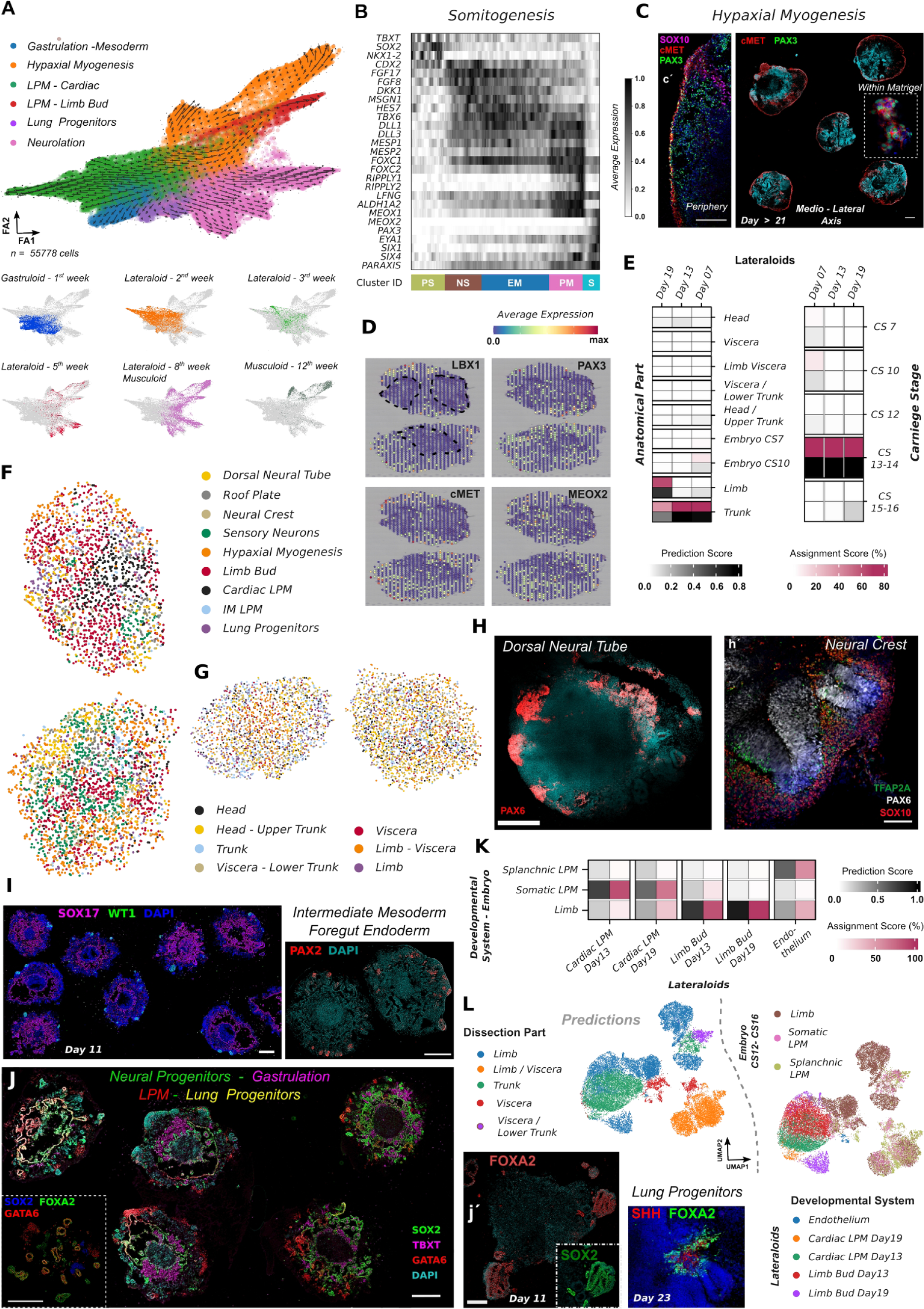
Lineage spatiotemporal development following gastrulation along the medio-lateral axis-Lateraloids. **(A)** Force-directed k-nearest neighbor graph on 55.778 cells at Day 7, Day 13, Day 19, Day 35, Day 56 and Day 84 and RNA velocity analysis indicating lineage progression following gastrulation along skeletal muscle, neural crest, dorsal neural tube, fore-gut endoderm and lateral plate mesoderm (cardiac field and limb bud mesenchyme) developmental trajectories. Feature plots highlight lineage representation at each stage. **(B)** Pseudotemporal ordering of cells related to somitogenesis clusters revealed gene dynamics from a tail bud /PS state, e.g. *NKX1-2, TBXT, FGF17, CDX2* that via posterior presomitic mesoderm (pPSM) e.g. *MSGN1, TBX6, HES7, MESP2*, and determination front formation,e.g. *RIPPLY2, LFNG, ALDH1A2,* promoted somitic mesoderm formation, e.g *PAX3, EYA1, SIX1, PARAXIS.* PS: Primitive streak, NS: Nascent mesoderm, EM: Emerging Mesoderm, PM:Paraxial Mesoderm, S: Somitic Mesoderm **(C)** Immunocytochemistry pictures on PAX3, SOX10, cMET markers at stages post Day 21 of GLM development, depict migration waves simulating hypaxial myogenesis along the medio-lateral axis for skeletal muscle lineage. Scale Bars: 500 um in c, 200 um in c′. **(D)** Spatial feature plots at Day 19, on *LBX1, PAX3, cMET and MEOX2* markers indicate migration waves distal to the initial GLM core structure (dashed lines) simulating hypaxial myogenesis migration. **(E)** Heatmap with percentage of certainty and assignment score from GLM cells along the hypaxial myogenesis trajectory from somite till migration stage upon unbiased mapping to the *in vivo* counterpart, Human Carnegie Stage (CS7-CS16) embryos. **(F)** Mapping of single cells from Day 19 GLMs and human CS12-CS16 developing embryo **(G)** to spatial GLM sections. Top section is derived from ventral to gastrulation plane. Bottom section from dorsal to gastrulation plane. **(H)** Immunocytochemistry pictures at 3^rd^ week of GLM development depicts a section plane that includes a section plane of dorsal neural tube and neural crest development/migration (right). Scale Bars: 500uM, 100uM in **h′ (I)** Immunocytochemistry pictures at Day 11 of GLM development depicts a section plane lower to gastrulation that includes SOX17^+^ fore-gut endodermal populations, surrounded from cells, WT1^+^/PAX2^+^, with intermediate mesodermal identity. Scale Bars: 500uM **(J)** Immunocytochemistry picture at Day 11 depicts organoids at section planes that combine gastrulating/mesodermal (TBXT^+^), neural (SOX2^+^), LPM (GATA6^+^) and fore-gut endodermal populations, pulmonary identity (GATA6^+^ /SOX2^+^/FOXA2^+^). Scale Bars: 500uM. **(K)** Heatmap with percentage of prediction and assignment score from GLM cells along the lateral plate mesoderm (LPM) development upon unbiased mapping to the *in vivo* counterpart, splachic, somatic and limb LPM derived from Human embryos between CS12-CS16, Carnegie Stages. **(L)** UMAP plots based on semi-supervised deep learning approach (SCANVI) to map GLM derived LPM at Day13 and Day19, on human CS12-CS16 LPM fetal reference atlas predicts somatic LPM derivation within GLM with trunk to limb axial anatomical identity.

Endodermal trajectory is directed towards anterior fore-gut endoderm development by establishing SOX17, SOX2, KRT8, GATA6, SHH and FOXA2 populations, an expression profiling that corresponds to early pulmonary endodermal specification^49,50^ **(Figure 2A,I,J, Figure S3A,F).** Surrounding the ŚOX17^+^ endodermal, and in the proximity of GATA6^+^ lateral plate mesodermal populations, between the 2^nd^ to 3^rd^ week, we could detect tubular structures at the periphery of GLM ventral development, expressing PAX2, PAX8, OSR1, OSR2, WT1 markers, a clear indication of intermediate mesodermal identity **(Figure 2A,I, Figure S3A,B,F)**. Paraxial mesodermal trajectory undergone somitogenesis, where from a primitive steak stage and through nascent and emergent mesoderm formation, directed its differentiation towards posterior presomitic mesoderm (pPSM) that via a determination front (DF) state, *LFNG, RIPPLY2,* somitic mesoderm (S) emerged **(Figure 2B).** Culture conditions directed the paraxial mesodermal trajectory towards hypaxial myogenesis and the generation of MET^+^/PAX3^+/^SOX10^−^ migration stream lateral to the gastrulating GLM structure **(Figure 2C)**. Spatial transcriptomics and single cell expression profiling highlights hypaxial myogenesis distal to GLM gastrulating structures and an expression profiling of PAX3^+^ /cMET^+^ myogenic progenitors that co-express LBX1, PITX2, CXCR4, Ephrin A5 and FGFR1 receptors, thereby indicating susceptibility to potential limb bub mesenchyme guidance signals influencing hypaxial myogenesis^51^ **(Figure 2C,D, 3D)**. Integration comparison to developing human embryos following gastrulation, stages CS12–CS16, demonstrates that GLM derived paraxial mesodermal populations simulate a trunk to limb proximal-distal transition similar to CS7 gastrulating – CS12-CS16 developing embryos **(Figure 2E)**. Upon migration within the matrigel droplet, GLM radial / medio-lateral development does not correlate anymore to craniocaudal or ventro-dorsal axis, but rather simulates medio-lateral and proximal distal axis during embryogenesis. Mapping of single cell expression profiling derived from human CS12-CS16 embryo^111^ and GLMs at Day 19 to spatial transcriptomic data from Day19 GLMs, further demonstrates craniocaudal axis development on the GLMs dorsoventral axis, as its dorsal portion that harbors neural crest development simulates head-upper trunk development, while its ventral harboring LPM, paraxial and endoderm development simulates trunk, limb and viscera development **(Figure 2F,G, Figure S3F,G)**.

### Lateraloid to Musculoid transition signals embryonic to fetal myogenic transition

The benchmark and novelty of GLM as a culture system is its ability to harbor migration events following dorsal neural tube, paraxial and lateral plate mesoderm formation. At all stages post initial patterning, we attempted to favor skeletal muscle lineage through constant HGF stimulation that compensates the absence of lateral plate mesoderm in musculoids at intervals associated to stages that musculoid culture grew beyond the initial matrigel droplet, past 35-42 days of GLM development. Spatial transcriptomic on Day 35 demonstrates reproducibility between different musculoids, with presence of neural/neural crest and limb bud mesenchymal trajectories, while the skeletal muscle trajectory spatial transcriptomics and immunocytochemistry pictures indicate presence at the periphery of developing musculoids **(Figure 3B, Figure S4B)**. Embryonic myogenic progenitors, PAX3^+^/cMET^+^, in close proximity to GLM-derived trunk structures, exhibited migrating potential, presenting with uncommitted (MYOD1^−^) and up-regulated HOX genes cluster expression profiling related to forelimb bud identity **(Figure 3D,E, Figure S4A)**. Along this migration stream, embryonic PAX3 myogenic progenitors triggered the fetal program and co-expressed PAX7 **(Figure 3A, Figure S4A**). Spatial mapping to human fetal reference atlas indicates embryonic to fetal myogenic transition in a radial outwards direction, with fetal stages occurring at the edge of matrigel droplet **(Figure 3B, Figure S4C).** Differential expression and gene regulatory analysis between embryonic/fetal, PAX3^+^/PAX7^+^, and fetal, PAX7^+^, populations indicated that during the embryonic fetal transition myogenic progenitors ceased proliferation, prolonged their G1 phase and up-regulated expression of extracellular matrix (ECM) proteins and proteins associated to fetal program, myogenic maturation and satellite cell response, e.g. KLF4, FOXO3, NFIA, NFIX, EGR1, EGR2 and EGR3 **(Figure 3E, Figure S6B,C).** Likewise, PAX3+ myogenic progenitors during human and mouse embryonic fetal development under the influence of developing mesenchyme adapted strong anatomical identity, while during the embryonic fetal transition, fetal progenitors ceased proliferation, entered a prolonged G1 phase and up-regulated ECM expression **(Figure S4F,S5B,C,E,I,K,J)**. To quantitatively assess the anatomical identity (“anatomical score”) and myogenesis onset (“myogenesis score”) of myogenic progenitors during human, mouse and musculoid skeletal muscle development, we calculated an anatomical and myogenesis score, where we took the expression levels of HOX genes and markers related to fetal myogenesis in individual cells into account (Table S2, methods). Scatter and spatial transcriptomic plots from human fetal developing limbs and GLMs, demonstrate anatomical scores for embryonic myogenic progenitors and myogenesis scores for fetal/postnatal muscle stem cells, while during growth phase the anatomical score is downregulated from areas where skeletal muscle develops **(Figure 3C,D, Figure S4E,F,K,J, S5H, Table S2).**

**Figure 3.**
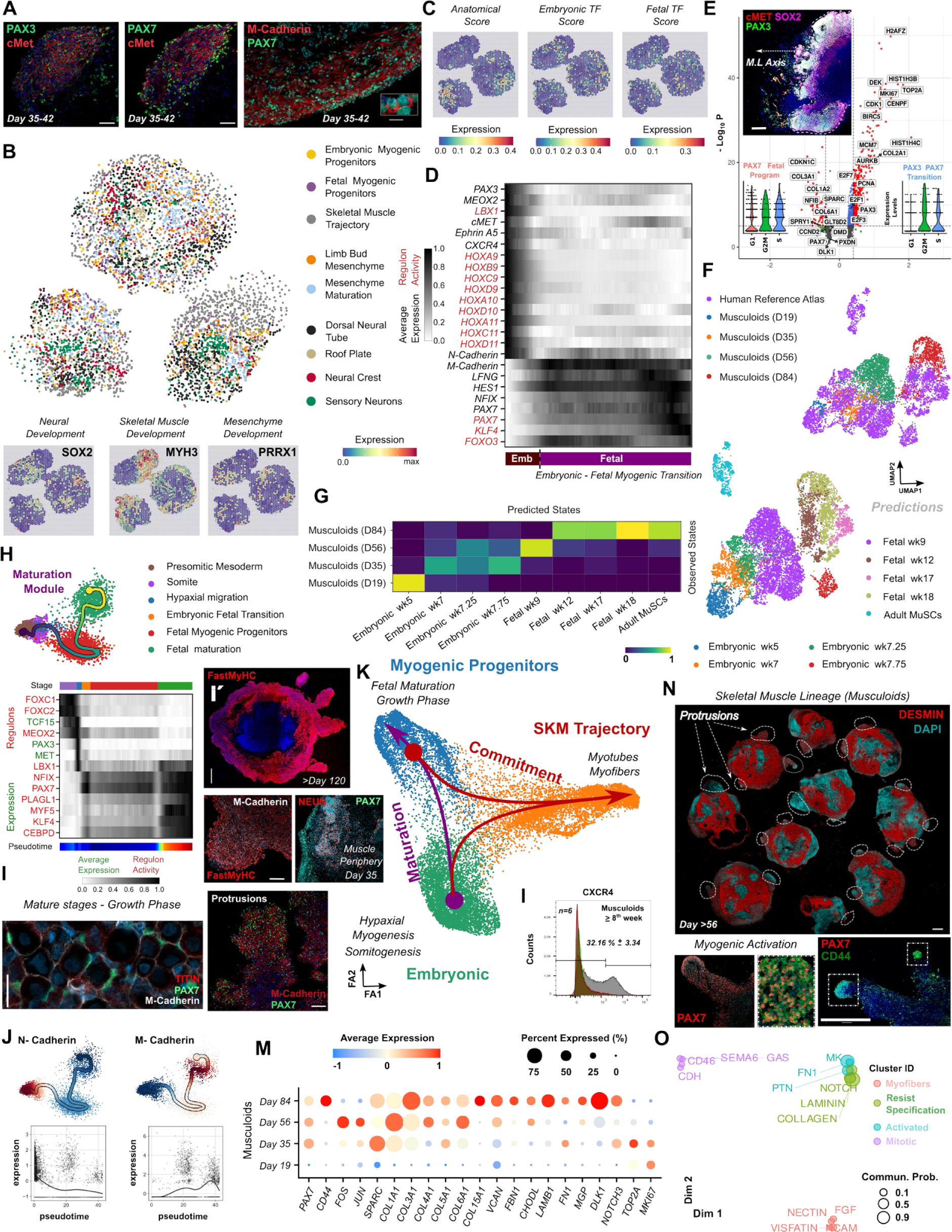
Lateraloid to Musculoid transition signals embryonic to fetal myogenic transition and recapitulation of skeletal muscle developmental trajectory. **(A)** Images at the edge of matrigel droplet depict embryonic (PAX3) to fetal myogenic transition (PAX7) and skeletal muscle system development. Scale Bar: 50um **(B)** Mapping of single cells from Day 35 GLM dataset to spatial GLM sections from the same stage indicates reproducibility on neural, skeletal muscle and mesenchymal clusters between different organoids. **(C)** Spatial feature plots depicting anatomical (HOX genes) and embryonic/fetal myogenic transcription factor scores based on selected genes **(D)** Pseudotemporal ordering of cells during embryonic, fetal myogenic transition reveals gene and GRN dynamics and highlight an axial/anatomical HOX gene upregulation for PAX3^+^/LBX1^+^ embryonic and absence on PAX7^+^/NFIX^+^ fetal myogenic progenitors. **(E)** Differential expression analysis between PAX7 and PAX3/PAX7 clusters highlights prolonged G1 phase, up-regulation of ECM proteins and cell cycle inhibitors in fetal myogenic progenitors. Violin plots depicting cell cycle stage on PAX7 and PAX3/PAX7 clusters. Picture depicts hypaxial migration process away from the GLM trunk-like core structure. Scale Bar: 200um **(F)** UMAP plots based on a semi-supervised deep learning (SCANVI) approach to map myogenic progenitors from GLM derived skeletal muscle developmental trajectory to the human skeletal muscle reference atlas demonstrates *in vitro* reconstruction till late fetal stages, maturation beyond the embryonic fetal transition stage for musculoid and *in vivo* matured PAX7 derived myogenic progenitors, while **(G)** Heatmap depicting the predicted and observed states for musculoid trajectory based on gene networks derived from the human reference myogenic map indicates developmental swift similar to human reference map and the presence of adult muscle stem cell gene networks within musculoids from Day 84. **(H)** Curved trajectory analysis on PCA space for musculoid skeletal muscle atlas. Pseudotime was calculated, by learning a principal curve on the 2 first PC components using the ElPiGraph algorithm. Pseudotemporal ordering of cells during musculoid skeletal muscle atlas reveals gene and GRN dynamics and highlight a continuous somitic e.g. FOXC1, FOXC2, TCF15, to hypaxial e.g. MEOX2, PAX3, LBX1 embryonic state that upon maturation opens fetal/postnatal developmental program e.g PAX7, NFIX, MYF5, KLF4. **(I)** Musculoid development at mature stages (>Day56) simulates muscle growth at the periphery of the organoid culture (FastMyHC^+^/PAX7^+/^NEUN^−^ sites). Protrusions, sites exceeding initial matrigel droplet limit, harbor the development of M-Cadherin positive myofibers and PAX7 positive myogenic progenitors at mature stages of musculoid development and muscle growth phase. Scale Bars: 100uM, 500um in i′ **(J)** Feature plots based on differential expression analysis between nodes along the musculoid derived curved trajectory, depicts M-Cadherin upregulation (skeletal muscle), N-Cadherin downregulation (mesenchyme) along pseudotime and indicates niche transition for myogenic progenitors. **(K)** Force-directed k-nearest neighbour graph indicating skeletal muscle trajectories present during musculoid development, one leads to myogenic progenitor maturation and one to myogenic commitment. **(L)** Histogram illustrating FACS quantification on CXCR4^+^ myogenic progenitors post 8^th^ week during musculoid development. Red histogram: unstained population, green histogram: isotype control, gray histogram: CXCR4^+^ population. **(M)** Dot plot illustrating expression of ECM-related genes and for genes related the activated, *CD44, JUN, FOS,* and a mitotic, *NOTCH3, TOP2A, MKI67,* state across the developmental trajectory on musculoid derived myogenic progenitors. The size of each circle reflects the percentage of cells in a cluster where the gene is detected, and the color reflects the average expression level within each cluster (blue, low expression; red, high expression). **(N)** DESMIN immunocytochemistry pictures at Day 56 indicate robust skeletal muscle development during GLM derivation. Dashed lines illustrate protrusions that harbor PAX7^+^CD44^+^ myogenic progenitor maturation. Scale Bar: 500um **(O)** Jointly projecting and clustering signaling pathways from 12^th^ week myogenic progenitors into a shared two-dimensional manifold according to their functional similarity. Each dot represents the communication network of one signaling pathway. Dot size is proportional to the total communication probability. Different colors represent different groups of signaling pathways.

Next, we examined whether musculoid-derived myogenic progenitors were able to surpass the embryonic fetal transition “barrier” stage^52^ and further maturate. Utilizing semi-supervised deep learning (SCANVI) model to map embryonic (Day 19) and 5^th^ week musculoid-derived myogenic progenitors into the human embryonic to fetal reference atlas, we demonstrate maturation beyond the embryonic fetal transition and generation of a pool of myogenic progenitors with fetal identity **(Figure S4D).** This deep learning model accurately predicts the PAX3/PAX7 transition by correlating musculoid derived PAX3^+^ progenitors to 5^th^ week, PAX3^+^/PAX7^+^ progenitors to 7^th^ week and PAX7^+^ progenitors to 9^th^ week of human fetal development **(Figure S4C,D).** Moreover, gene regulatory network analysis verified that during the 8^th^ week myogenic progenitors further up-regulated NFIX, KLF4, PAX7, MYF5 and entered the fetal program **(Figure S6A-C).**

To evaluate myogenesis spatiotemporal development in musculoids, we applied principal graph learning on PCA space to reconstruct a differentiation tree using the scFates pipeline^96,97^. Linearity deviation assessment indicated linearity with low deviance (bridge populations) for top ranking genes (deviance <0.024) and continuity on musculoids curved trajectory. From an anterior primitive streak and posterior presomitic/somitic state, musculoids established an embryonic myogenic progenitors migration stream and promoted fetal maturation **(Figure 3H).** Differential expression and gene regulatory network analysis along pseudotime reveals forelimb pattering, e.g HOXA9, HOXB9, HOXC9 at presomitic (TBX6) and somitic (PAX3,FOXC1,FOXC2, TCF15) level, while upon hypaxial migration (LBX1,MET,MEOX2) level, we could detect progression till the stylopod level with HOXA10, HOXC10, HOXA11 upregulation **(Figure 4H, Figure S4M).** Following that level PAX3 myogenic progenitors ceased patterning, underwent embryonic to fetal transition, generated musculature and maturated till late fetal stage with MYF5, CD44, VEGFA and ECM related genes upregulation **(Figure 4H, Figure S4M,N).** At this stage, PAX7 myogenic progenitors positioned at the periphery of musculature and interacted with underlying myofibers via M-Cadherin, while by expressing CXCR4 they were still susceptible to attractive cues from their environment **(Figure 3A,I,L)**. Human skeletal muscle trajectory exhibits similar behavior by upregulating gene ontology terms related to vasculature development and ECM organization at late fetal/adult muscle stem cell stages **(Figure S5D,E).**

**Figure 4.**
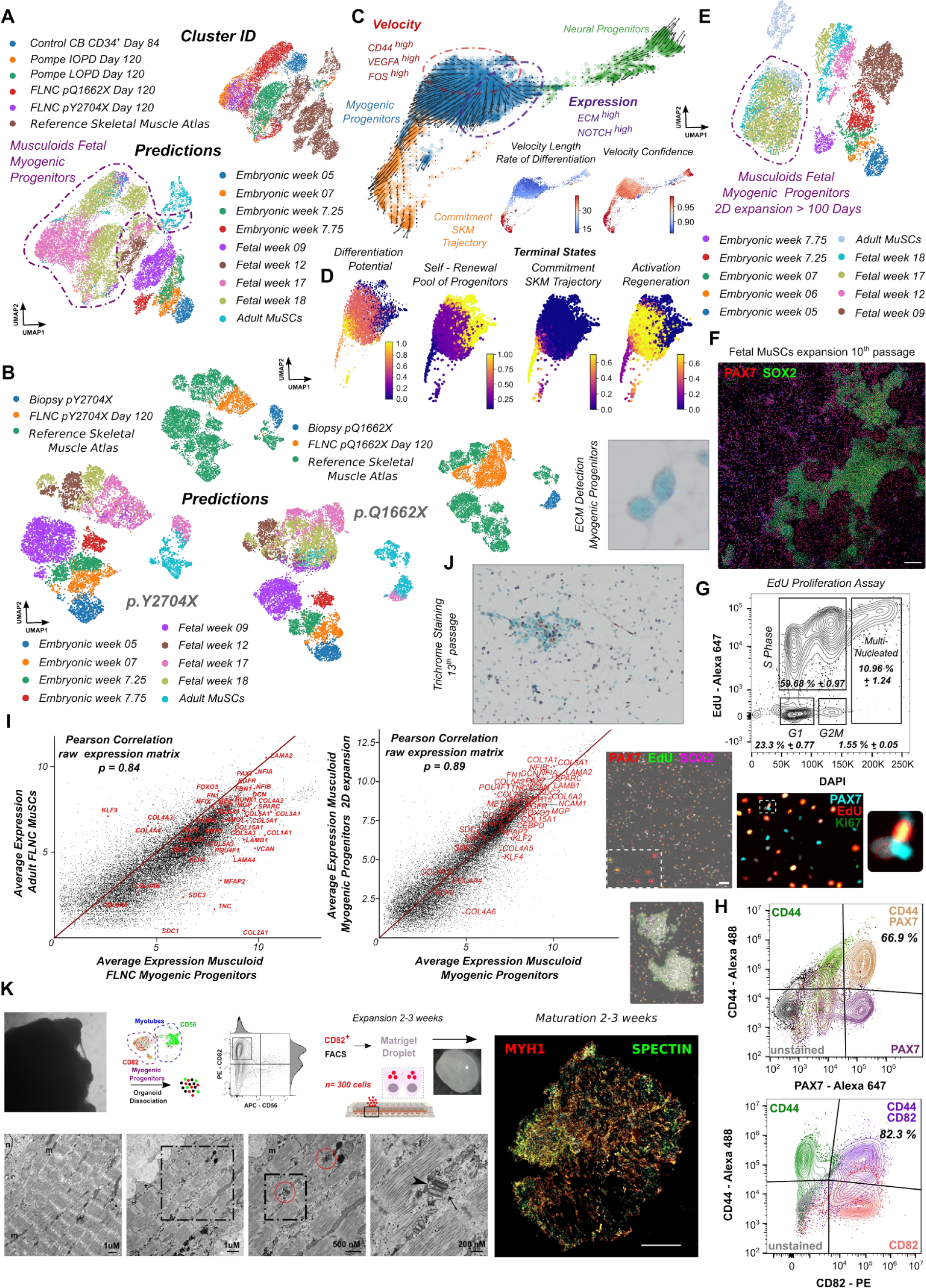
In vitro derivation and characterization of late fetal muscle stem cells. **(A)** UMAP plots based on a semi-supervised deep learning (SCANVI) approach to map GLM derived myogenic progenitors from Day84 - Day120 from different lines to the human skeletal muscle reference atlas predicts a maturation state between the late fetal (week 17-18) and adult muscle stem cells (MuSCs) identity. **(B)** Semi-supervised deep learning (SCANVI) approach to map biopsy derived MuSCs and their equivalent musculoid derived myogenic progenitors, predicts a late fetal (week 17-18) and adult muscle stem cells (MuSCs) identity for both biopsy and musculoid myogenic progenitors. **(C)** Force-directed k-nearest neighbor graph and RNA velocity analysis on a dataset from 2D expansion (>100 Days, 10 passages) late fetal myogenic progenitors (established at GLM level) and neural progenitors, indicates a pool of fetal myogenic progenitors with a high velocity rate for PAX7, ECM, Notch signaling following three independent trajectories related to self-renewal, activation/regeneration and commitment states. **(D)** Examining differentiation potential and cell fate probabilities during myogenic progenitor differentiation using Palantir algorithm, excludes bipotent cell fate probabilities, for self-renewal, activation/regeneration and commitment states. **(E) S**emi-supervised deep learning (SCANVI) approach to map GLM derived late fetal myogenic progenitors followed by 2D expansion (>100 Days, 10 passages) predicts a late fetal (week 17-18) and adult muscle stem cells (MuSCs) identity and demonstrates sustainability and self-renewal capability for musculoid myogenic progenitors. **(F)** Immunocytochemistry picture illustrating PAX7^+^ fetal myogenic and SOX2^+^ neural progenitors, established at GLM level (Day56 -Day 84) till late fetal stage, followed by 100 Days of 2D expansion. Scale Bar: 500um **(G)** Contour plots illustrating FACS cell cycle analysis on late fetal myogenic progenitors expanded for more than 100 Days in 2D. Before analysis cells incubated with 5uM EdU overnight (18hr). Immunocytochemistry images indicate proliferating (EdU^+^/MKI67^+^) and dormant (EdU^−^/MKI67^−^) fetal myogenic progenitors (PAX7^+^), and proliferating neural progenitors (EdU^+^/SOX2^+^). Scale Bar: 100um **(H)** Contour plots illustrating FACS quantification on CD82^+^, CD44^+^ and PAX7^+^ fetal myogenic progenitors established at GLM level (Day56 - Day84) till late fetal stage, followed by 100 Days of 2D expansion. Black contour: unstained population, green contour: CD44 FMO control, red contour: CD82 FMO control, magenta contour: PAX7 FMO control. **(I)** Scatter plot depicting average expression of genes between biopsy derived MuSCs and GLM derived myogenic progenitors from same patients at musculoid level (left, Day 120), and from GLM derived myogenic progenitors at musculoid level (Day 84) and GLM derived late fetal myogenic progenitors expanded in two dimension. Selected genes from the late fetal, adult MuSCs core program, *NFIX, MYF5, PAX7,* and genes related to specification resistance, *NOTCH, ECM,* are highlighted in red. **(J)** Trichrome staining on late fetal myogenic progenitors, followed by 2D expansion indicates ECM expression in single cells. **(K)** Scheme depicting an approach for expanding CD82^+^ myogenic progenitors on matrigel droplets and promote myofiber maturation. Immunocytochemistry picture depicts MYH1 and SPECTRIN expression on mature myofibers. Gating strategy for isolating CD82 pure positive myogenic fetal progenitors from organoid cultures for subsequent analyses. Scale Bar: 500uM. Ultrastructure images from myofibers after CD82^+^ myogenic progenitor FACS isolation and expansion on matrigel droplets. Before sarcomeres exhibit random orientation. Signs of maturity e.g. aligned sarcomeres, presence of triad like structure and striated plasma membrane are visible following CD82^+^ myogenic progenitor expansion. m: mitochondria, glyc: glycogen, n:nucleus. Red circles: triad-like structures. Arrow-head: T-Tubule, arrow: sarcoplasmatic reticulum.

Pseudotime analysis on gene regulatory networks (GRNs) and expression profiling during mouse, human and musculoid embryonic to fetal myogenic development, further indicates mesenchymal to myogenic developmental program and niche transition for myogenic progenitors, as we could detect N-Cadherin downregulation and M-Cadherin upregulation as differential expressed genes along myogenic progenitors’ developmental trajectory **(Figure 3J, Figure S4F,S5I).** We could further verify M-Cadherin expression on fetal myogenic progenitors and myofibers during fetal myogenesis occurring at the periphery of musculoids cultures **(Figure 3A,I).** On musculoids, fetal myogenesis onset occurred during 5^th^ week, on mouse forelimbs within E12.0 - E13.0 stage while in human hind-limbs during 7^th^ week of fetal development. At all cases, myogenesis followed limb patterning. This phenotype could also attribute the reduction on proliferation for fetal myogenic progenitors in a niche dependent manner, but more importantly sets an onset as structure and organ formation for the skeletal muscle system during development **(Figure S5E,I,J).** Force-directed *k*-nearest neighbor graph analysis demonstrated a dependent loop between myogenic progenitor maturation and commitment along embryonic-fetal stages. We could describe the presence of a skeletal muscle commitment trajectory that influences myogenic progenitor maturation during embryonic-fetal transition, as well as a secondary skeletal muscle commitment trajectory from myogenic progenitor mature stages that sustains skeletal muscle niche environment for myogenic progenitors during their growth phase **(Figure 3K,N, Figure S6K)**

Next using semi-supervised deep (SCANVI) learning classification and developmental score analysis we demonstrate that musculoid trajectory is equivalent to the human myogenic reference trajectory **(Figure 3F, Figure S6D).** We further verified through gene regulatory network analysis and deep learning (SCANVI) model that musculoid myogenic progenitors established a pool of progenitor cells with an identity between the fetal to adult stage **(Figure 3G,H, Figure S6A,B).** Pseudotemporal progression on myogenic progenitors indicated an absence of committed markers, such as MYOD1, MYOG and MYH3 (**Figure 4A)**, and further verified this “mosaic” profile, e.g. KLF4, CD44, CD82 and NFIX, between fetal and adult satellite cell markers on the 12^th^ week myogenic progenitors. At this stage, Myf5 expression on mouse fetal myogenic progenitors establishes the developmental progenitors of adult satellite cells^53^, a behavior that was observed during musculoid pseudotemporal trajectory and gene regulatory network analysis **(Figure S6C).** Subclustering on myogenic progenitors indicated that they reside in the activated, resisting specification and mitotic states. Progenitors in an activated, *CD44^+^, JUN^+^, FOS^+^*, undifferentiated state formed the main pool responsible for culture progression and bulge formation **(Figure S6E,H,I,J)**. Intercellular communication analysis indicated that late fetal musculoid derived myogenic progenitors were under the influence of NOTCH, laminin and collagen signaling. For those signaling pathways, inferred outgoing communication patterns suggested the mitotic and resisting cluster as the main secreting source and the activated cluster, through inferred incoming communication patterns, as their target **(Figure S6F,G)**. Manifold and classification learning analysis of signaling networks indicated functional similarity in activated and resisting specification cluster-related signaling pathways, further suggesting that these pathways exhibited similar/redundant roles in the fate of myogenic progenitors **(Figure 3O)**. During musculoid development, ECM expression appeared on myogenic progenitors first during the embryonic to fetal transition, followed by strong up-regulation at more mature stages in the 12^th^ week **(Figure 3M).** In human development similar appears during the late fetal stage (17^th^ −18^th^ week) and was also observed in mouse satellite cells during regeneration^109^ and in human satellite cells **(Figure 4I, Figure S7).**

### In vitro derivation and characterization of late fetal muscle stem cells

We further verified through semi-supervised deep learning (SCANVI) modelling to human reference skeletal muscle atlas that musculoid myogenic progenitors from different lines exhibit high reproducibility (Pearson correlation, p=0.93-0.96), and establish a pool of progenitor cells that express the core program of late fetal muscle stem cells, with an identity between the fetal to adult stage **(Figure 4A, Figure S6L)**. Next, we speculated that if our system truly harbors the derivation of fetal muscle stem cells, we would be able to long-term propagate them in an undifferentiated expanding state, since the default program of fetal muscle stem cells is to exhibit properties of resisting specification in cell autonomous manner^44^. Strikingly, in two cases, one upon dissociation and sorting/re-plating CD44, CD82 populations of the initial musculoid culture post 56 Days and, secondly via spontaneous detachment of myogenic progenitors from the initial musculoid culture (3D) followed by monolayer propagation in musculoid maturation medium, we were able to sustain them in an uncommitted, undifferentiated state for at least 10 passages (approx. 100 days) in two dimensions. In the second case, together with myogenic progenitors (n= 4037 cells, 73,82%), neural stem cells, (PAX6, SOX2, ASCL1-positive) were present in the culture but upon long-term propagation accounted only for a small fraction of the culture (n= 696 cells, 12,72%)**(Figure 4C)**. Characterization via FACS quantification indicates sustainable propagation for CD44, CD82 and PAX7 myogenic populations, with 60% of the total population to account for PAX7^+^ myogenic progenitors (passage #10) **(Figure 4H, Figure S8B).** Similar results are observed on the immunocytochemistry level at different passages for PAX7 and MYOD1-positive populations **(Figure 4F, Figure S8D,E).** Trichrome staining verifies ECM production on myogenic progenitors from that stage, a trait highly-associated with specification resistance **(Figure 4J).**. This observation was further verified by investigating their expression profiling at single cell resolution. RNA velocity analysis on myogenic progenitors propagated for 100 days (passage #10) highlights low rate of differentiation for myogenic progenitors and trajectories related to self-renewal, activation/regeneration and commitment **(Figure 4C).** Investigation on myogenic progenitor differentiation potential and cell fate probabilities using Palantir algorithm indicates that from a self-renewal state, CD44^+^, ECM^+^, NOTCH^+^ signalling upregulation, the pool of progenitors enter an activation regeneration state,*CD44^+^, FOS^+^,VEGFA^+^,* that leads to commitment, *MYH3^+^,MYOG^+^,* while self-renewal, activation states excludes cell fate probability towards commitment **(Figure 4C,D, Figure S8H).** Cell-cell communication analysis on musculoid derived myogenic progenitors expanded in 2D, further indicates cross-talk between myogenic progenitors states and self-regulation during specification resistance via NOTCH, laminin and collagen signaling pathways **(Figure S8F,G).**

Next, we investigated the cell cycle state of myogenic progenitors propagated in 2D via FACS and immunocytochemistry based EdU proliferation assay (overnight, 18hr). Strikingly, we could detect 23.3% ± 0.77 s.d of the cell population residing in G1 phase, predominantly presence of myogenic progenitors (SOX2^−^), while the majority of the cells reside in S phase (59.68% ± 0.97 s.d), and a small fraction (10.96% ± 1.24 s.d) to be multi-nucleated myotubes **(Figure 4G, Figure S8A,B)**. Consequently, myogenic progenitors via ECM upregulation were not only able to resist specification, but also to reside in a dormant non-dividing state. Trajectory analysis based on semi-supervised SCANVI deep learning model to human skeletal muscle reference atlas further highlights, similar to musculoid derived (3D, Day84), late fetal to adult muscle satellite cells (MuSCs) identity **(Figure 4E)**. Differential expression analysis between musculoid derived fetal myogenic progenitors and fetal myogenic progenitors expanded in 2D for 10 passages (100 days) highlight high similarity (Pearson correlation, p=0.89) with both expressing the core program, *ECM^+^, MYF5^+^,NFIX^+^, KLF4^+^,NCAM1^+^, MET^+^, EGFR^+^,* from late fetal muscle stem cells **(Figure 4I).**

Next we sought out to investigate whether our system establishes the developmental pool of adult muscle stem cells and whether adult and fetal populations share common properties. Since the musculoid approach simulates a constant skeletal muscle stem cell activation process, comparison should be made with a system that simulates an ongoing degeneration environment and promotes adult MuSCs activation, such as degenerative myopathies. To achieve this, we generated filaminopathy (FLNC) patient derived hiPSC lines and compared myogenic populations derived via the musculoid approach and those from corresponding patient biopsies suffering from a degenerative myopathy associated with FLNC aggregate formation^54^. Profiling skeletal muscle biopsies at single cell resolution **(Figure S9A,B,D,E)** and integrative comparison to human nuclei skeletal muscle reference map **(Figure S9G),** indicated an ongoing inflammation, with high upregulation of RUNX1, KLF6, LYVE1 on immune system related clusters **(Figure S9H)**. Cell-cell communication analysis demonstrated in a less severe case (p.Q1662X Biopsy), interactions between fibro-adipogenic progenitors, regenerating myofibers and immune clusters, while in a more severe case (p.Y2704X Biopsy), associated with increased fibrosis and proportion of immune cells, interactions only between fibro-adipogenic progenitors and immune system related clusters **(Figure S9C,F,G)**. Furthermore, the higher proportion of satellite cells in comparison to reference atlas and upregulation on gene regulatory networks, such as JUN, FOS and RUNX1, indicated an active regenerative state for MuSCs **(Figure S9I)**. Consensus clustering between p.Y2704X and p.Q1662X adult muscle stem cells indicates clear separate assignment for adult muscle datasets, while differential expression matrix for each cluster highlights presence of a cluster associated with myogenic commitment and active regeneration predominantly of the p.Q1662X Biopsy **(Figure S10F)**. Deep learning SCANVI model indicates musculoid progenitors as late fetal (week 18) but interestingly indicates FLNC satellite cells with a mosaic late fetal and adult identity **(Figure 4B).** This observation led us to in-depth investigate the dynamics present within both populations. Integrative and differentiation expression analysis between musculoid derived fetal myogenic progenitors and adult muscle stems cells from corresponding biopsies highlight high similarity (Pearson correlation, p=0.84) with both expressing the core MuSCs program, *ECM, MYF5,NFIX, KLF4,NCAM1, MET, EGFR* **(Figure 4I)**. This indicates default states for adult and fetal muscle stem cells related to specification resistance and regeneration potential. GRN network analysis indicates that adult p.Q1662X and pY2704X MuSCs within a degeneration environment upregulate the transcription factor RUNX1, a myogenic state, PAX7^+^/RUNX1^+^, that within musculoid culture corresponds to myogenic commitment and concludes to myogenic differentiation for myogenic progenitors**(Figure S10E).**, Hence, musculoid culture reassembles differentiation dynamics similar to *in vivo* states and serve as a platform to investigate the skeletal muscle stem cell activation process within a regenerating environment.

We reasoned that during ongoing regeneration in both systems, the niche environment for adult muscle stem cells and musculoid myogenic progenitors should be mainly the adjacent myofibers. Receptor - ligand analysis using NucheNet^104^ to infer ligand-receptor interactions between control and p.Y2704X, p.Q1662X FLNC skeletal muscle biopsies, and to detect expression of downstream target genes within the MuSCs, indicates presence of the MuSCs receptor repertoire, FGFR1, EGFR, NCAM1, ITGB1,MET, CDH15, NOTCH2, NOTCH3, and upregulating TGFB1 ligand activity on p.Y2704X, p.Q1662X FLNC skeletal muscle biopsies, a pathway that is described to promote cell cycle arrest and quiescence on adult MuSCs^55,56^, and during muscle regeneration in mouse model^109^ **(Figure S7G, S10C,D)**. Thus, niche environment at the biopsy level promotes a quiescent dormant state for MuSCs populations upon activation. On the contrary, receptor - ligand analysis between control and p.Y2704X, p.Q1662X FLNC musculoid derived regenerative myofibers and myogenic progenitors, highlights upregulation on BMP7 and IGF2 ligand activity, pathways described to regulate muscle growth and the generation of the adult muscle stem cell pool^57^, a trait of an active regenerative state in our system able to support the growth phase of the embryo **(Figure S10A,B)**. Reactome^106^ and gene ontology analysis on deferentially expressed genes between biopsy and GLM derived muscle stem cells highlights significant upregulation of markers (**Table S4)** associated with myogenic commitment on adult muscle stem cells and markers associated to ECM, cell cycle for fetal muscle stem cells. This is a clear distinction of the function between these sub-types during development, one to contributing to immediate muscle repair during postnatal stages and the other to growth during fetal development. **(Figure S10F,G)**.

Musculoids provide a developmental model of human hypaxial myogenesis that involves presence of variety of lineages and states at early stages, with secondary myogenesis occurring only at the periphery of the structure. Thus, at ultra-structural level, we detect functional sarcomeres on myotubes at random orientation^77^ (**Video S1).** To counteract this limitation, we sorted CD82 positive musculoid derived myogenic progenitors and re-plated them onto Matrigel droplets **(Figure 4K**, 300 cells per droplet). Interestingly, upon expansion and maturation we could establish myofiber networks that harbored synchronous contractions and contained mature aligned sarcomeres with presence of T-Tubules and triad-like structures **(Figure 4K, Video S2).** Isolating CD82^+^ myogenic progenitors from infantile and late onset Pompe lines at musculoid levels, followed by terminal differentiation, we could describe an early disease phenotype at myotube level by detecting significant glycogen accumulation via PAS immunocytochemistry staining and quantitative glycogen assay, when compared to wild type control line **(Figure S11B)**. In Morbus Pompe, a metabolic disease characterized via glycogen accumulation at the skeletal muscle level, it is reported in human patients and in mouse models that MuSCs maintain regenerative capacity but fail to repair disease-associated muscle damage, and it remains an open question whether the absence of an activating signal or the presence of an inhibitory factor from the niche environment contributes to this phenotype^58,59,60^. Using our musculoid approach to model muscle regeneration in infantile (IOPD) and late (LOPD) onset Pompe lines, we profiled both onsets at single cell resolution, where myogenic progenitors expressed the core fetal muscle stem cell program (**Figure S6L, S11A)**. Interestingly, receptor - ligand analysis to infer ligand-receptor interactions between control and infantile or late onset skeletal regenerating myofibers and myogenic progenitors, highlights TGFB1 ligand activity on LOPD and IOPD myogenic progenitors during regeneration (**Figure S11C-F)**, a pathway that promotes quiescence upon activation on adult and mouse MuSCs. Thus, this phenotype resembles the alterations of regenerative capacity, by affecting the transient amplification of MuSCs following their activation, but not on the pool of MuSCs of Pompe patients.

## Discussion

In summary, we report here a robust three-dimensional *in vitro* organoid model of skeletal muscle organogenesis at forelimb level of human development using human PSCs. At the gastrulation stage, GLMs exhibit spatiotemporal organization **(Figure 5C)** and establish developmental trajectories for skeletal muscle, neural crest, lateral plate mesoderm and fore-gut endoderm lineages. GLMs, as a culture system uniquely model migration events along the medio-lateral and proximal-distal axes for lineages that shape body patterning during embryogenesis, while when comparing to the human developing embryo their patterning is equivalent to Carnegie Stage CS 13/14 **(Figure 5)**. Continuous SF/HGF stimulation simulated culture conditions that during embryogenesis control the migration of hypaxial precursors from the dermomyotomal lip into the limb bud^62^, thereby allowed us to promote skeletal muscle lineage patterning and maturation until the 18^th^ week of human fetal development.

**Figure 5.**
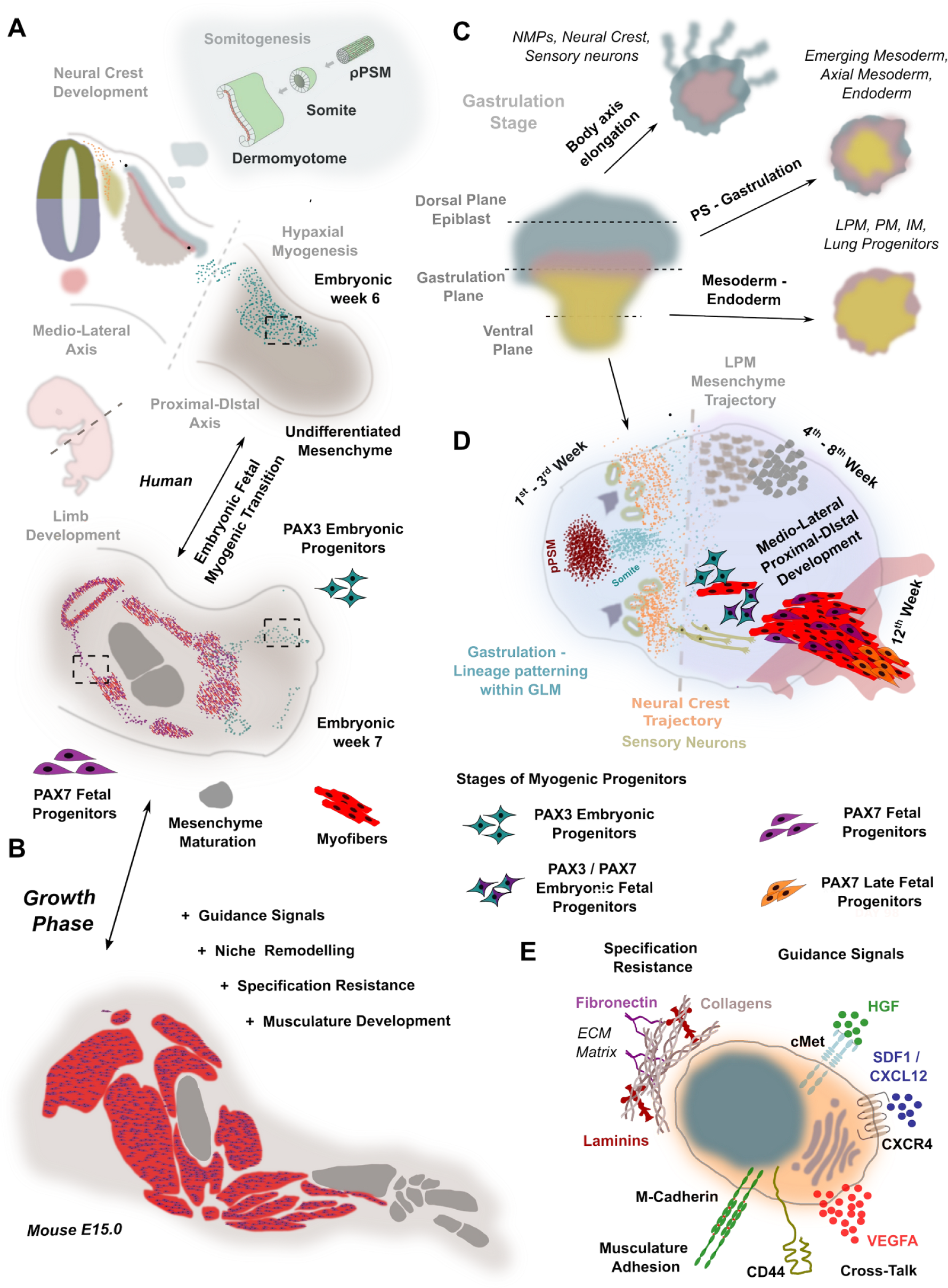
Gastruloids-Lateraloids-Musculoids (GLM) reassemble skeletal muscle lineage development at forelimb level. **(A)** A schematic representation of skeletal muscle lineage development during embryonic/fetal stages at forelimb region (transverse plane) during human fetal development **(B)** A schematic representation of skeletal muscle lineage growth during fetal stages at forelimb region. **(C)** Scheme explaining lineage spatial arrangement at gastrulation stage during GLM early patterning **(D)** A schematic comparison of a typical GLM migration dynamics (plane view), with key stages and lineage development along the medio-lateral, proximal-distal axes following gastruloid patterning for muscle organogenesis. **(E)** Scheme illustrating a late fetal muscle stem cell and highlighting its main functions on niche remodeling. resisting specification, guide signals, attachment / generating musculature and attracting vasculature via VEGFA expression.

In GLMs and during human and mouse limb development, the embryonic to fetal transition for myogenic progenitors coincided with transition for their niche environment, that was further associated from mesenchyme to myofiber development. In a mouse model, it is reported that only embryonic myoblasts express a Hox code along the antero-posterior axis^61^. Here we further describe that during human and mouse fetal limb myogenesis and the musculoid hypaxial migration stage, the cMET/PAX3 myogenic progenitors under the influence of mesenchyme as their niche environment, upregulated gene regulatory networks related to axial anatomical identity and were mitotically active. Following the embryonic to fetal myogenic transition, the PAX7 fetal myogenic progenitors, by switching their niche environment to myofibers, are characterized by ECM upregulation, prolonged G1 phase and downregulation of HOX gene expression. In line, replacing PAX3 with PAX7 gene in mouse embryos caused detrimental deficits at the stage of migration and proliferation for myogenic progenitors and lead to severe musculature defects in a proximal – distal manner at later stages^63^.

During embryogenesis, this transition ideally correlates with the end of patterning and the start of growth for a specific region along the proximal distal axis during human limb development, a developmental process that would require potent myogenic progenitors able of resisting specification and promoting self-renewal. In a growth model for myogenic transition, PAX3 myogenic progenitors at the hypaxial migration stage should first position within the fore/hind-limb (E10.5 – E12 in mouse, 5^th^ week to 7^th^ week in humans), followed by embryonic to fetal myogenic transition. Such model would require an early axial anatomical identity specification followed by an ability for PAX7 myogenic progenitors to resist specification in a cell autonomous manner and still be able to respond to guidance signals from their surroundings. Consequently, myogenic progenitor maturation could be divided into two stages during fetal development. One where they abolish their embryonic identity (axial/anatomical specification) and a second where via musculature interaction they maturate and unlock their late fetal/ adult stem cell identity.

Bio-engineering approaches that simulate myofibers niche environment succeed to preserve short-term quiescence of adult muscle stem cells but fail to promote long-term expansion^64^, while 2D and 3D hiPSCs myogenic differentiation approaches by mimicking development generate myogenic progenitors^32,48,65,66,67,68,69,71^ profiled with an upregulated anatomical embryonic developmental program (**Figure S12),** a myogenic state that is associated with the embryonic-early fetal transition stage and is characterized as the developmental barrier for *in vitro* differentiation approaches^52^. To date, in a proof of concept study^72^, only in vivo matured human PAX7 myogenic progenitors, following engraftment into immunodeficient mice, surpass the embryonic-fetal barrier, supporting our claim of advanced in vitro maturation in our organoid system. Here, we demonstrate that musculoid culture, under minimal conditions, establishes and promotes maturation of late fetal myogenic progenitors which are further associated with long-term *in vitro* expansion, self-renewal and specification resistance. Furthermore, musculoids’ radial extension uniquely facilitates the distinction between each developmental stage of myogenic progenitors **(Figure 5D)**.

Altogether, GLMs succeed to reassemble the *in vivo* skeletal muscle environment at all developmental stages of human fetal myogenesis at forelimb level and are able to promote *in vitro* PAX7^+^ myogenic progenitor maturation, which are able to resist specification in a cell autonomous manner **(Figure 5E)**. In conclusion, our culture system presents i.) a novel and detailed organoid developmental model for investigating mechanisms of human skeletal muscle organogenesis, ii.) for the first time the *in vitro* derivation of fetal muscle stem cells and iii.) provides new insights on disease onset and mechanisms for congenital muscular dystrophies and on neural crest, lateral plate and fore-gut development following gastrulation.

## Acknowledgments

We are grateful to Drs. Karl Köhrer, Tobias Lautwein, Patrick Petzsch, and Thorsten Wachtmeister, Genomics & Transcriptomics Laboratory, Heinrich-Heine-University Düsseldorf for performing single-cell RNAseq experiments with their Illumina HiSeq platform and data provision. We are grateful to Anja Tietz, Institute of Translational Neurology, University Hospital Münster, Dr. Anika Witten and Prof. Dr. Monika Stoll, Core Genomics Facility Medical Faculty Westfalische Wilhelms-Universitäts Münster for performing single cell experiments with their Illumina platforms. We are grateful to Dr. Kristina Döring, Theodora Wange and Prof. Dr. med. Huu Phuc Nguyen for their assistance and access on Tapestation (Agilent) automated gel electrophoresis system. We thank Dirk Richter for helping with software/hardware installation/troubleshooting. We further would like to thank Ingrid Gelker, Manuela Haustein, Karina Mildner, Max Planck Institute Münster and Rana Houmany, Boris Burr, Ruhr-University Bochum as well as Michaela Zaik, Anja Schreiner, Janine Mertens-Rill, Heimer Institute for Muscle Research, BG-University Hospital Bergmannsheil for their technical assistance. This work was supported by research grants from the Technology Innovation Program (20015148, Development of Neural/Vascular/Muscular-Specific Peptides-conjugated Bioink and Volumetric Muscle Tissue) funded by the Ministry of Trade, Industry & Energy (MOTIE), South Korea.

## Author contributions

L.M. conceived the study, designed and performed experiments, single cell and spatial RNAseq experiments and bioinformatic analyses, interpreted the results, and wrote the manuscript with input from all authors. N.M.D, L.V, H.Z, D.Z., M.S., performed experiments and analyzed data, I.N.L, H.W.J., J.H.Y., G.M.z.H., performed single cell RNAseq experiments B.B.S., H.R.S., M.V. H.Z., acquired funding, supervised study, provided study materials and a collaborative environment. All authors read and approved the final manuscript.

## Methods

### hiPSCs culture

Human induced pluripotent stem cell (hiPSC) lines, Cord Blood iPSC^73^, FLNC p.Q1662X, HIMRi001-A, and FLNC p.Y2704X, HIMRi005-A^74,75^,IOPD, HIMRi006-A and LOPD, HIMRi007-A^76^, Gibco episomal iPSCs (line A18945) and lines described in^77^, were cultured in TESR-E8 (StemCell Technologies) or StemFlex (ThermoFischer Scientitfic) on Matrigel GFR (Corning) coated 6 well plates. Patient cells were collected at the University Hospital Bergmannsheil. Ethical approval was obtained from the ethics committee of the Ruhr-University Bochum, Medical Faculty (15-5401, 08/2015).

### GLM differentiation protocol

Prior differentiation, undifferentiated human iPSCs, 60-70% confluent, were enzymatically detached and dissociated into single cells using TrypLE Select (ThermoFisher Scientific). Embryoid bodies formed via the hanging drop approach, with each droplet containing 4×10^3^ human single PSCs in 20 μl were cultured hanging on TESR-E8 supplemented with Polyvinyl Alcohol (PVA) at 4mg/ml (SigmaAldrich) and rock inhibitor (Y-27632) at 10μM (StemCell Technologies) at the lid of Petri dishes. The next day, embryoid bodies at the size of 250-300μm embedded into Matrigel and cultured in DMEM/F12 basal media (ThermoFisher Scientific) supplemented with Glutamine (ThermoFisher Scientific), Non Essential Amino Acids (ThermoFisher Scientific), 100x ITS-G (ThermoFisher Scientific), (Basal Media) 3μM CHIR99021 (SigmaAldrich) and 0.5μM LDN193189 (SigmaAldrich). On day 3, human recombinant basic Fibroblast Growth Factor (bFGF) (Peprotech) at 10ng/ml final concentration was added to the media. Subsequently, on day 5 the concentration of bFGF was reduced at 5ng/ml and the media was further supplemented with 10nM Retinoic Acid (SigmaAldrich). To promote axial elongation the cytokine and growth factor cocktail, CHIR99021 (3μM), LDN193189 (0.5μM), bFGF (10ng/ml), Retinoic Acid (10nM), was applied from day7 to day11. Alternative the differentiation media during standard GLM differentiation protocol on day 7, supplemented only with human recombinant Sonic hedgehog (hShh) (Peprotech) at 34ng/ml, human recombinant WNT1A (Peprotech) at 20ng/ml and 0.5μM LDN193189. On day 11 the cytokine composition of the media was changed to 10ng/ml of bFGF and human recombinant Hepatocyte Growth Factor (HGF) at 10ng/ml (Peprotech). From day 15 onwards, the basal medium supplemented with ITS-X (ThermoFisher Scientific) and human recombinant HGF at 10ng/ml. In the first 3 days of the differentiation the medium was changed daily, from 3^rd^ till 30^th^ every second day, while from day 30 onwards every third day.

### Expanding musculoid derived fetal muscle stem cells in two dimensions (2D)

Musculoids exhibiting the typical protrusions that contain myofibers and myogenic progenitors (Figure 4I,N) post 8^th^ week of three dimensional development, at 12^th^ week, either: a) dissociated into single cells by incubation at 37°C with TryplE Select for 10 min, followed by surface antigen staining with PE-labelled anti-human CD82 (Biolegend, clone TS2/16) and Alexa 488-labelled anti-human CD44 (eBioscience, clone IM7, FITC) antibodies for 20 min incubation on ice. Cells washed twice with 1% BSA staining solution and before FACS sorting, the dissociated cells were passed through a 70 μm cell strainer to remove any remaining aggregates.. Fluorescent minus one (FMO) controls were used for correct gating **(Figure 4H)**. CD44/CD82 positive sorted cells were seeded on matrigel coated plated in musculoid maturation media (DMEM/F12, NEAA, P/S, L-Glutamine, + 10nh/ul HGF) supplemented with 10uM Rock inhibitor at a density of 150000 −200000 cells per cm^2^. Or b) By transfering them with 1 mL cutted pippete tip to new matrigel coated plated. Musculoids by an attaching-detaching process leave myogenic progenitors, myofibers on the matrigel surface, that upon propagation reach 100% confluency. In both approiaches, for 2D expansion myogenic progentiors are passaged at 90%-100% confluncy at 1:2 ratio. Myogenic progenitors are passaged enzymatically with TryplE Select and the musculoid maturation media the day of passaging contains rock inbitor (10uM), that is ommited the day after. Media change occurs every second date, as myogenic progenitors require active HGF within the media for expansion/propagation.

### Replating CD82^+^ myogenic for secondary skeletal muscle-like organoid formation

#### Re-plating CD82^+^ myogenic to establish myofiber networks and promote myofiber maturation

Post 8w-12w musculoids were dissociated into single cells by incubation at 37°C within papain solution for 1-2 h, followed by incubation with TryplE Select for 10 min. For surface antigen staining cells were incubated for 20 min with APC-labelled anti-human CD56 (Biolegend, clone TS2/16) and PE-labelled anti-human CD82 (Biolegend, clone ASL-24) antibodies and washed twice with 1% BSA staining solution. Then the dissociated cells were passed through a 40 μm cell strainer to remove any remaining aggregates. Briefly before FACS sorting to discriminate between dead and live cells DAPI was added to the samples. DAPI-negative / CD82 positive cells were collected using a FACSAria Fusion cell sorter (BD Biosciences). Fluorescent minus one (FMO) controls were used for correct gating **(Figure 4K).** Before sorting 300 CD82 positive events/cells into each well of a 96 well plate, we generated a matrigel droplet in each well. For that we applied 30ul of matrigel in each well and let it polymerize for 20-30min. Upon matrigel polymerization we fill the well with 150ul of DMEM/F12, ITS-X basal media supplemented with 10ug/ml HGF. Upon sorting the cells the next day and for the next 2 media changes, we proceeded with 50% Basal media (+10uh/ml HGF) and 50% SkGM™-2 Skeletal Muscle Cell Growth Medium (Lonza). Subsequently, we shifted completelly to Skeletal Muscle Cell Growth Medium till cells covered the whole matrigel sufcace before we induce maturation with DMEM/F12, ITS-X, N2 muscle fushion maturation media. Upon that stage cells were cultured up to a month. At the expansion phase (Lonza Skeletal Muscle Cell Growth Medium) media changed daily while during maturation every second day.

### Immunocytochemistry

#### Cryosection Immunochemistry

Organoids from different stages were fixed on 4% paraformaldehyde overnight at 4°C under shakings conditions, dehydrated (30% sucrose o/n incubation) and embedded in OCT freezing media. Cryosections were acquired on a Leica CM3050s Cryostat. For the immunostaining process, cryosections were rehydrated with PBS and followed by permeabilization once with 0.1% Tween-20 in PBS, (rinsed 3x with PBS), and then with 0.1% Triton-X in PBS (rinsed 3x with PBS). Subsequently, the sections were blocked with 1% BSA / 10% NGS or 10% FBS in PBS for 1hr at room temperature. Primary antibody incubations were performed overnight at 4°C, where secondary antibody incubations for 2hr at room temperature.

#### EdU staining

At 180 days post diferentiation, 12 weeks for myogenic progenitor maturation at musculoid level, and 100 Days expansion in 2D (10 passages), musculoid derived myogenic progenitor incubated overnight (18 hr) with EdU at a final concentration of 5 μM. To detect EdU, the sections were processed with Click-iT EdU Alexa Fluor 647 cell proliferation kit (Invitrogen) following the manufacturer’s instructions. The samples were incubated with secondary antibodies after the click reaction for detecting EdU.

#### Primary Antibodies

anti-Brachyury/TBXT (R&DSystems, 1:250), anti-TBX6 (R&DSystems, 1:200), anti-PAX3 (DHSB, 1:250), anti-PAX7 (DHSB, 1:250), anti-PAX6 (Cell Signalling, clone D3A9V, 1:200), anti-SOX10 (R&DSystems,1:125), anti-FOXA2(R&DSystems,1:200), anti-Ki67 (ThermoFisher Scientific, clone SolA15, 1:100), anti-MYH1 (DHSB,1:100), anti-MYOD1 (Cell Signalling, D8G3, 1:200), anti-TFAP2A (DHSB, 3B5, 1:100), anti-SOX2 (ThermoFisher Scientific, clone Btjce, 1:100; Cell signalling, clone D6D9, 1:200), anti-CD44 (eBioscience, clone IM7, 1:100), anti-NeuN (Abcam, EPR12763, 1:200), anti-CDX2 (Biogenex, clone CDX2-88, 1:200), anti-GATA6 (Cell Signalling, clone D61E4, 1:200), anti-Desmin (Abcam, clone Y66, 1:200), anti-GSC (R&DSystems, 1:200), anti-SOX17 (R&DSystems, 1:200), anti-PAX2 (Biolegend, clone Poly19010, 1:200), anti-WT1 (Cell Signalling, clone D8I7F, 1:200), anti-MET (Cell Signalling, clone D1C2, 1:200), anti-NANOG (Cell Signalling, clone D73G4, 1:200), anti-M-Cadherin (Cell Signalling, clone D4B9L, 1:200), anti-Spectrin (Novocastra, NCL-SPEC1,1:30), Mouse anti-FastMyHC (SigmaAldrich, clone MY-32, 1:300).

#### Secondary antibodies

Alexa Fluor® 647 AffiniPure Fab Fragment Goat Anti-Mouse IgM, μ Chain Specific (Jackson Immunoresearch Laboratories, 1:100), Rhodamine RedTM-X (RRX) AffiniPure Goat Anti-Mouse IgG, Fcγ Subclass 1 Specific (Jackson Immunoresearch Laboratories,1:100), Alexa Fluor® 488 AffiniPure Goat Anti-Mouse IgG, Fcγ subclass 2a specific (Jackson Immunoresearch Laboratories,1:100), Alexa Fluor 488, Goat anti-Rat IgG (H+L) Cross-Adsorbed Secondary Antibody, (ThermoFisher Scientific, 1:500), Alexa Fluor 488, Donkey anti-Mouse IgG (H+L) Cross-Adsorbed Secondary Antibody, (ThermoFisher Scientific, 1:500), Alexa Fluor 647, Donkey anti-Goat IgG (H+L) Cross-Adsorbed Secondary Antibody, (ThermoFisher Scientific, 1:500), Alexa Fluor 488, Donkey anti-Goat IgG (H+L) Cross-Adsorbed Secondary Antibody, (ThermoFisher Scientific, 1:500), Alexa Fluor 568, Donkey anti-Rabbit IgG (H+L) Cross-Adsorbed Secondary Antibody, (ThermoFisher Scientific, 1:500). Images were acquired on a ZEISS LSM 770 inverted confocal microscope.

### Flow Cytometry

#### EdU assay

At 180 days post diferentiation, 12 weeks for myogenic progenitor maturation at musculoid level, and 100 Days expansion in 2D (10 passages), musculoid derived myogenic progenitor incubated overnight (18 hr) with EdU at a final concentration of 5 μM. The next day musculoid derived myogenic progenitor cultures were dissociated into single cells by incubation at 37°C within TryplE Select solution for 5-10 min. Then the dissociated cells were passed through a 40 μm cell strainer to remove any remaining aggregates. To detect EdU, the cells were processed with Click-iT EdU Alexa Fluor 647 Flow Cytometry Assay Kit (Invitrogen) according to manufacturer instructions, nuclei werestained with Dapi solution and then analyzed on a FACSAria Fusion flow cytometer (BD Biosciences). FACs data fcs files were processed with FlowJo v10 (BD Biosciences).

#### FACS isolation and quantification of CD44^+^, CD82^+^, PAX7^+^,MYOD1^+^, CXCR4^+^ myogenic cell population^78,79^

Organoids during 8^th^ - 16^th^ week post differentiation were dissociated into single cells by incubation with Papain solution till we could observe complete dissociation upon gentle shaking (30 min – 1 h). To acquire singlets, the cells were filtered through a 40 μm cell strainer and washed with 1% BSA solution. For surface antigen staining cells were incubated for 20 min with FITC-labelled anti-human CD44 (eBioscience, clone IM7) and PE-labelled anti-human CD82 (Biolegend, clone ASL-24) antibodies and washed twice with 1% BSA staining solution. Briefly before FACS sorting to discriminate between dead and live cells DAPI was added to the samples. DAPI-negative / CD82 positive cells were collected using a FACSAria Fusion cell sorter (BD Biosciences). Fluorescent minus one (FMO) controls were used for correct gating (Figure 4H). For CXCR4 quantification, the PE anti-human CD184[CXCR4] (Biolegend, clone 12G5) was applied together with the corresponding isotype control for setting the gating: PE Mouse IgG2a, κ Isotype Ctrl antibody (Biolegend, clone MOPC-173) (Figure 3L)..

### Single cell RNA sequencing expression profiling

#### Organoid Samples and cDNA library preparation

Single cells acquired in suspension following 1hr incubation with solution containing papain and EDTA for organoids at stages of ongoing development within the matrigel droplet, e.g. Day 7, Day 13, Day 19, Day 35, Day 56, Day 84, or with 15-20 min TryplE incubation for 2D cultured (>100 Days) musculoid derived myogenic progenitors and for organoids at stages that the organoid development takes place at bulge/protrusion sites, e.g. Day 120 organoids. After dissociation, cell number and viability was estimated, cells were re-suspended on solution containing 0.5% BSA and processed using the Chromium Single Cell 3′ Reagent Kits (v3): Single Cell 3′ Library & Gel Bead Kit v3 (PN-1000075), Single Cell B Chip Kit (PN-1000073) and i7 Multiplex Kit (PN-120262) (10x Genomics) according to the manufacturer’s instructions. Then, the cDNA library was run on an Illumina HiSeq 3000 or Novoseq as 150-bp paired-end reads.

### Single-nuclei isolation of human skeletal muscle biopsies

Skeletal muscle tissues from Vastus Lateralis (p.Y2704X) and Soleus (p.Q1662X) were sampled by the surgeon and immediately frozen in liquid nitrogen. Single-nuclei isolation was performed on ice using the Chromium nuclei isolation kit according to the manufacturer’s instructions. After dissociation, nuclei number was estimated and processed immediately using the Chromium Single Cell 3′ Reagent Kits (v3).

### Spatial gene expression assay

Frozen Gastruloids/Lateraloids/Musculoids samples from Day 5, Day 9, Day19 and Day 35 were embedded in OCT (Tissue-Tek) and cryosectioned (Thermo Cryostar). The 12-µm section was placed on the pre-chilled Optimization slides (Visium, 10X Genomics, PN-1000193) and the optimal lysis time was determined. The tissues were treated as recommended by 10X Genomics and the optimization procedure showed an optimal permeabilization time of 18 min of digestion and release of RNA from the tissue slide. Spatial gene expression slides (Visium, 10X Genomics, PN-1000187) were used for spatial transcriptomics following the Visium User Guides. Brightfield histological images were taken using a 20X objective on the Olympus IX83 fluorescent inverted microscope and images were stitched and analyzed with the cellSens software. Next generation sequencing libraries were prepared according to the Visium user guide. Libraries were loaded at 300 pM and sequenced on a NextSeq 1000 System (Illumina) as recommended by 10X Genomics.

#### Single cell RNA seq analysis

Sequencing data raw files were processed using Cell Ranger software (v5.0.1), following the set of analysis pipelines suggested by 10x Genomics, and the reads aligned to the human genome and transcriptome (hg38, provided by 10x Genomics) with the default alignment parameters. For constructing the GLM developmental trajectory presented in figure 2a we included intronic reads, while for musculoid developmental trajectory and myogenic subsequent myogenic comparison we excluded intronic reads at the default alignment Cell Ranger parameters. Seurat^80,81,82,83,84^ or ScanPy^85^ pipeline was then applied for further data processing. For Seurat based normalization we followed the SCT-transform approach. To denoise the graph, in ScanPy pipeline we applied the MAGIC approach^86^, an unsupervised non-parametric algorithm to impute and de-noise biological single-cell RNA-seq data sets. Interoperability between Seurat and anndata was achieved via *h5 Seurat* function, to generate h5ad objects, while for generating loom files via *as.loom* function. Before comparing datasets genes related to metabolism such as mitochondrial and ribosomal genes, were excluded from subsequent analysis. In both approaches, the sequencing depth, proportions of mitochondrial transcripts, cell cycle effects^87^, and when analyzing skeletal muscle lineage genes associated to stress during myogenic progenitor dissociation^88^ were also regressed out. Unless otherwise noted, cells with less than 300 detected genes and those with mitochondrial transcripts proportion higher than 5 percent were excluded. For myogenic comparison between reference and myogenic differentiation datasets cells with less than 500 detected genes and genes detected in less than 5 cells we excluded form analysis, exception was only adult and juvenile datasets^52^, where the threshold was adjusted to 200-300 genes per cell and 3 cells per gene. Finally, as myogenic progenitor related clusters were considered the ones expressing muscle related genes e.g. *PAX3, PITX2, PAX7, SIX1, MYOD1* in the absence of neural markers. For consensus clustering of p.Q1662X and p.Y2704X FLNC adult muscle stem cells, first we converted seurat object into SingleCellExperiment objects, followed by SC3 R package pipeline^89^.

#### Spatial transcriptomics analysis

Sequencing data raw files were processed using Spatial Ranger software (v2.1.1), following the set of analysis pipelines suggested by 10x Genomics, and the reads aligned to the human genome and transcriptome (hg38, provided by 10x Genomics) with the default parameters. Seurat or ScanPy pipeline was then applied for further data processing and visualization. For mapping the single-cell and spatial transcriptomic datasets we applied the CytoSPACE pipeline^90^, where labeled single-cell expression matrices mapped onto coordinates (spots) of spatial transcriptomic datasets. Before mapping single cell and spatial datasets, we subset the spatial dataset to avoid any spatial spots with no expression values (nCount_Spatial>0), and removed ribosomal and mitochondrial genes from the expression matrices.

#### Gene regulatory network analysis

For analyzing and clustering datasets based on their gene regulatory network we applied pySCENIC pipeline^91^ on Scanpy normalized datasets. Cells with more than 6,000 or less than 200 detected genes, as well as those with mitochondrial transcripts proportion higher than 5% were excluded. Sequencing depth and proportions of mitochondrial transcripts and cell cycle effects, and genes associated to stress during myogenic progenitor dissociation when analyzing skeletal muscle lineage only, were also regressed out. When merging datasets for comparative analysis genes related to metabolism such as mitochondrial and ribosomal genes, were excluded from the normalised matrix. For GRN analysis we applied genes as ranking type and therefore we applied hg38 cisTarget databases to all datasets, (https://resources.aertslab.org/cistarget/).

#### Pseudotime analysis

GLM datasets processed with Scanpy and upon normalisation, cell cycle genes, sequencing depth and stress related genes were regressed and we further proceeded with the generation of Force-directed layouts of single-cell graphs using the ForceAtlas2 algorithm^92^. To denoise the graph, we projected the dataset in DiffusionMap space and for calculating the trajectory inference we applied the Partition-based graph abstraction (PAGA) approach^93^. Pseudo-spatiotemporal orderings were constructed by randomly selecting a root cell from the following clusters: somitogenesis cluster **(musculoid trajectory, Figure 3H)**; Anterior Primitive streak cluster **(Somitogenesis, Figure 2B)** and calculating the diffusion pseudotime distance of all remaining cells relative to the root. On Seurat normalised datasets we calculated the pseudotime by converting them into SingleCellExperiment objects, calculate the eigenvector values for each cell using destiny package (k=100) and order them by diffusion map pseudotime^94^. During ScanPy pipeline, for investigating cell fate probabilities and align cells along differentiation trajectories we applied Palantir algorithm and pipeline^95^.

#### Advanced Pseudotime calculation

To simulate *in silico* myogenesis on musculoids, we applied principal graph learning on PCA space to reconstruct a differentiation tree using the ScFates package^96^, following the basic curved trajectory analysis pipeline. For principal graph learning we applied EIPiGraph^97^. In order to verify that the trajectory we are seeing is not the result of a linear mixture of two population (caused by doublets), we performed the Linearity deviation assessment test where we could describe continuity with putative progenies (paraxial mesodermal clusters), putative bridge (hypaxial migration and embryonic to fetal transition) and putative progenies (late fetal myogenic progenitors).

#### Integrative Mapping

For integrative analysis we used the Seurat(v4) package for finding anchors between reference atlases and those from differentiation protocols, using 5000 anchors (SCT transform) or 2000 anchors (log Normalization) and regressing out cell cycle genes, sequencing depth and stress related genes before integration. For predictions based on integrative analysis, following FindTransferAnchors step we applied the TransferData option from Seurat in Seurat object metadata.

#### Developmental, anatomical, fetal/postnatal and myogenesis score

Developmental score was calculated as described^52^. Briefly, we used the ‘‘AddModuleScore’’ function to calculate embryonic and adult score using a list of differentially expressed genes (DEGs) between satellite cells and embryonic myogenic progenitor clusters. Differentially expressed genes (DEGs) were identified by ‘‘FindMarkers,’’ function using ‘‘MAST’’ test^98^. In addition, we passed the same parameters as we did when scaling the data (‘‘S.Score,’’ ‘‘G2M.Score,’’ ‘‘Stress’’ and ‘‘total Count’’) to the vars.to.regress’ argument to regress out the effects of the cell cycle, dissociation-related stress as well as cell size/sequencing depth on the identification of DEGs. The developmental score was further calculated by subtracting embryonic from the adult score. The list of genes for calculating the embryonic and adult score are listed in **Table S1**. Anatomical, embryonic TF, fetal/postnatal TF, ECM and myogenesis score calculation were calculated via Seurat′s ‘‘AddModuleScore’’ function. List of genes for calculating each score are listed in **Table S2**. List of genes for calculating embryonic TF, fetal/postnatal TF score we applied the genes described in human skeletal muscle reference atlas^52^.

#### Deep learning classification

Seurat normalised datasets converted to anndata object via h5.seurat. For deep learning classification the raw.matrix from each dataset was used. From the Scarches package^99^, the semi-supervised deep learning approach was applied using single-cell annotation variational inference (SCANVI) algorithm^100^ for training the reference atlas and transfer existing neural networks to model and train query datasets.

#### RNA velocity

For generating count matrices for the pre-mature (unspliced) and mature (spliced) RNA abundances in our samples we applied the velocyto pipeline^101^ on FASTQ files. Subsequently, we pre-processed datasets using Scanpy pipeline and for calculating the RNA velocity we applied the ScVelo package pipeline^102^.

#### Cell-Cell communication analysis

To investigate cell-cell communications among activated, mitotic, specification resistance myogenic progenitors and myofiber related clusters from 12^th^ week musculoids we applied the CellChat R package^103^, while for modeling intercellular communication by linking ligands to target genes and identify receptor-ligand interactions between musculature and MuSCs on mouse, p,Q1662X/p.Y2704X Biopsies or musculoid myogenic progenitors datasets we applied the Nichenet R package^104^

#### Gene ontology enrichment analysis

Gene ontology (GO) enrichment was performed on Differentially expressed genes (DEGs), genes are listed in listed in **Table S3,S4**. using Metascape^105^ (http://metascape.org/gp/index.html#/main/step1) against GO terms belonging to “Biological Processes” and “Hallmark Processes”. Reactome analysis^106^ (https://reactome.org/) was performed on differentially expressed genes (DEGs) between GLM derived fetal and biopsy derived adult MuSCs p.Q1662X,p.Y2704X lines. Genes are listed in in **Table S4**.

### Transmission Electron Microscopy (TEM)

Organoid cultures were prepared for electron microscopy according to standard protocols. First, they were fixed in 2 % glutaraldehyde, 2 % paraformaldehyde in 0.1 M cacodylate buffer, pH 7.2 for at least 3 hours at room temperature. Before embedding samples were postfixed in 1 % osmium tetroxide including 1.5 % potassium cyanoferrate and dehydrated stepwise in ethanol with finally 0.5 % uranyl acetate en-bloc staining during 70 % ethanol. Then the samples were infiltrated in epon. Ultrathin sections of the polymerized sample blocks were cut in different orientation to cover all morphological features of the organoid culture. Representative pictures were imaged at the electron microscope (Tecnai 12-biotwin, Thermofisher scientific, the Netherlands) with a 2K CCD camera (Veleta, EMSIS, Muenster).

### Trichrome staining (Gomori)

Gomori’s trichrome staining was conducted using a ready-to-use kit (Trichrome Stain (Gomori) Kit, HT10, Sigma-Aldrich) as described by the manufacturer.

### Glycogen assay

Samples (three biological replicates) were washed twice with PBS and harvested in Ampuwa® using a cell scraper. The cell lysate was immediately frozen and stored in liquid nitrogen The amount of glycogen was determined as duplicates by a glycogen assay kit, which was used according to the manufacturer’s protocol for colorimetric quantification. An Agilent BioTek ELX808 micro-plate reader was used to measure light absorbance at 562 nm of the colorimetric marker reaction. The amount of glycogen in the cell extracts was calculated by comparison to a standard series of glycogen solutions. The quantity of glycogen was normalized to the protein content of the sample, which was determined by a Micro BCA™ Protein Assay Kit (Thermo Fischer). The kit was used according to the manufacturer’s instructions and samples were measured as triplicates. Results are presented in units of μg glycogen/μg protein

### Cytochemical PAS staining

Cells were cultured on Matrigel®-coated cover slips (in 6 well plates) and fixed with 4 % PFA for 10 min. Before the samples were incubated with 100 % isopropyl alcohol for 5 min, they were washed three times with PBS. Next the cells were incubated in 0.5 % periodic acid for 5 min, and then washed in distilled water (5 min) and tap water (1 min). After 12 min of incubation in Schiff’s reagent, cells were washed again for 6 min in first distilled water (5min) and then tap water (1 min). Finally the cells were covered with Aquatex® and imaged using an Olympus IX83 microscope in a bright field setup.

### Reproducibility

GLMs that showed the typical time-lapse development with dorsal neural tube epithelium and ventrally mesenchymal LPM, somitic development (successfully underwent gastrulation), followed by dense continuous migration between Day 17 to Day 35 (time-points 1 to 4), as well as were able to generate a dense network and bulge formation beyond Day 56 (time-points 5 to 6) were considered successful. Batches and not individual GLMs were potential source of variability **(Figure 1A,H, 2I,J, 3N)**, namely when GLMs underwent gastrulation, followed by mesoderm segregation skeletal muscle lineage was always present. Thus, we consider a time-point for analysis to be consistent when it was associated with similar morphological changes (Figure 1,2,3) for a minimum of six independent batches during musculoid development for each cell line tested in this study.

Selection criteria, mentioned above, on distinguishing right forming organoids established continuous developmental trajectories similar to *in vivo* fetal development with overlapping lineage representation for all our samples (n=11 scRNAseq samples, n=5 Spatial Transcriptomics samples with more than 100.000 cells analyzed. Similar behavior exhibited n=7 Bulk RNA seq samples and additional n=4 scRNAseq samples described in^77^. In both cases each time-point was from independent derivation. For all following experiments, only successfully formed GLMs were used; these musculoids showed similar results in all experiments. Specifically, IF stainings were repeated with at least three independent musculoids for each line. TEM was performed on three control musculoids for adipogenesis and musculature investigation and on three CD82^+^ skeletal muscle cultures (Figure 4K, S4I) for investigating musculature maturation.

### Data availability

The gene expression datasets generated and analyzed during the current study are available in the Gene Expression Omnibus repository: Single-cell RNA sequencing data have been deposited under accession GSE210069 and Spatial transcriptomics data under accession GSE262361. The following public datasets were used for scRNA-seq analysis: the Mouse E9.5-E10.5 trunk neural crest^107^, mouse forelimb development^108^, mouse satellite cells during regeneration^109^, human gastrulation^9^, human developing embryo post gastrulation^110,111^, human skeletal muscle reference^52,112^ and human limb development^113^. All additional data supporting the findings of this study are available within the article and its Supplementary Information. All other raw data used for plotting in the figures are provided as source data. Source data are provided with this paper.

---

**Figure S1.**
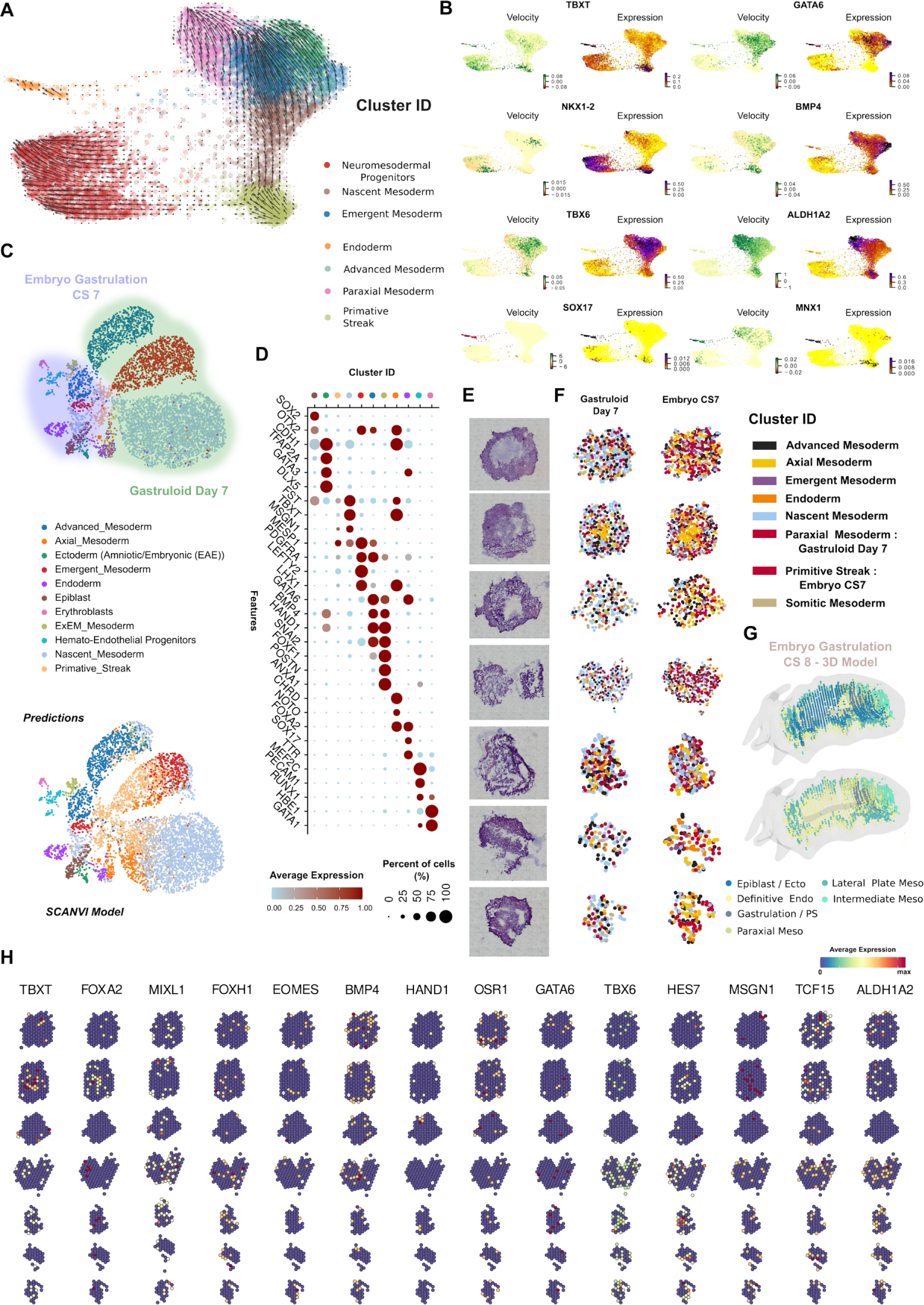
GLM characterization at early stages and comparison to gastrulating human embryo^9^ (Carnegie Stage, CS7). **(A)** RNA velocity analysis on force directed graph embedding indicates from a primitive streak state mesodermal lineage segregation towards lateral plate and paraxial mesodermal trajectories, and presence of an endodermal and neural cluster with pre-neuromesodermal progenitor identity. **(B)** Feature plots indicate RNA velocity dynamics and expression on key markers for each lineage, and highlight BPM4 upregulation for lateral plate mesoderm and Retinoc acid (ALDH1A2) for paraxial mesoderm **(C)** UMAP plots based on a semi-supervised deep learning (SCANVI) approach, trained on human gastrulating embryo dataset, to map population from GLM at gastrulation stage to human gastrulating embryo (CS7) demonstrates *in vitro* reconstruction of mesodermal lineage segregation and dynamics following gastrulation. **(D)** Dot plot illustrating expression of representative mesodermal, endodermal, epiblast and endothelial genes for the characterization of clusters at human gastrulating embryo. The size of each circle reflects the percentage of cells in a cluster where the gene is detected, and the color reflects the average expression level within each cluster (blue, low expression; red, high expression). **(E)** H&E histology pictures of GLMs at Day 5 **(F)** 3D model of human gastrulation, CS8 Carnegie stage, based on spatial transcriptomics, depicts mesodermal lineage segregation following gastrulation^46^. (https://cs8.3dembryo.com/#/model3d/embryo) **(G)** Mapping of single cells from Day 7 GLMs and human CS7 gastrulating embryo to spatial GLM sections from Day 5 indicate similar mesodermal population dynamics on sections from gastrulating /ventral to gastrulation planes. **(H)** Spatial feature plots at Day 5 on primitive streak, mesoendodermal markers, *TBXT, FOXA2, MIXL1, FOXH1, EOMES,,* lateral plate mesodermal, *BMP4, HAND1, OSR1, GATA6,* and paraxial mesodermal, *TBX6, HES7, MSGN1, TCF15, ALDH1A2 markers,* indicate reproducibility between different GLMs at a gastrulating to mesoendodermal ventral section plane.

**Figure S2.**
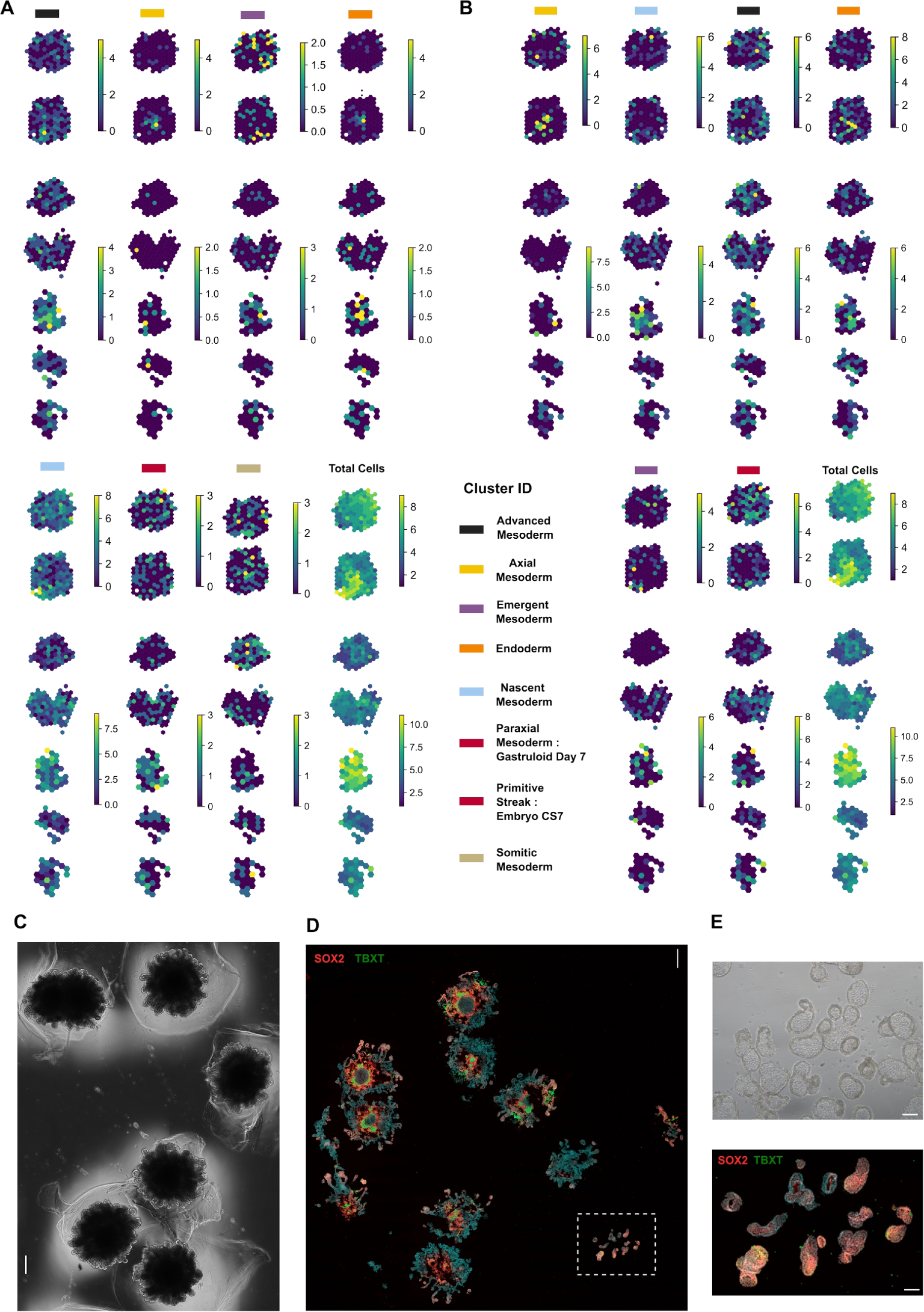
Deciphering spatial organization during GLM development at gastrulation stage via spatial transcriptomics and immunocytochemistry. **(A)** Heatmap illustrating cluster assignment per spot during Mapping of single cells from Day 7 GLMs and human CS7 gastrulating embryo **(B)** to spatial GLM sections from Day 5 indicate spots with overlapping and distinct pattern expression for each cluster. **(C)** Brightfield image at Day 11 illustrating neuro-mesodermal (NMP) mediated body axis elongation from dorsal sites during GLM development upon continuation of the CHIR99021, LDN198189, bFGF and Retinoc acid stimulation following gastrulation at Day 7. **(D)** Immunocytochemistry pictures on TBXT, SOX2 markers indicates gastrulating populations at the core (TBXT^+^) of the structure and NMP progenitors (SOX2^+^/TBXT^+^) at the edge of body axis -like elongating structures. **(E)** Brightfield image illustrating the body axis-like elongating structures together with an immunostaining section image at the level of those structures indicates NMP progenitors (SOX2^+^/TBXT^+^) identity. Scale Bars: 500um in **(C),(D),** 100um in **(E).**

**Figure S3.**
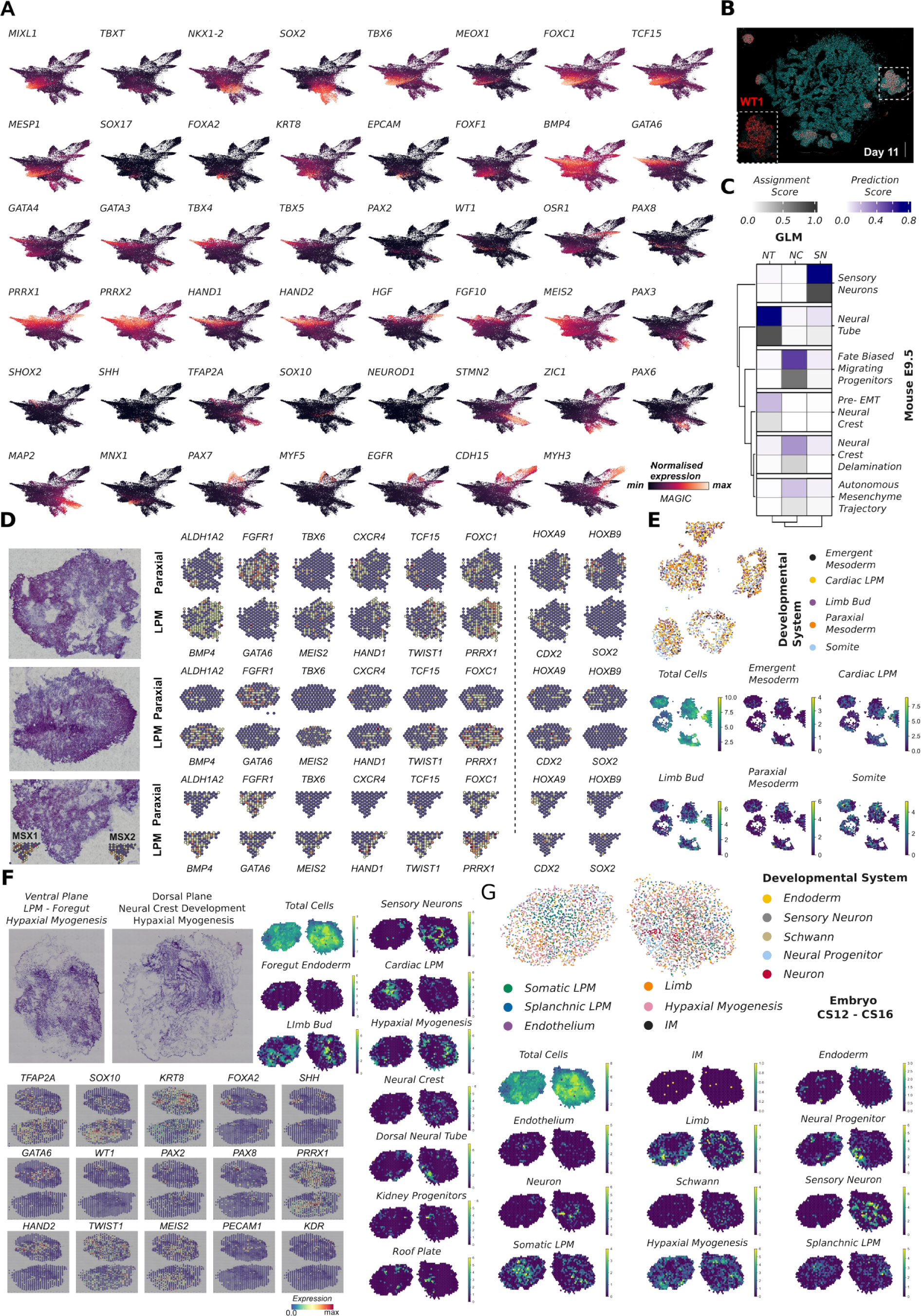
Lineage progression investigation within GLM following gastrulation via spatial transcriptomics and comparison to human developing embryo (Carnegie Stage, CS12-CS16)^111^. **(A)** Feature plots from force-directed k-nearest neighbor graph on 55.778 cells at Day 7, Day 13, Day 19, Day 35, Day 56 and Day 84 illustrate MAGIC imputed average expression levels for gastrulating/mesodermal/endodermal genes, *TBXT, MILX1, MESP1, MEOX1, TBX6, FOXF1,* markers along the skeletal muscle trajectory, *PAX7, MYF5, EGFR, MYH3, CDH15*, endodermal fore-gut markers, *MNX1, KRT8, SOX17, SHH, EPCAM, FOXA2,* cardiac field/limb bud initiation markers *HAND1, HAND2, GAT6, GATA3, GATA4, SHOX2, FGF10, MEIS2,* limb bud mesenchyme markers, *HGF, PRRX1, PRRX2*, and neural tube, *PAX3, PAX6, ZIC1*, neural crest/sensory neurons *TFAP2A, SOX10, STMN2, MAP2*. **(B)** Immunocytochemistry picture at Day 11 indicates presence of WT1^+^ intermediate mesodermal cells at the periphery of GLMs, ventral to gastrulation plane. Scale Bar: **(C)** Heatmap with hierarchical clustering depicts the percentage of prediction and assignment score from GLM cells along the dorsal neural tube/neural crest development with evidence of sensory branch bifurcation from the main migrating NC stream upon unbiased mapping to the *in vivo* counterpart, mouse E9.5 stages. NC: Neural Crest, NT: Neural Tube, SN: Sensory neurons. **(D)** H&E hitology pictures of GLMs at Day 9. Spatial feature plots at Day 9 on lateral plate mesodermal (LPM), *BMP4, GATA6, MEIS2, HAND1, TWIST1, PRRX1,* paraxial(PM)/somitic/dermomyotomal, ALDH1A2*, FGFR1, TBX6, CXCR4, TCF15, FOXC1,* and *HOX genes HOXA9,HOXB9,* in the absence of neural marker SOX2 indicate reproducibility between different GLMs during mesodermal segregation and presence of LPM/PM populations at the same section plane. **(E)** Mapping of single cells from Day 7 GLMs to spatial GLM sections from Day 9 indicate similar mesodermal population dynamics on sections from gastrulating ventral to gastrulation planes. Heatmap illustrating cluster assignment per spot during mapping of single cells from Day 7 GLMs to spatial GLM sections from Day 9 indicate spots with overlapping and distinct pattern expression for each mesodermal cluster. **(F)** H&E hitology pictures of GLMs at Day 19. Spatial feature plots at Day 19 on neural crest, *TFAP2A, SOX10,* fore-gut/endodermal *KRT8, FOXA2, SHH,* intermediate mesodermal*, WT1, PAX2, PAX8,* cardiac field/limb bud*, GATA6, HAND2, TWIST1, MEIS2, PRRX1,* and endothelial, *PECAM1, KDR* markers. Heatmap illustrating the mapping of single cells per spot from Day 19 GLMs to spatial GLM sections from Day 19, indicate presence of a ventral and a dorsal to gastrulation plane with distinct pattern of mesodermal/endodermal and neural clusters retrospectively. **(G)** Mapping of single cells from CS12-CS16 human developing embryo to spatial GLM sections from Day 19 indicate similar spatial pattern to Day 19 GLMs. Heatmap illustrating cluster assignment per spot during Mapping of single cells from CS12-CS16 human developing embryo to spatial GLM sections from Day 19 indicate spots with overlapping and distinct pattern expression for each cluster.

**Figure S4.**
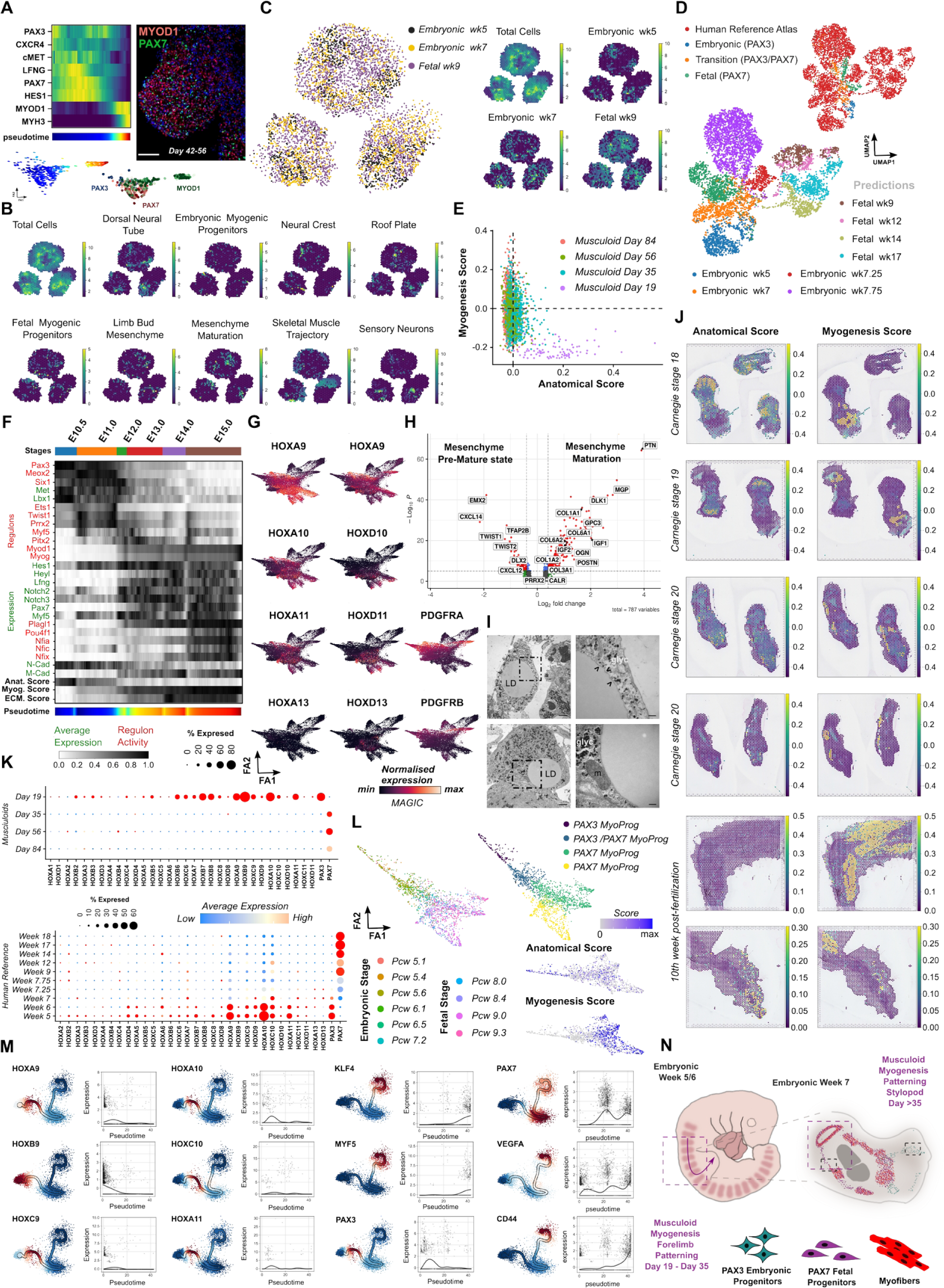
Characterization of embryonic-fetal myogenic progenitors during GLM, mouse and human limb development. **(A)** Pseudotemporal ordering of cells related to embryonic fetal transition revealed gene dynamics on Notch signaling, e.g. HES1, LFNG, upon PAX7 upregulation in myogenic progenitors. GLM picture following embryonic/fetal transition, (> Day 35) highlights absence of committed, PAX7^+^/MYOD1^+^ myogenic progenitors at the periphery and at protrusion sites. Scale Bar: 100uM **(B)** Heatmap illustrating cluster assignment per spot during **m**apping of single cells from Day 35 GLMs to spatial GLM sections from Day 35 indicate spots with overlapping and distinct pattern expression for each cluster. **(C)** Mapping of single cells during embryonic/fetal myogenic transition (embryonic week 5, embryonic/fetal week 7, fetal week 9) from human reference atlas GLM sections from Day 35 indicates spatial distribution of myogenic progenitors with fetal identity most prominent at the periphery of GLM structure (edge of initial matrigel droplet). **(D)** UMAP plots based on semi-supervised deep learning approach to map 5^th^ week GLM derived, PAX3^+^, PAX3^+^/PAX7^+^ and PAX7^+^ myogenic progenitors to the human embryonic - fetal reference atlas predict maturation beyond the embryonic fetal transition stage and indicate the generation of myogenic progenitors with early fetal identity. **(E)** Scatter plot of anatomical – myogenesis score for myogenic progenitors at embryonic (Day19), embryonic/fetal (Day35), fetal (Day56) and late fetal (Day84) stages during GLM development. **(F)** Pseudotemporal ordering of cells during embryonic to fetal mouse myogenic transition (E9.5 – E15.0) reveals gene and GRN dynamics from an embryonic anatomical related mesenchymal state e.g Pax3, Lbx1, Meox2, Twist1, N-Cadherin to a fetal myogenic state e.g Pax7, Nfix, Myf5, Plagl1, M-Cadherin. **(G)** Feature plots from force-directed k-nearest neighbor graph on 55.778 cells at Day 7, Day 13, Day 19, Day 35, Day 56 and Day 84 illustrate MAGIC imputed average expression levels for HOX genes, *HOXA9, HOXB9, HOXA10, HOXD10, HOXA11,HOXD11,* and fibro-adipogenic progenitor markers, *PDGFRA, PDGFRB,* indicate forelimb bud identity for mesenchymal clusters **(H)** Differential expression analysis between mesenchymal clusters during 5^th^ week of musculoid development indicates mesenchyme maturation towards a chondrogenic/osteogenic state. **(I)** Pictures at ultra-structural level highlight developing lipid droplet, derived from fibro-adipogenic progenitors, covered by one leaflet of rER membrane during pre-adipocyte formation. Mitochondria are located in close contact to lipid droplet. LD: Lipid Droplet, gly:glycogen, m: mitochondria, arrows: rER Scale Bar: 1uM, 200nM in magnification. **(J)** Spatial feature plots during human hindlimb spatiotemporal development^113^, Carnegie Stages 18,19,20 and 10^th^ week post-fertilization, indicate absence of anatomical score based on HOX gene clusters during on going myogenesis, thus signaling the growth phase of the embryo. **(K)** Dot plot showing the expression of HOX genes across GLM and human embryonic-fetal development. The size of each circle reflects the percentage of cells in a cluster where the gene is detected, and the color reflects the average expression level within each cluster (blue, low expression; red, high expression). **(L)** Single cell expression profiling depicting embryonic fetal myogenic transition, derived from different dataset^113^ and technology (10x Genomics vs Drop-Seq in^52^), indicates similar dynamic with axial anatomical identity for embryonic myogenic progenitors and myogenic core program upregulation followed by downregulation of the axial anatomical program for fetal myogenic progenitors. **(M)** Curved trajectory analysis highlights significant upregulation on HOXA9, HOXB9, HOXC9, during somitic/hypaxial myogenesis stages, followed by significant upregulation on HOXA10, HOXC10, HOXA11 genes at hypaxial myogenesis, *PAX3^+^*.stage followed by upregulation of the core late fetal program, *PAX7, MYF5, KLF4, CD44,* during GLM development. **(N)** Scheme illustrating musculoid patterning during early stages and its correlation to human fetal development.

**Figure S5.**
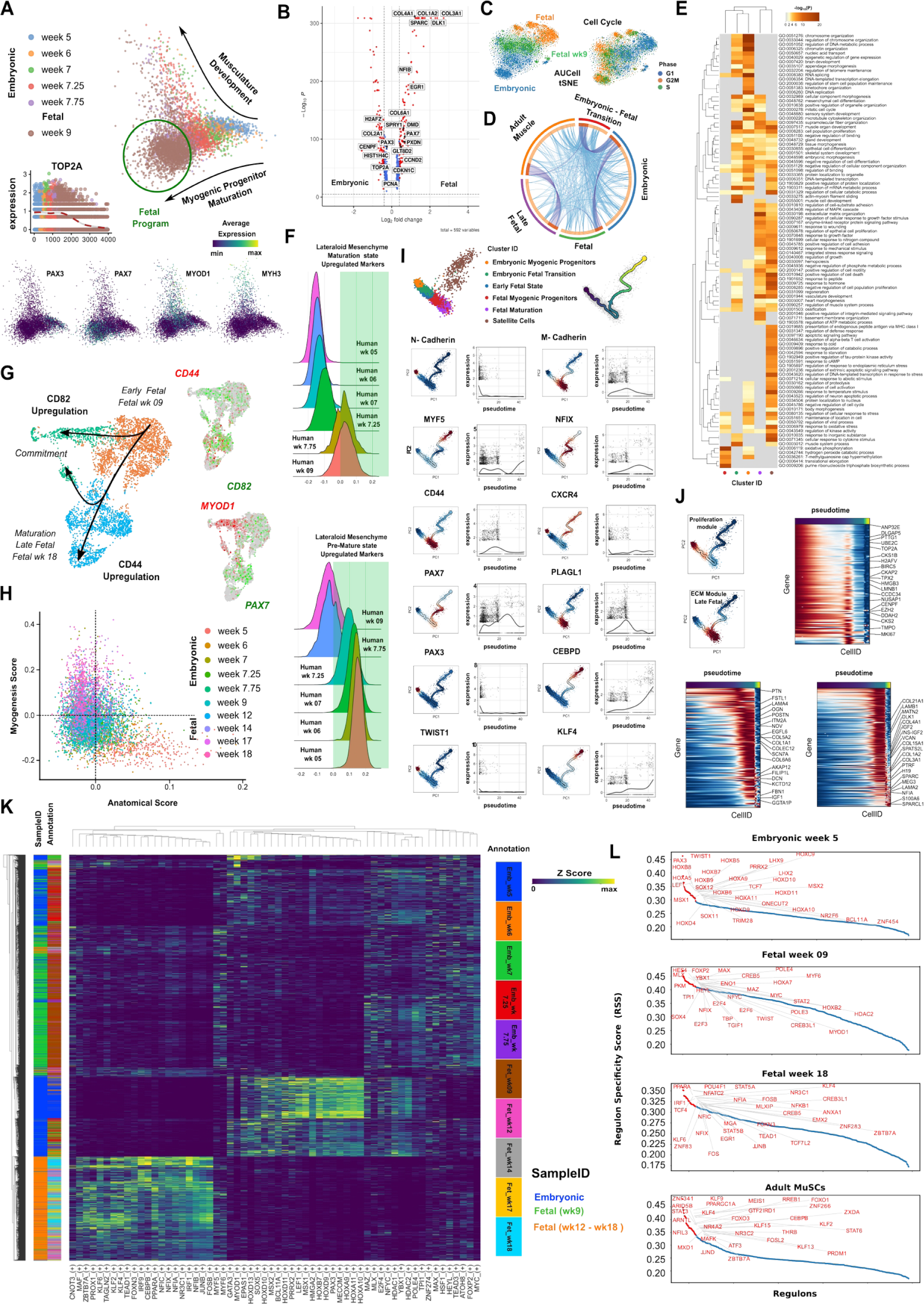
Developmental trajectory of human myogenic progenitors at embryonic, fetal and postnatal stages. **(A)** Datasets from 5^th^ to 9^th^ of human skeletal muscle reference map^52^ depict embryonic fetal myogenic transition on PCA embedding. Feature plots on embryonic (PAX3^+^), fetal (PAX7^+^) and committed (MYH3, MYOD1) markers, indicate developmental trajectories associated with maturation of myogenic progenitors and musculature development during embryonic fetal transition. Pseudotemporal ordering of myogenic progenitors cells highlights downregulation of mitotic marker, *TOP2A*, during embryonic fetal transition. **(B)** Differential expression analysis between embryonic and fetal stages highlights downregulation of mitotic markers, up-regulation of ECM proteins and cell cycle inhibitors on fetal myogenic progenitors and upregulation of cell cycle related genes during embryonic stages. **(C)** Aucell TSNE plot based on active gene regulatory networks highlights cell cycle stages during embryonic fetal reference myogenic map. **(D)** Overlap between gene lists at the gene level, where purple curves link identical genes; including the shared term level, where blue curves link genes that belong to the same enriched ontology term. The inner circle represents gene lists, where hits are arranged along the arc. Genes that hit multiple lists are colored in dark orange, and genes unique to a list are shown in light orange. **(E)** Heatmap of enriched terms across input gene lists for each stage during human skeletal muscle lineage fetal development colored by p-values. **(F)** Ridge plot of developmental score distribution of mesenchyme progenitors across *in vivo* or *in vitro* stages based on the difference between up-regulated markers of GLM mesenchyme progenitors at mature and premature stages, indicate similar developmental pattern within GLMs and m human reference atlas, (Supplementary Table 5). **(G)** UMAP together with feature plots during fetal maturation (fetal week 09-18) indicates CD44,CD82 upregulation during fetal maturation on myogenic progenitors, while MYOD1,CD82 regulation leads to commitment and differentiation. **(H)** Scatter plot of anatomical – myogenesis score for myogenic progenitors during embryonic, fetal human skeletal muscle development. **(I)** Curved trajectory analysis on PCA space for human skeletal muscle reference atlas. Pseudotime was calculated, by learning a principal curve on the 2 first PC components using the ElPiGraph algorithm. Significantly changing genes at each stage, embryonic, fetal, postnatal, for myogenic progenitors along the human myogenic trajectory. **(J)** Curved trajectory analysis on PCA space for human skeletal muscle reference atlas. Pseudotime was calculated, by learning a principal curve on the 2 first PC components using the ElPiGraph algorithm. Clustering based on significantly changing features along pseudotime highlights presence of a proliferation module at embryonic stages followed by an ECM module following embryonic to fetal transition during late fetal stages. **(K)** Heatmap on gene regulatory networks from embryonic (5wk - 7.75wk) and fetal (9wk - 18wk) stages, highlights HOX genes and PAX3 expression during embryonic and NFIX, NFIC, NFIA, NFIB and KLF4 expression during fetal stages. **(L)** Regulon score specificity (RSS) at human reference skeletal muscle developmental trajectory highlights HOX GRNs upregulation at embryonic stages and myogenic related GRN, NFIX,NFIA,NFIB, NFIC, KLF4, FOXO3, CEPBD, at fetal postnatal stages.

**Figure S6.**
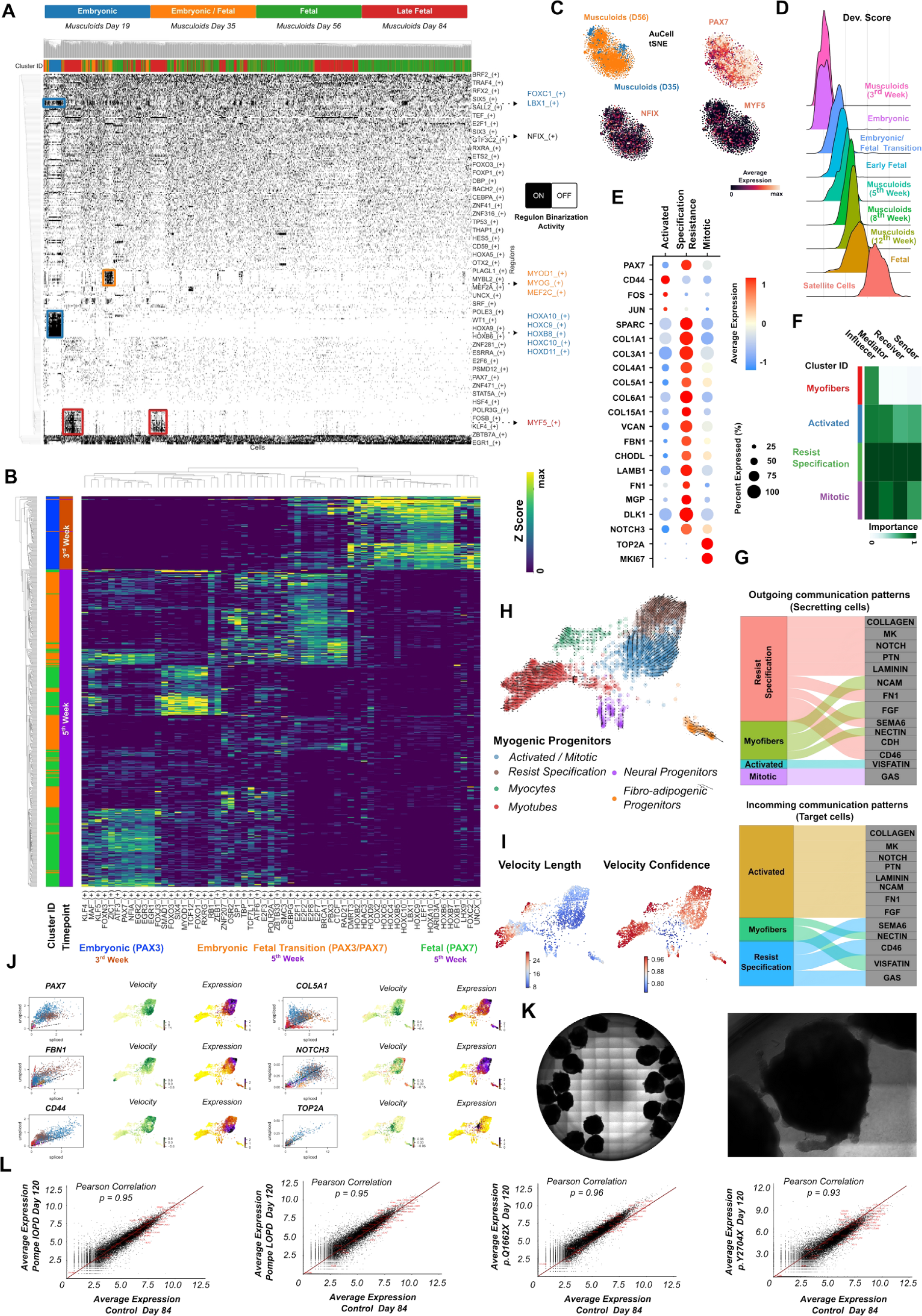
Investigating gene regulatory networks via SCENIC pipeline during GLM development and human reference skeletal muscle atlas. **(A)** Binary activity regulon matrix during musculoid skeletal muscle developmental trajectory. Cluster labels correspond to datasets from each stage (Day 19, Day 35, Day 56, Day 84). Master regulators are color matched with the cell types they control. **(B)** Heatmap on gene regulatory networks highlighting HOX gene expression on 3^rd^ week embryonic clusters, cell cycle regulation during 5^th^ week and during embryonic (PAX3^+^) to fetal (PAX7^+^) transition up-regulation of PAX7/NFIA/KLF4/FOXO3 on 5^th^ week PAX7^+^ fetal clusters together within the EGR family. **(C)** Aucell tSNE plot based on gene regulatory networks depicts further up-regulation on NFIX, PAX7, MYF5 fetal transcription factor between 5^th,^ and 8^th^ week musculoid derived myogenic progenitors. **(D)** Ridge plot of developmental score distribution of myogenic progenitors during musculoid development and across *in vivo* or *in vitro* stages based on the difference between up-regulated satellite cell and embryonic markers from human reference myogenic atlases. **(E)** Dot plot illustrating expression of specific genes for the activated, resisting specification, and a mitotic cluster of 12^th^ week musculoid derived myogenic progenitors. The size of each circle reflects the percentage of cells in a cluster where the gene is detected, and the color reflects the average expression level within each cluster (blue, low expression; red, high expression). **(F)** Heatmap shows the relative importance of each cell group based on the computed four network centrality measures of the Notch signaling network. **(G)** The inferred outgoing and incoming communication patterns of secreting cells, show the correspondence between the inferred latent patterns and cell groups, as well as signaling pathways. The thickness of the flow indicates the contribution of the cell group or signaling pathway to each latent pattern. **(H)** RNA velocity UMAP plot on 12^th^ week GLM dataset highlights the pool of activated myogenic progenitors along a trajectory of myogenic specification resistance and another that leads to myogenic commitment and myotube formation. **(I)** Myogenic progenitor cluster further characterized by the low rate of differentiation (Velocity length) in comparison to myotube cluster that is characterized by high rate of differentiation, while both clusters show high confidence among the cells. **(J)** Feature and scatter plots depicting RNA velocity and gene expression on selected markers representing the activated, CD44, resist specification, COL5A1, FBN1, and mitotic, TOP2A, state of myogenic progenitors. **(K)** Brightfield images depicting morphology of mature developed musculoids with presence of the characteristic protrusions-like structures, where myogenic progenitor maturation occurs **(L)** Scatter plot depicting average expression of genes between control GLM derived myogenic progenitors, Day84, and GLM derived myogenic progenitors from FLNC and Pompe patients lines at GLM level (left, Day 120). Selected genes from the late fetal, adult MuSCs core program, *NFIX, MYF5, PAX7,* and genes related to specification resistance, *NOTCH, ECM,* are highlighted in red.

**Figure S7.**
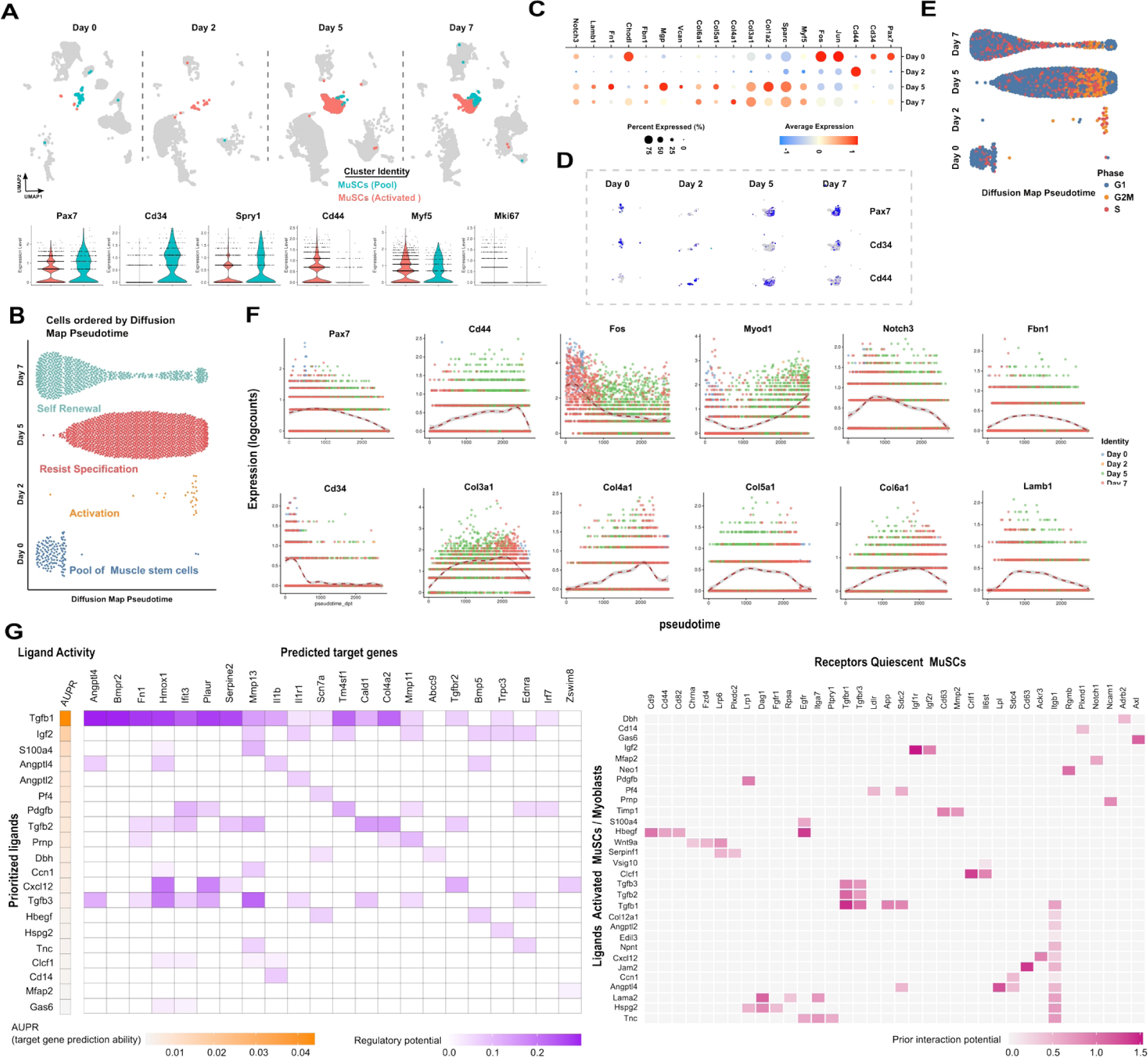
Evaluating adult mouse muscle stem cells at different stages during regeneration at single cell resolution ^109^. **(A)** UMAP plot depicting muscle stem cell populations at day 0, day 2, day 5 and day 7 together with violin plots on activated, CD44, MYF5 and quiescent markers e.g. PAX7, CD34 and SPRY1. **(B)** Ordering of cells by diffusion map pseudotime demonstrates distinct states of muscle stem cells developmental trajectory during muscle regeneration and highlights the self renewal process. **(C)** Dot plot showing the expression of extracellular matrix genes, dormant genes e.g. Pax7, Cd34, activated genes e.g. Cd44, Jun and Fos during muscle stem cell regeneration process. The size of each circle reflects the percentage of cells in a cluster where the gene is detected, and the color reflects the average expression level within each cluster (blue, low expression; red, high expression). **(D)** UMAP plots on Pax7, Cd34 and Cd44 distinguish activated, quiescent and self renewal populations during muscle regeneration. **(E)** Diffusion map pseudotime of muscle stem cells and cell cycle state during each stage. **(F)** Pseudotemporal ordering of muscle stem cells progenitors from day 0, day 2, day 5 and day 7 depicting gene expression trend for extracellular matrix genes, Pax7, Cd34, and Cd44, Jun, Fos along diffusion map pseudotime. **(G)** Outcome of NicheNet’s ligand activity prediction for the 20 ligands best predicting the quiescent gene set, Nichenet analysis performed on differentially expressed genes between Day 7 (return to quiescence for MuSCs, receiver cell type: Quiescent MuSCs) and Day 5 (activation /transient amplification of MuSCs, sender/niche cell type: Activated/Committed MuSCs), better predictive ligands are ranked higher. Heatmap to infer receptors and top-predicted target genes of ligands that are top-ranked in the ligand activity NicheNet’s analysis (left). Ligand-receptor network inference for top-ranked ligands between activated/committed MuSCs ligands and receptors strongly expressed in quiescent MuSCs cells (right).

**Figure S8.**
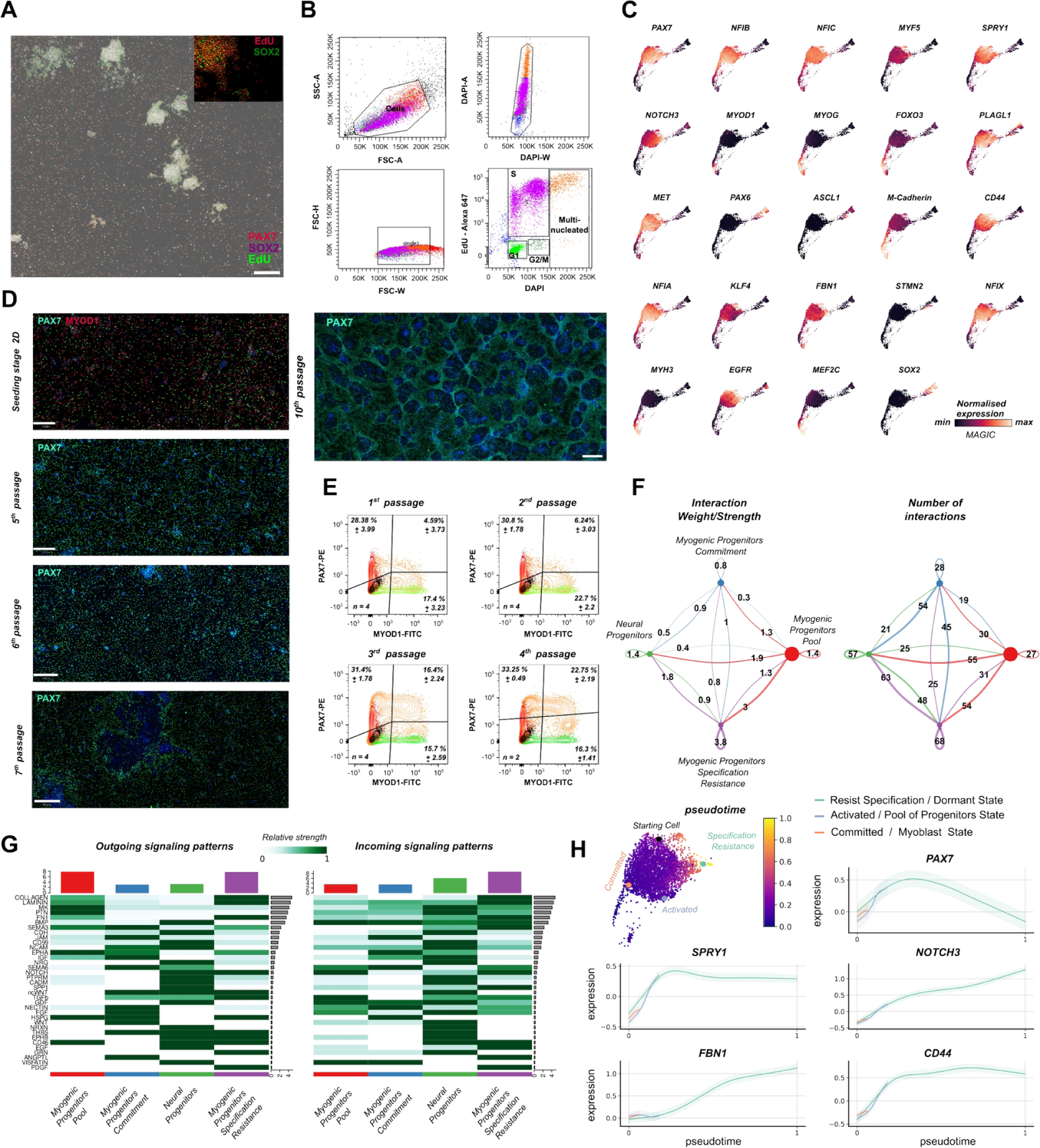
Characterization and *in vitro* sustainable expansion of late fetal myogenic progenitors. **(A)** Immunocytochemistry images indicate proliferating, EdU^+^/PAX7^+^, and dormant, EdU^−-^/PAX7^+^, fetal myogenic progenitors, and proliferating neural progenitors (EdU^+^). **(B)** Density plots indicating the FACS gating during cell cycle analysis of fetal myogenic progenitors derived via GLM and expanded in 2D. Plot investigating area vs width for DAPI signal indicates presence of multi-nucleated proliferating cells in the culture. Before analysis cells incubated with 5uM EdU overnight (18hr). Colors correspond to the same cell population among the plots**. (C)** Feature plots from force-directed k-nearest neighbor graph on fetal myogenic progenitors at Day 180, derived via GLM approach (Day56-Day84) and expanded in 2D for 10 passages, illustrate MAGIC imputed average expression levels for core late fetal muscle stem cell program*, PAX7, MYF5, NOTCH3, KLF4, PLAGL1, NFIX, NFIA, NFIB, NFIC, CD44,* absence of committed markers, *MYOD1, MYOG, MEF2C*. **(D)** Immunocytochemistry images from late fetal myogenic progenitors, derived via GLM and expanded in 2D, at different passages indicate at the seeding stage an initial presence of committed, MYOD1^+^/PAX7^−^, and uncommitted, MYOD1^−^/PAX7^+^, cell populations, followed by sustainable propagation of uncommitted myogenic progenitors for at least 10 passages (> 100 Days). **(E)** Contour plots illustrating FACS quantification on MYOD1^+^ and PAX7^+^ fetal myogenic progenitors established at GLM level (Day56 - Day84) till late fetal stage, followed by 2D expansion. Black contour: unstained population, green contour: MYOD1 FMO control, red contour: PAX7 FMO control. n=number of samples and quantification represents mean values ± standard deviation. **(F)** The inferred outgoing and incoming communication patterns of secreting cells, show the correspondence between the inferred latent patterns and cell groups, as well as signaling pathways. The thickness of the flow indicates the contribution of the cell group or signaling pathway to each latent pattern. **(G)** Heatmap indicating signals contributing most to the outgoing or incoming signalling of myogenic progenitor and neural clusters in datasets of expanded in 2D for 10 passages myogenic progenitors. **(H)** Pseudotime analysis on myogenic progenitors (GLM derived, 2D expansion for 10 passages) using Palantir pipeline. Gene expression trends for *SPRY1*, *CD44, FBN1, NOTCH3,* highlight upregulation on their expression along pseudotime. Trends are colored based on states/cells presented in pseudotime illustration (circles). Initial (black circle) and activated (magenta circle), resist specification (cyan circle), committed (orange circle) state were calculated/selected based on RNA velocity analysis presented in Fig. 4c. Shaded region represents 1 s.d. Scale Bars: 500um in **(C)**,**(D,** 10^th^ passage), 400um in **(D** 6^th^,7^th^ passage), 250um in **(D,** seeding, 5^th^ passage).

**Figure S9.**
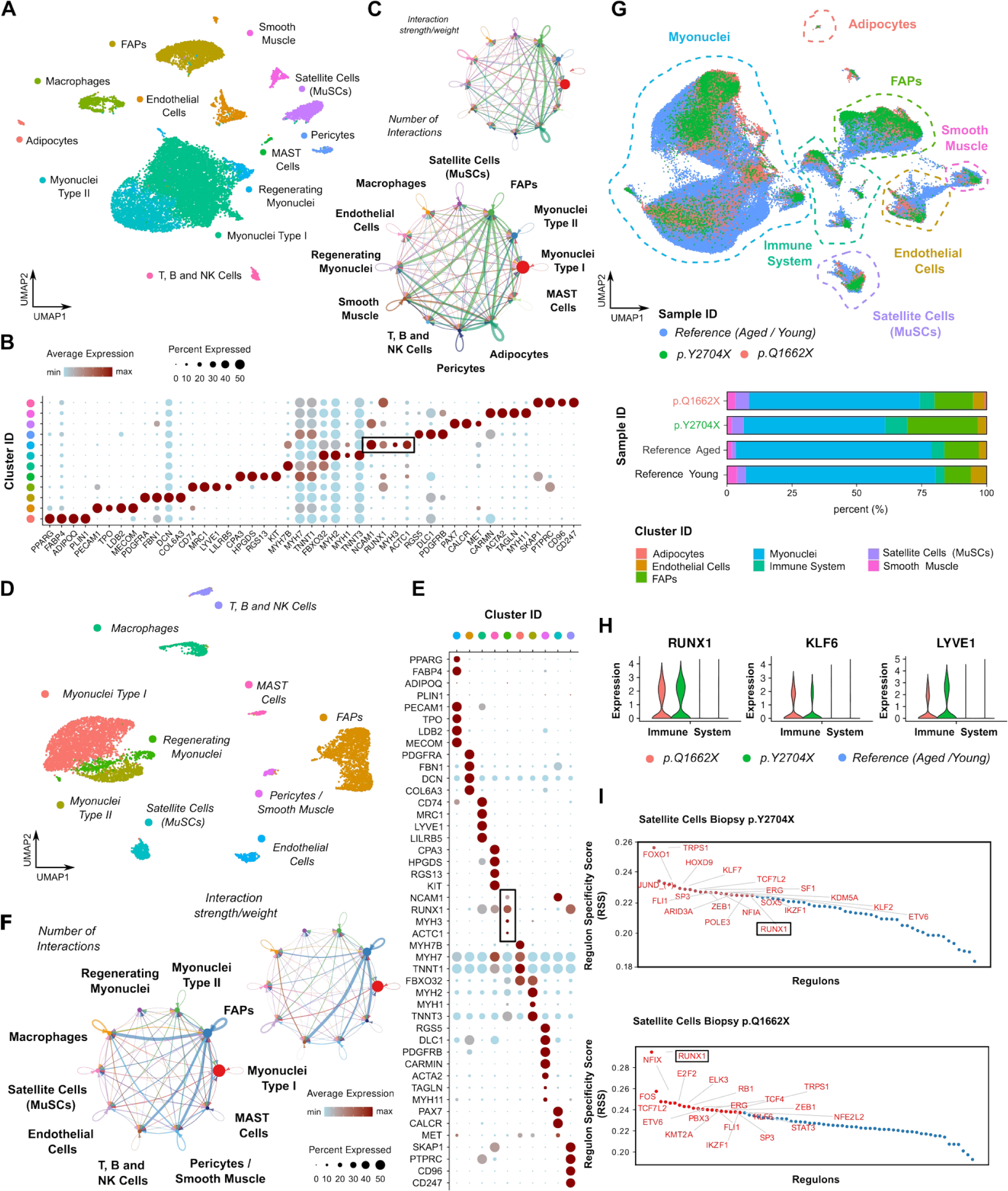
Characterization of patient derived biopsies (FLNC p.Q1662X, FLNC p.Y2704X) at single cell resolution. **(A)** UMAP analysis on FLNC p.Q1662X Biopsy depicting cluster composition, together with **(B)** dot plot with representative markers for each cluster. The size of each circle reflects the percentage of cells in a cluster where the gene is detected, and the color reflects the average expression level within each cluster (blue, low expression; red, high expression). **(C)** Aggregated cell-cell communication network indicating the number of significant ligand-receptor pairs between any pair of two cell populations. The edge width is proportional to number of ligand-receptor pairs. **(D)** UMAP analysis on FLNC p.Y2704X Biopsy depicting cluster composition, together with **(E)** dot plot with representative markers for each cluster. The size of each circle reflects the percentage of cells in a cluster where the gene is detected, and the color reflects the average expression level within each cluster (blue, low expression; red, high expression). **(F)** Aggregated cell-cell communication network indicating the number of significant ligand-receptor pairs between any pair of two cell populations. The edge width is proportional to number of ligand-receptor pairs. **(G)** Integrative comparison between FLNC p.Q1662X, FLNC p.Y2704X biopsies and skeletal muscle reference atlas composed from young (19-22Yr) and aged (70-76Yr) datasets, indicate via the increased adult muscle stem cells and immune system representation within FLNC biopsies, signs of ongoing regeneration. FLNC p.Y2704X biopsy further exhibits increased fibrosis, a sign of disease progression. **(H)** Violin plots depicting expression of RUNX1, KLF6, LYVE1 markers on immune related clusters from FLNC p.Q1662X, FLNC p.Y2704X biopsies and those from young (19-22Yr) and aged (70-76Yr) skeletal muscle reference atlas, indicate ongoing immune response on FLNC patient derived biopsies. **(I)** Regulon score specificity (RSS) at human skeletal muscle biopsies highlights GRNs from nuclear factor I family, NFIX, NFIA and GRN involved in MuSCs activation, FOS, JUN, RUNX1.

**Figure S10.**
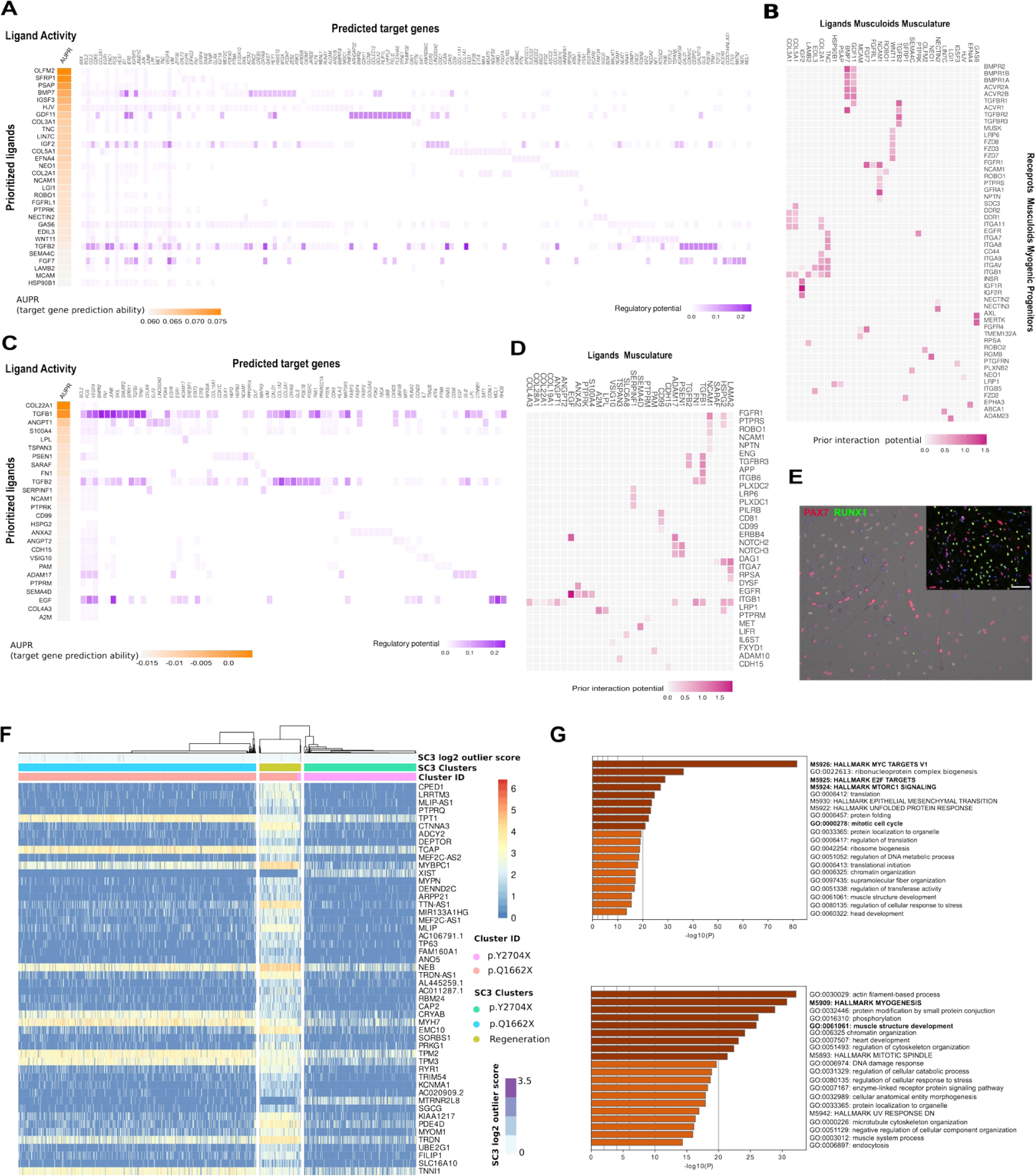
Characterization of control and patient derived fetal and adult MuSCs (FLNC p.Q1662X, FLNC p.Y2704X) at single cell resolution. **(A)** Outcome of NicheNet’s ligand activity prediction for the 20 ligands best predicting the GLM derived fetal myogenic progenitors niche gene set, Nichenet analysis performed on deferentially expressed genes between FLNC p.Q1662X, FLNC p.Y2704X (receiver cell type: Myogenic progenitors) and control GLM derived musculoid datasets (sender/niche cell type: musculature), better predictive ligands are ranked higher. Heatmap to infer receptors and top-predicted target genes of ligands that are top-ranked in the ligand activity NicheNet’s analysis **(B)** Ligand-receptor network inference for top-ranked ligands between myotube/myofiber ligands and receptors strongly expressed in myogenic progenitors. **(C)** Outcome of NicheNet’s ligand activity prediction for the 20 ligands best predicting the adult MuSCs niche gene set, Nichenet analysis performed on deferentially expressed genes between FLNC p.Q1662X, FLNC p.Y2704X (receiver cell type: MuSCs, satellite cells) and control MuSCs (sender/niche cell type: musculature), better predictive ligands are ranked higher. Heatmap to infer receptors and top-predicted target genes of ligands that are top-ranked in the ligand activity NicheNet’s analysis **(D)** Ligand-receptor network inference for top-ranked ligands between myofiber associated ligands and receptors strongly expressed in adult MuSCs. **(E)** Immunocytochemistry images from FLNC biopsies and musculoid derived myogenic progenitors indicate committed, RUNX1^+^/PAX7^+^, uncommitted RUNX1^−^/PAX7^+^, myogenic progenitors. Brightfield plane indicates presence of RUNX1^+^/PAX7^−^ nuclei at myoblast/myotubes. Scale Bar: 100uM **(F)** Differential expression analysis and SC3 clustering for Biopsy derived MuSCs (FLNC p.Q1662X, FLNC p.Y2704X) datasets, indicate committed and regeneration state predominantly for p.Q1662X MuSCs. **(G)** Selected enriched GO terms from DEGs enriched in p.Q1662X, p.Y2704X FLNC adult MuSCs versus GLM derived p.Q1662X, p.Y2704X FLNC myogenic progenitors (top). Selected enriched GO terms from DEGs enriched in GLM derived p.Q1662X, p.Y2704X FLNC myogenic progenitors versus p.Q1662X, p.Y2704X FLNC adult MuSCs (bottom).

**Figure S11.**
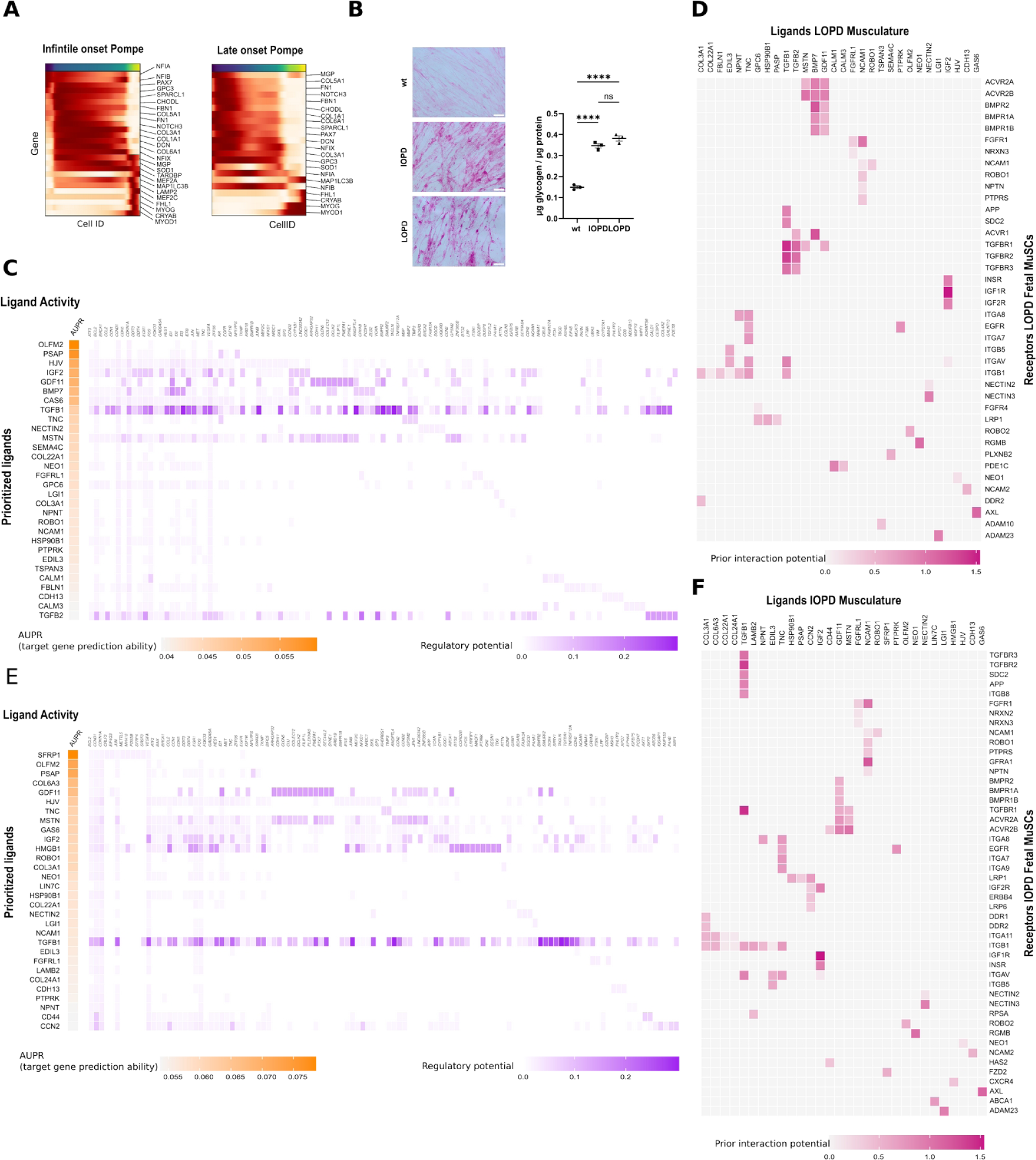
Profiling GLM derived myogenic progenitors from infantile and late onset Pompe lines at single cell resolution. **(A)** Curved trajectory analysis on PCA space for human skeletal muscle reference atlas. Pseudotime was calculated, by learning a principal curve on the 2 first PC components using the ElPiGraph algorithm. Significantly changing genes at each stage, embryonic, fetal, postnatal, for myogenic progenitors along the human myogenic trajectory. **(B)** Cytochemical PAS staining and glycogen assay on control and IOPD, LOPD myotubes following differentiation indicates significant glycogen accumulation in IOPD, LOPD myotubes compared to control line. Statistical analysis was performed by a one-way ANOVA and showed a p-value <0.0001 for the comparison of wt with IOPD/LOPD. Scale Bar: 50 μm. n=4, Statistics: *p<0.05, **p<0.01, ***p<0.001, ****p<0.0001, ns: not significant. **(C)** Outcome of NicheNet’s ligand activity prediction for the 20 ligands best predicting the GLM derived fetal myogenic progenitors niche gene set, Nichenet analysis performed on deferentially expressed genes between late onset Pompe (receiver cell type: Myogenic progenitors) and control GLM derived musculoid datasets (sender/niche cell type: musculature), better predictive ligands are ranked higher. Heatmap to infer receptors and top-predicted target genes of ligands that are top-ranked in the ligand activity NicheNet’s analysis **(D)** Ligand-receptor network inference for top-ranked ligands between myotube/myofiber ligands and receptors strongly expressed in myogenic progenitors. **(E)** Outcome of NicheNet’s ligand activity prediction for the 20 ligands best predicting the GLM derived fetal myogenic progenitors niche gene set, Nichenet analysis performed on deferentially expressed genes between infantile onset Pompe (receiver cell type: Myogenic progenitors) and control GLM derived musculoid datasets (sender/niche cell type: musculature), better predictive ligands are ranked higher. Heatmap to infer receptors and top-predicted target genes of ligands that are top-ranked in the ligand activity NicheNet’s analysis **(F)** Ligand-receptor network inference for top-ranked ligands between myotube/myofiber ligands and receptors strongly expressed in myogenic progenitors.

**Figure S12.**
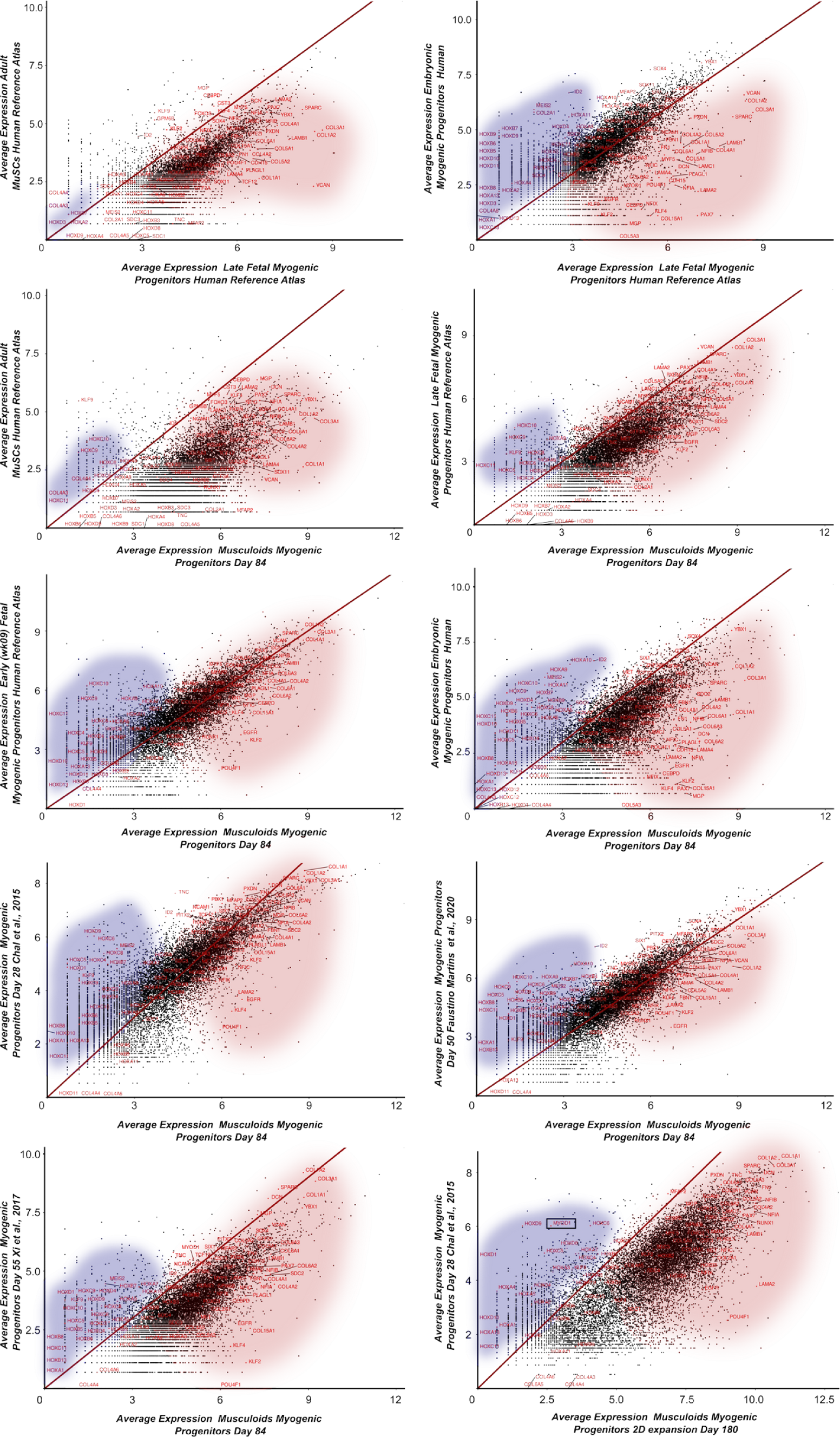
Profiling myogenic progenitors derived via GLM and 2D or 3D *in vitro* differentiation approaches. **(A)** Scatter plots illustrating “pseudobulk” profiles derived following integration of single cell RNA seq data using the `AggregateExpression` command from Seurat. Gene expression profile comparison between adult, fetal and embryonic stages of myogenic progenitors during human development, between GLM derived and adult, fetal and embryonic *in vivo* myogenic progenitors, and between GLM derived myogenic progenitors and those derived from the state of the art 2D or 3D *in vitro* differentiation approaches. Red circle indicates ECM and fetal/adult muscle stem cells related genes. Blue circle indicates Hox gene expression.

## References

1 Nieuwkoop, P. D. (1952). Activation and organization of the central nervous system in amphibians. Part III. Synthesis of a new working hypothesis. J. Exp. Zool. 120, 83–108.

2 Nieuwkoop, P. D. and Nigtevecht, G. V. (1954). Neural activation and transformation in explants of competent ectoderm under the influence of fragments of anterior notochord in *Urodeles*. J. Embryol. Exp. Morphol. 2, 175–193.

3 Tzouanacou E, Wegener A, Wymeersch FJ, Wilson V, Nicolas JF. (2009). Redefining the progression of lineage segregations during mammalian embryogenesis by clonal analysis. Dev Cell. 17, 365–376.

4 Brown JM, Storey KG. (2000). A region of the vertebrate neural plate in which neighbouring cells can adopt neural or epidermal fates. Curr Biol. 10, 869–872.

5 Iimura, T., Pourquié, O. (2006). Collinear activation of *Hoxb* genes during gastrulation is linked to mesoderm cell ingression. Nature 442, 568–571.

6 Cambray, N., Wilson, V. (2007). Two distinct sources for a population of maturing axial progenitors. Development 134, 2829–2840.

7 Olivera-Martinez, I., Harada, H., Halley, P. A., Storey, K. G. (2012). Loss of FGF-dependent mesoderm identity and rise of endogenous retinoid signalling determine cessation of body axis elongation. PLoS biology, 10, e1001415.

8 Henrique, D., Abranches, E., Verrier, L., Storey, K. G. (2015). Neuromesodermal progenitors and the making of the spinal cord. Development, 142, 2864–2875.

9 Tyser, R. C. V., Mahammadov, E., Nakanoh, S., Vallier, L., Scialdone, A., Srinivas, S. (2021). Single-cell transcriptomic characterization of a gastrulating human embryo. Nature 600, 285–289.

10 Prummel, K.D., Nieuwenhuize, S., Mosimann, C. (2020). The lateral plate mesoderm. Development 147, dev175059.

11 Begemann G., Schilling T.F., Rauch G.J., Geisler R., Ingham P.W. (2001). The zebrafish neckless mutation reveals a requirement for raldh2 in mesodermal signals that pattern the hindbrain. Development 128, 3081–3094.

12 Grandel, H., Lun, K., Rauch, G. J., Rhinn, M., Piotrowski, T., Houart, C., Sordino, P., Küchler, A. M., Schulte-Merker, S., Geisler, R., Holder, N., Wilson, S. W., Brand, M. (2002). Retinoic acid signalling in the zebrafish embryo is necessary during pre-segmentation stages to pattern the anterior-posterior axis of the CNS and to induce a pectoral fin bud. Development 129, 2851–2865.

13 Gibert Y., Gajewski A., Meyer A., Begemann G. (2006). Induction and prepatterning of the zebrafish pectoral fin bud requires axial retinoic acid signaling. Development 133, 2649–2659.

14 Niederreither K., Subbarayan V., Dollé P., Chambon P. (1999). Embryonic retinoic acid synthesis is essential for early mouse post-implantation development. Nat. Genet. 21, 444–448.

15 Stratford T., Horton C., Maden M. (1996).Retinoic acid is required for the initiation of outgrowth in the chick limb bud. Curr. Biol. 6, 1124–1133.

16 Nishimoto, S., Wilde, S. M., Wood, S.,Logan, M. P. (2015). RA Acts in a Coherent Feed-Forward Mechanism with Tbx5 to Control Limb Bud Induction and Initiation. Cell reports, 12, 879–891.

17 van den Brink, S. C., Baillie-Johnson, P., Balayo, T., Hadjantonakis, A. K., Nowotschin, S., Turner, D. A., Martinez Arias, A. (2014). Symmetry breaking, germ layer specification and axial organisation in aggregates of mouse embryonic stem cells. Development 141, 4231–4242.

18 van den Brink, S. C., Alemany, A., van Batenburg, V., Moris, N., Blotenburg, M., Vivié, J., Baillie-Johnson, P., Nichols, J., Sonnen, K. F., Martinez Arias, A., van Oudenaarden, A. (2020). Single-cell and spatial transcriptomics reveal somitogenesis in gastruloids. Nature 582, 405–409.

19 Turner, D. A., Girgin, M., Alonso-Crisostomo, L., Trivedi, V., Baillie-Johnson, P., Glodowski, C. R., Hayward, P. C., Collignon, J., Gustavsen, C., Serup, P., Steventon, B., P Lutolf, M., Arias, A. M. (2017). Anteroposterior polarity and elongation in the absence of extra-embryonic tissues and of spatially localised signalling in gastruloids: mammalian embryonic organoids. Development 144, 3894–3906.

20 Moris, N., Anlas, K., van den Brink, S. C., Alemany, A., Schröder, J., Ghimire, S., Balayo, T., van Oudenaarden, A., Martinez Arias, A.(2020). An in vitro model of early anteroposterior organization during human development. Nature 582, 410–415.

21 Veenvliet, J. V., Bolondi, A., Kretzmer, H., Haut, L., Scholze-Wittler, M., Schifferl, D., Koch, F., Guignard, L., Kumar, A. S., Pustet, M., Heimann, S., Buschow, R., Wittler, L., Timmermann, B., Meissner, A., Herrmann, B. G.(2020). Mouse embryonic stem cells self-organize into trunk-like structures with neural tube and somites. Science, 370, eaba4937.

22 Girgin, M. U., Broguiere, N., Hoehnel, S., Brandenberg, N., Mercier, B., Arias, A. M., Lutolf, M. P.(2021). Bioengineered embryoids mimic post-implantation development in vitro. Nat Commun 12, 5140.

23 Olmsted, Z.T., Paluh, J.L. (2021). Co-development of central and peripheral neurons with trunk mesendoderm in human elongating multi-lineage organized gastruloids. Nat Commun 12, 3020.

24 Hashmi, A., Tlili, S., Perrin, P., Lowndes, M., Peradziryi, H., Brickman, J. M., Martínez Arias, A., Lenne, P. F.(2022). Cell-state transitions and collective cell movement generate an endoderm-like region in gastruloids. eLife, 11, e59371.

25 Libby, A. R. G., Joy, D. A., Elder, N. H., Bulger, E. A., Krakora, M. Z., Gaylord, E. A., Mendoza-Camacho, F., Butts, J. C., McDevitt, T. C.(2021). Axial elongation of caudalized human organoids mimics aspects of neural tube development. Development, 148, dev198275.

26 Gouti, M., Tsakiridis, A., Wymeersch, F. J., Huang, Y., Kleinjung, J., Wilson, V., Briscoe, J. (2014). In vitro generation of neuromesodermal progenitors reveals distinct roles for wnt signalling in the specification of spinal cord and paraxial mesoderm identity. PLoS biology 12, e1001937.

27 Tsakiridis, A., Huang, Y., Blin, G., Skylaki, S., Wymeersch, F., Osorno, R., Economou, C., Karagianni, E., Zhao, S., Lowell, S., Wilson, V.(2014). Distinct Wnt-driven primitive streak-like populations reflect in vivo lineage precursors. Development 141, 1209–1221.

28 Turner, D. A., Hayward, P. C., Baillie-Johnson, P., Rué, P., Broome, R., Faunes, F., Martinez Arias, A. (2014). Wnt/β-catenin and FGF signalling direct the specification and maintenance of a neuromesodermal axial progenitor in ensembles of mouse embryonic stem cells. Development 141, 4243–4253.

29 Denham, M., Hasegawa, K., Menheniott, T., Rollo, B., Zhang, D., Hough, S., Alshawaf, A., Febbraro, F., Ighaniyan, S., Leung, J., Elliott, D. A., Newgreen, D. F., Pera, M. F., Dottori, M. (2015). Multipotent caudal neural progenitors derived from human pluripotent stem cells that give rise to lineages of the central and peripheral nervous system. Stem cells 33, 1759–1770.

30 Lippmann, E. S., Williams, C. E., Ruhl, D. A., Estevez-Silva, M. C., Chapman, E. R., Coon, J. J., Ashton, R. S.(2015). Deterministic HOX patterning in human pluripotent stem cell-derived neuroectoderm. Stem cell reports 4, 632–644.

31 Gouti, M., Delile, J., Stamataki, D., Wymeersch, F. J., Huang, Y., Kleinjung, J., Wilson, V., Briscoe, J.(2017). A Gene Regulatory Network Balances Neural and Mesoderm Specification during Vertebrate Trunk Development. Developmental cell 41, 243–261.e7.

32 Faustino Martins, J. M., Fischer, C., Urzi, A., Vidal, R., Kunz, S., Ruffault, P. L., Kabuss, L., Hube, I., Gazzerro, E., Birchmeier, C., Spuler, S., Sauer, S., Gouti, M.(2020). Self-Organizing 3D Human Trunk Neuromuscular Organoids. Cell stem cell 26, 172–186.e6.

33 Yamanaka, Y., Hamidi, S., Yoshioka-Kobayashi, K., Munira, S., Sunadome, K., Zhang, Y., Kurokawa, Y., Ericsson, R., Mieda, A., Thompson, J. L., Kerwin, J., Lisgo, S., Yamamoto, T., Moris, N., Martinez-Arias, A., Tsujimura, T., Alev, C. (2023). Reconstituting human somitogenesis in vitro. Nature 614, 509–520.

34 Miao, Y., Djeffal, Y., De Simone, A., Zhu, K., Lee, J. G., Lu, Z., Silberfeld, A., Rao, J., Tarazona, O. A., Mongera, A., Rigoni, P., Diaz-Cuadros, M., Song, L. M. S., Di Talia, S., Pourquié, O. (2023). Reconstruction and deconstruction of human somitogenesis in vitro. Nature 614, 500–508.

35 Sanaki-Matsumiya, M., Matsuda, M., Gritti, N., Nakaki, F., Sharpe, J., Trivedi, V., Ebisuya, M. (2022). Periodic formation of epithelial somites from human pluripotent stem cells. Nat Commun 13, 2325.

36 Gribaudo, S., Robert, R., van Sambeek, B., Mirdass, C., Lyubimova, A., Bouhali, K., Ferent, J., Morin, X., van Oudenaarden, A., Nedelec, S. (2023). Self-organizing models of human trunk organogenesis recapitulate spinal cord and spine co-morphogenesis. Nat Biotechnol.

37 Rossi, G., Broguiere, N., Miyamoto, M., Boni, A., Guiet, R., Girgin, M., Kelly, R. G., Kwon, C., Lutolf, M. P.(2021). Capturing Cardiogenesis in Gastruloids. Cell stem cell 28, 230–240.e6.

38 Lewis-Israeli, Y. R., Wasserman, A. H., Gabalski, M. A., Volmert, B. D., Ming, Y., Ball, K. A., Yang, W., Zou, J., Ni, G., Pajares, N., Chatzistavrou, X., Li, W., Zhou, C., Aguirre, A. (2021). Self-assembling human heart organoids for the modeling of cardiac development and congenital heart disease. Nat Commun 12, 5142.

39 Hofbauer, P., Jahnel, S. M., Papai, N., Giesshammer, M., Deyett, A., Schmidt, C., Penc, M., Tavernini, K., Grdseloff, N., Meledeth, C., Ginistrelli, L. C., Ctortecka, C., Šalic, Š., Novatchkova, M., Mendjan, S (2021). Cardioids reveal self-organizing principles of human cardiogenesis. Cell, 184, 3299–3317.e22.

40 Drakhlis, L., Biswanath, S., Farr, C. M., Lupanow, V., Teske, J., Ritzenhoff, K., Franke, A., Manstein, F., Bolesani, E., Kempf, H., Liebscher, S., Schenke-Layland, K., Hegermann, J., Nolte, L., Meyer, H., de la Roche, J., Thiemann, S., Wahl-Schott, C., Martin, U., Zweigerdt, R. (2021). Human heart-forming organoids recapitulate early heart and foregut development. Nat Biotechnol 39, 737–746.

41 Frith, T. J., Granata, I., Wind, M., Stout, E., Thompson, O., Neumann, K., Stavish, D., Heath, P. R., Ortmann, D., Hackland, J. O., Anastassiadis, K., Gouti, M., Briscoe, J., Wilson, V., Johnson, S. L., Placzek, M., Guarracino, M. R., Andrews, P. W., Tsakiridis, A. (2018). Human axial progenitors generate trunk neural crest cells in vitro. eLife, 7, e35786.

42 Fan, Y., Hackland, J., Baggiolini, A., Hung, L. Y., Zhao, H., Zumbo, P., Oberst, P., Minotti, A. P., Hergenreder, E., Najjar, S., Huang, Z., Cruz, N. M., Zhong, A., Sidharta, M., Zhou, T., de Stanchina, E., Betel, D., White, R. M., et al. (2023). hPSC-derived sacral neural crest enables rescue in a severe model of Hirschsprung’s disease. Cell stem cell 30, 264–282.e9.

43 Urciuolo, A., Quarta, M., Morbidoni, V., Gattazzo, F., Molon, S., Grumati, P., Montemurro, F., Tedesco, F. S., Blaauw, B., Cossu, G., Vozzi, G., Rando, T. A., Bonaldo, P. (2013). Collagen VI regulates satellite cell self-renewal and muscle regeneration. Nat Commun 4, 1964.

44 Tierney, M. T., Gromova, A., Sesillo, F. B., Sala, D., Spenlé, C., Orend, G., Sacco, A. (2016). Autonomous Extracellular Matrix Remodeling Controls a Progressive Adaptation in Muscle Stem Cell Regenerative Capacity during Development. Cell reports 14, 1940–1952.

45 Warmflash, A., Sorre, B., Etoc, F., Siggia, E. D., Brivanlou, A. H. (2014). A method to recapitulate early embryonic spatial patterning in human embryonic stem cells. Nat Methods 11, 847–854.

46 Xiao, Z., Cui, L., Yuan, Y., He, N., Xie, X., Lin, S., Yang, X., Zhang, X., Shi, P., Wei, Z., Li, Y., Wang, H., Wang, X., Wei, Y., Guo, J., Yu, L. (2024). 3D reconstruction of a gastrulating human embryo. Cell, S0092-8674(24)00357–X.

47 Ikeya, M., Lee, S. M., Johnson, J. E., McMahon, A. P., Takada, S. (1997). Wnt signalling required for expansion of neural crest and CNS progenitors. Nature 389, 966–970.

48 Chal, J., Oginuma, M., Al Tanoury, Z., Gobert, B., Sumara, O., Hick, A., Bousson, F., Zidouni, Y., Mursch, C., Moncuquet, P., Tassy, O., Vincent, S., Miyanari, A., Bera, A., Garnier, J. M., Guevara, G., Hestin, M., Kennedy, L., Hayashi, S., Drayton, B., et al. (2015). Differentiation of pluripotent stem cells to muscle fiber to model Duchenne muscular dystrophy. Nat Biotechnol 33, 962–969.

49 Chen, Y. W., Huang, S. X., de Carvalho, A. L. R. T., Ho, S. H., Islam, M. N., Volpi, S., Notarangelo, L. D., Ciancanelli, M., Casanova, J. L., Bhattacharya, J., Liang, A. F., Palermo, L. M., Porotto, M., Moscona, A., Snoeck, H. W. (2017). A three-dimensional model of human lung development and disease from pluripotent stem cells. Nat Cell Biol 19, 542–549.

50 Keijzer, R., van Tuyl, M., Meijers, C., Post, M., Tibboel, D., Grosveld, F., Koutsourakis, M. (2001). The transcription factor GATA6 is essential for branching morphogenesis and epithelial cell differentiation during fetal pulmonary development. Development 128, 503–511.

51 Choi, S., Ferrari, G., Tedesco, F.S. (2020). Cellular dynamics of myogenic cell migration: molecular mechanisms and implications for skeletal muscle cell therapies. EMBO Mol Med. 12, e12357.

52 Xi, H., Langerman, J., Sabri, S., Chien, P., Young, C. S., Younesi, S., Hicks, M., Gonzalez, K., Fujiwara, W., Marzi, J., Liebscher, S., Spencer, M., Van Handel, B., Evseenko, D., Schenke-Layland, K., Plath, K., Pyle, A. D. (2020). A human skeletal muscle atlas identifies the trajectories of stem and progenitor cells across development and from human pluripotent stem cells. Cell Stem Cell, 27, 158–176 e10.

53 Biressi, S., Bjornson, C. R., Carlig, P. M., Nishijo, K., Keller, C., Rando, T. A. (2013). Myf5 expression during fetal myogenesis defines the developmental progenitors of adult satellite cells. Developmental biology 379, 195–207.

54 Sellung, D., Heil, L., Daya, N., Jacobsen, F., Mertens-Rill, J., Zhuge, H., Döring, K., Piran, M., Milting, H., Unger, A., Linke, W. A., Kley, R., Preusse, C., Roos, A., Fürst, D. O., Ven, P. F. M. V., Vorgerd, M.(2023). Novel Filamin C Myofibrillar Myopathy Variants Cause Different Pathomechanisms and Alterations in Protein Quality Systems. Cells 12, 1321.

55 Carlson, M. E., Conboy, M. J., Hsu, M., Barchas, L., Jeong, J., Agrawal, A., Mikels, A. J., Agrawal, S., Schaffer, D. V., Conboy, I. M. (2009). Relative roles of TGF-beta1 and Wnt in the systemic regulation and aging of satellite cell responses. Aging cell, 8, 676–689.

56 Carlson, M., Hsu, M. Conboy, I. (2008). Imbalance between pSmad3 and Notch induces CDK inhibitors in old muscle stem cells. Nature 454, 528–532.

57 Stantzou, A., Schirwis, E., Swist, S., Alonso-Martin, S., Polydorou, I., Zarrouki, F., Mouisel, E., Beley, C., Julien, A., Le Grand, F., Garcia, L., Colnot, C., Birchmeier, C., Braun, T., Schuelke, M., Relaix, F., Amthor, H. (2017). BMP signaling regulates satellite cell-dependent postnatal muscle growth. Development 144, 2737–2747.

58 Schaaf, G. J., van Gestel, T. J., Brusse, E., Verdijk, R. M., de Coo, I. F., van Doorn, P. A., van der Ploeg, A. T., Pijnappel, W. W. (2015). Lack of robust satellite cell activation and muscle regeneration during the progression of Pompe disease. Acta neuropathol commun 3, 65.

59. Schaaf, G. J., van Gestel, T. J. M., In’t Groen, S. L. M., de Jong, B., Boomaars, B., Tarallo, A., Cardone, M., Parenti, G., van der Ploeg, A. T., Pijnappel, W. W. M. P. (2018). Satellite cells maintain regenerative capacity but fail to repair disease-associated muscle damage in mice with Pompe disease. Acta neuropathol commun 6, 119.

60 Lagalice, L., Pichon, J., Gougeon, E., Soussi, S., Deniaud, J., Ledevin, M., Maurier, V., Leroux, I., Durand, S., Ciron, C., Franzoso, F., Dubreil, L., Larcher, T., Rouger, K., Colle, M. A. (2018). Satellite cells fail to contribute to muscle repair but are functional in Pompe disease (glycogenosis type II). Acta neuropathol commun 6, 116.

61 Biressi, S., Tagliafico, E., Lamorte, G., Monteverde, S., Tenedini, E., Roncaglia, E., Ferrari, S., Ferrari, S., Cusella-De Angelis, M. G., Tajbakhsh, S., Cossu, G. (2007). Intrinsic phenotypic diversity of embryonic and fetal myoblasts is revealed by genome-wide gene expression analysis on purified cells. Developmental biology 304, 633–651.

62 Dietrich, S., Abou-Rebyeh, F., Brohmann, H., Bladt, F., Sonnenberg-Riethmacher, E., Yamaai, T., Lumsden, A., Brand-Saberi, B., Birchmeier, C. (1999). The role of SF/HGF and c-Met in the development of skeletal muscle. Development 126, 1621–1629.

63 Relaix, F., Rocancourt, D., Mansouri, A., Buckingham, M. (2004). Divergent functions of murine Pax3 and Pax7 in limb muscle development. Genes & development, 18, 1088–1105.

64 Quarta, M., Brett, J. O., DiMarco, R., De Morree, A., Boutet, S. C., Chacon, R., Gibbons, M. C., Garcia, V. A., Su, J., Shrager, J. B., Heilshorn, S., Rando, T. A. (2016). An artificial niche preserves the quiescence of muscle stem cells and enhances their therapeutic efficacy. Nat Biotechnol 34, 752–759.

65 Xi, H., Fujiwara, W., Gonzalez, K., Jan, M., Liebscher, S., Van Handel, B., Schenke-Layland, K., Pyle, A. D. (2017). In Vivo Human Somitogenesis Guides Somite Development from hPSCs. Cell Rep. 18, 1573–1585.

66 Choi, I. Y., Lim, H., Estrellas, K., Mula, J., Cohen, T. V., Zhang, Y., Donnelly, C. J., Richard, J. P., Kim, Y. J., Kim, H., Kazuki, Y., Oshimura, M., Li, H. L., Hotta, A., Rothstein, J., Maragakis, N., Wagner, K. R., Lee, G.(2016). Concordant but varied phenotypes among Duchenne muscular dystrophy patient-specific myoblasts derived using a human iPSC-based model. Cell Rep. 15, 2301–2312.

67 Shelton, M., Metz, J., Liu, J., Carpenedo, R. L., Demers, S. P., Stanford, W. L., Skerjanc, I. S. (2014). Derivation and expansion of PAX7-positive muscle progenitors from human and mouse embryonic stem cells. Stem Cell Rep.orts 3, 516–529.

68 Borchin, B., Chen, J., Barberi, T. (2013). Derivation and FACS-mediated purification of PAX3+/PAX7+ skeletal muscle precursors from human pluripotent stem cells. Stem Cell Reports 1, 620–631.

69 Rao, L., Qian, Y., Khodabukus, A., Ribar, T.,Bursac, N. (2018). Engineering human pluripotent stem cells into a functional skeletal muscle tissue. Nat. Commun. 9, 126.

70 Maffioletti, S. M., Sarcar, S., Henderson, A. B. H., Mannhardt, I., Pinton, L., Moyle, L. A., Steele-Stallard, H., Cappellari, O., Wells, K. E., Ferrari, G., Mitchell, J. S., Tyzack, G. E., Kotiadis, V. N., Khedr, M., Ragazzi, M., Wang, W., Duchen, M. R., Patani, R., Zammit, P. S., Wells, D. J., et al. (2018). Three-Dimensional human iPSC-derived artificial skeletal muscles model muscular dystrophies and enable multilineage tissue engineering. Cell Rep. 23, 899–908.

71 Darabi, R., Arpke, R. W., Irion, S., Dimos, J. T., Grskovic, M., Kyba, M., Perlingeiro, R. C. (2012). Human ES- and iPS-derived myogenic progenitors restore DYSTROPHIN and improve contractility upon transplantation in dystrophic mice. Cell stem cell, 10, 610–619.

72 Sun, C., Kannan, S., Choi, I. Y., Lim, H., Zhang, H., Chen, G. S., Zhang, N., Park, S. H., Serra, C., Iyer, S. R., Lloyd, T. E., Kwon, C., Lovering, R. M., Lim, S. B., Andersen, P., Wagner, K. R., Lee, G.(2022). Human pluripotent stem cell-derived myogenic progenitors undergo maturation to quiescent satellite cells upon engraftment. Cell stem cell, 29, 610–619.e5.

## References (methods)

73 Dorn, I., Klich, K., Arauzo-Bravo, M. J., Radstaak, M., Santourlidis, S., Ghanjati, F., Radke, T. F., Psathaki, O. E., Hargus, G., Kramer, J., Einhaus, M., Kim, J. B., Kögler, G., Wernet, P., Schöler, H. R., Schlenke, P., Zaehres, H. (2015). Erythroid differentiation of human induced pluripotent stem cells is independent of donor cell type of origin. Haematologica 100, 32–41.

74 Daya, N. M., Mavrommatis, L., Zhuge, H., Athamneh, M., Roos, A., Gläser, D., Doering, K., Zaehres, H., Vorgerd, M., Güttsches, A. K. (2023). Generation of a human iPSC line (HIMRi001-A) from a patient with filaminopathy. Stem Cell Res. 72, 103210.

75 Daya, N. M., Döring, K., Zhuge, H., Volke, L., Stab, V., Dietz, J., Athamneh, M., Roos, A., Zaehres, H., Güttsches, A. K., Mavrommatis, L., Vorgerd, M. (2024). Generation of two hiPSCs lines of two patients carrying truncating mutations in the dimerization domain of filamin C. Stem Cell Res. 76, 103320.

76 Generation of two induced pluripotent stem cell lines (HIMRi006-A and HIMRi007-A) from Pompe patients with infantile and late disease onset. Stem Cell Res. Under Revision.

77 Mavrommatis, L., Jeong, H. W., Kindler, U., Gomez-Giro, G., Kienitz, M. C., Stehling, M., Psathaki, O. E., Zeuschner, D., Bixel, M. G., Han, D., Morosan-Puopolo, G., Gerovska, D., Yang, J. H., Kim, J. B., Arauzo-Bravo, M. J., Schwamborn, J. C., Hahn, S. A., Adams, R. H., Schöler, H. R., Vorgerd, M., Brand-Saberi B., Zaehres, H. (2023). Human skeletal muscle organoids model fetal myogenesis and sustain uncommitted PAX7 myogenic progenitors. Elife. 12, RP87081.

78 Alexander, M. S., Rozkalne, A., Colletta, A., Spinazzola, J. M., Johnson, S., Rahimov, F., Meng, H., Lawlor, M. W., Estrella, E., Kunkel, L. M., Gussoni, E. (2016). CD82 Is a Marker for Prospective Isolation of Human Muscle Satellite Cells and Is Linked to Muscular Dystrophies. Cell Stem Cell 19, 800–807.

79 Sherwood, R. I., Christensen, J. L., Conboy, I. M., Conboy, M. J., Rando, T. A., Weissman, I. L., Wagers, A. J. (2004). Isolation of adult mouse myogenic progenitors: functional heterogeneity of cells within and engrafting skeletal muscle. Cell, 119, 543–554.

80 Satija, R., Farrell, J. A., Gennert, D., Schier, A. F., Regev, A.(2015). Spatial reconstruction of single-cell gene expression data. Nat. Biotechnol. 33, 495–502.

81 Butler, A., Hoffman, P., Smibert, P., Papalexi, E., Satija, R. (2018). Integrating single-cell transcriptomic data across different conditions, technologies, and species. Nat. Biotechnol. 36, 411–420.

82 Stuart, T., Butler, A., Hoffman, P., Hafemeister, C., Papalexi, E., Mauck, W. M., 3rd, Hao, Y., Stoeckius, M., Smibert, P., Satija, R. (2019). Comprehensive Integration of Single-Cell Data. Cell 177, 1888–1902.e21.

83 Hao, Y., Hao, S., Andersen-Nissen, E., Mauck, W. M., 3rd, Zheng, S., Butler, A., Lee, M. J., Wilk, A. J., Darby, C., Zager, M., Hoffman, P., Stoeckius, M., Papalexi, E., Mimitou, E. P., Jain, J., Srivastava, A., Stuart, T., Fleming, L. M., Yeung, B., Rogers, A. J., et al. (2021). Integrated analysis of multimodal single-cell data. Cell 184, 3573–3587.e29.

84 Hao, Y., Stuart, T., Kowalski, M. H., Choudhary, S., Hoffman, P., Hartman, A., Srivastava, A., Molla, G., Madad, S., Fernandez-Granda, C., Satija, R.(2024). Dictionary learning for integrative, multimodal and scalable single-cell analysis. Nat Biotechnol 42, 293–304.

85 Wolf, F., Angerer, P., Theis, F. (2018). SCANPY: large-scale single-cell gene expression data analysis. Genome Biol 19, 15.

86 van Dijk, D., Sharma, R., Nainys, J., Yim, K., Kathail, P., Carr, A. J., Burdziak, C., Moon, K. R., Chaffer, C. L., Pattabiraman, D., Bierie, B., Mazutis, L., Wolf, G., Krishnaswamy, S., Pe’er, D. (2018). Recovering Gene Interactions from Single-Cell Data Using Data Diffusion. Cell 174, 716–729.

87 Tirosh, I., Izar, B., Prakadan, S. M., Wadsworth, M. H., 2nd, Treacy, D., Trombetta, J. J., Rotem, A., Rodman, C., Lian, C., Murphy, G., Fallahi-Sichani, M., Dutton-Regester, K., Lin, J. R., Cohen, O., Shah, P., Lu, D., Genshaft, A. S., Hughes, T. K., Ziegler, C. G., Kazer, S. W., et al. (2016). Dissecting the multicellular ecosystem of metastatic melanoma by single-cell RNA-seq. Science 352, 189–196.

88 van den Brink, S. C., Sage, F., Vértesy, Á., Spanjaard, B., Peterson-Maduro, J., Baron, C. S., Robin, C., van Oudenaarden, A. (2017). Single-cell sequencing reveals dissociation-induced gene expression in tissue subpopulations. Nat. Methods 14, 935–936.

89 Kiselev, V. Y., Kirschner, K., Schaub, M. T., Andrews, T., Yiu, A., Chandra, T., Natarajan, K. N., Reik, W., Barahona, M., Green, A. R., Hemberg, M. (2017). SC3: consensus clustering of single-cell RNA-seq data. Nat Methods 14, 483–486.

90 Vahid, M. R., Brown, E. L., Steen, C. B., Zhang, W., Jeon, H. S., Kang, M., Gentles, A. J., Newman, A. M. (2023). High-resolution alignment of single-cell and spatial transcriptomes with CytoSPACE. Nat Biotechnol 41, 1543–1548.

91 Aibar, S., González-Blas, C. B., Moerman, T., Huynh-Thu, V. A., Imrichova, H., Hulselmans, G., Rambow, F., Marine, J. C., Geurts, P., Aerts, J., van den Oord, J., Atak, Z. K., Wouters, J., Aerts, S., et al. (2017). SCENIC: single-cell regulatory network inference and clustering. Nat. Methods 14,1083–1086.

92 Jacomy, M., Venturini, T., Heymann, S., Bastian, M. (2014). ForceAtlas2, a continuous graph layout algorithm for handy network visualization designed for the Gephi software. PLoS One 9, e98679.

93 Wolf, F.A. et al. (2019). PAGA: graph abstraction reconciles clustering with trajectory inference through a topology preserving map of single cells. Genome Biol. 20, 59.

94 Angerer, P., Haghverdi, L., Büttner, M., Theis, F. J., Marr, C., Buettner, F. (2016) destiny: diffusion maps for large-scale single-cell data in R. Bioinformatics 32, 1241–1243.

95 Setty, M., Kiseliovas, V., Levine, J., Gayoso, A., Mazutis, L., Pe’er, D. (2019). Characterization of cell fate probabilities in single-cell data with Palantir. Nat Biotechnol 37, 451–460.

96 Faure, L., Soldatov, R., Kharchenko, P.V., Adameyko I. (2022). scFates: a scalable python package for advanced pseudotime and bifurcation analysis from single cell data. Bioinformatics, btac746.

97 Albergante, L., Mirkes, E., Bac, J., Chen, H., Martin, A., Faure, L., Barillot, E., Pinello, L., Gorban, A., Zinovyev, A. (2020). Robust and Scalable Learning of Complex Intrinsic Dataset Geometry via ElPiGraph. Entropy 22, 296.

98 Finak, G., McDavid, A., Yajima, M., Deng, J., Gersuk, V., Shalek, A. K., Slichter, C. K., Miller, H. W., McElrath, M. J., Prlic, M., Linsley, P. S., Gottardo, R. (2015). MAST: a flexible statistical framework for assessing transcriptional changes and characterizing heterogeneity in single-cell RNA sequencing data. Genome biology 16, 278.

99 Lotfollahi, M., Naghipourfar, M., Luecken, M. D., Khajavi, M., Büttner, M., Wagenstetter, M., Avsec, Ž., Gayoso, A., Yosef, N., Interlandi, M., Rybakov, S., Misharin, A. V., Theis, F. J. (2022). Mapping single-cell data to reference atlases by transfer learning. Nat. Biotechnol. 40, 121–130.

100 Xu, C., Lopez, R., Mehlman, E., Regier, J., Jordan, M. I., Yosef, N. (2021). Probabilistic harmonization and annotation of single-cell transcriptomics data with deep generative models. Molecular systems biology 17, e9620.

101 La Manno, G., Soldatov, R., Zeisel, A., Braun, E., Hochgerner, H., Petukhov, V., Lidschreiber, K., Kastriti, M. E., Lönnerberg, P., Furlan, A., Fan, J., Borm, L. E., Liu, Z., van Bruggen, D., Guo, J., He, X., Barker, R., Sundström, E., Castelo-Branco, G., Cramer, P., et al. (2018). RNA velocity of single cells. Nature 560, 494–498.

102 Bergen, V., Lange, M., Peidli, S., Wolf, F. A., Theis, F. J.(2020). Generalizing RNA velocity to transient cell states through dynamical modeling. Nat. Biotechnol. 38, 1408–1414.

103 Jin, S., Guerrero-Juarez, C. F., Zhang, L., Chang, I., Ramos, R., Kuan, C. H., Myung, P., Plikus, M. V., Nie, Q. (2021). Inference and analysis of cell-cell communication using CellChat. Nat. Commun. 12, 1088.

104 Browaeys, R., Saelens, W., Saeys, Y. (2020). NicheNet: modeling intercellular communication by linking ligands to target genes. Nat Methods 17, 159–162.

105 Zhou, Y., Zhou, B., Pache, L., Chang, M., Khodabakhshi, A. H., Tanaseichuk, O., Benner, C., Chanda, S. K (2019). Metascape provides a biologist-oriented resource for the analysis of systems-level datasets. Nat Commun 10, 1523.

106 Jassal, B., Matthews, L., Viteri, G., Gong, C., Lorente, P., Fabregat, A., Sidiropoulos, K., Cook, J., Gillespie, M., Haw, R., Loney, F., May, B., Milacic, M., Rothfels, K., Sevilla, C., Shamovsky, V., Shorser, S., Varusai, T., Weiser, J., Wu, G., et al. (2020). The reactome pathway knowledgebase. Nucleic Acids Res. 48, D1498–D503.

107 Soldatov, R., Kaucka, M., Kastriti, M. E., Petersen, J., Chontorotzea, T., Englmaier, L., Akkuratova, N., Yang, Y., Häring, M., Dyachuk, V., Bock, C., Farlik, M., Piacentino, M. L., Boismoreau, F., Hilscher, M. M., Yokota, C., Qian, X., Nilsson, M., Bronner, M. E., Croci, L., et al. (2019). Spatiotemporal structure of cell fate decisions in murine neural crest. Science 364, eaas9536.

108 He, P., Williams, B. A., Trout, D., Marinov, G. K., Amrhein, H., Berghella, L., Goh, S. T., Plajzer-Frick, I., Afzal, V., Pennacchio, L. A., Dickel, D. E., Visel, A., Ren, B., Hardison, R. C., Zhang, Y., Wold, B. J. (2020). The changing mouse embryo transcriptome at whole tissue and single-cell resolution. Nature 583, 760–767.

109 De Micheli, A. J., Laurilliard, E. J., Heinke, C. L., Ravichandran, H., Fraczek, P., Soueid-Baumgarten, S., De Vlaminck, I., Elemento, O., Cosgrove, B. D. (2020). Single-cell analysis of the muscle stem cell hierarchy identifies heterotypic communication signals involved in skeletal muscle regeneration. Cell Rep. 30, 3583–3595 e5.

110 Zeng, B., Liu, Z., Lu, Y., Zhong, S., Qin, S., Huang, L., Zeng, Y., Li, Z., Dong, H., Shi, Y., Yang, J., Dai, Y., Ma, Q., Sun, L., Bian, L., Han, D., Chen, Y., Qiu, X., Wang, W., Marín, O., et al. (2023). The single-cell and spatial transcriptional landscape of human gastrulation and early brain development. Cell Stem Cell 30, 851–866.e7.

111 Xu, Y., Zhang, T., Zhou, Q., Hu, M., Qi, Y., Xue, Y., Nie, Y., Wang, L., Bao, Z., Shi, W. (2023). A single-cell transcriptome atlas profiles early organogenesis in human embryos. Nat Cell Biol 25, 604–615.

112 Perez, K., Ciotlos, S., McGirr, J., Limbad, C., Doi, R., Nederveen, J. P., Nilsson, M. I., Winer, D. A., Evans, W., Tarnopolsky, M., Campisi, J., Melov, S. (2022). Single nuclei profiling identifies cell specific markers of skeletal muscle aging, frailty, and senescence. Aging 14, 9393–9422.

113 Zhang, B., He, P., Lawrence, J. E. G., Wang, S., Tuck, E., Williams, B. A., Roberts, K., Kleshchevnikov, V., Mamanova, L., Bolt, L., Polanski, K., Li, T., Elmentaite, R., Fasouli, E. S., Prete, M., He, X., Yayon, N., Fu, Y., Yang, H., Liang, C., et al. (2023). A human embryonic limb cell atlas resolved in space and time. Nature.

